# Striatal cell-type-specific encoding stability and reorganization for goal-directed behavior and habit

**DOI:** 10.1101/2025.01.26.634924

**Authors:** Melissa Malvaez, Alvina Liang, Baila S. Hall, Jacqueline R. Giovanniello, Natalie Paredes, Julia Y. Gonzalez, Garrett J. Blair, Ana C. Sias, Michael D. Murphy, Wanyi Guo, Alicia Wang, Malika Singh, Nicholas K. Griffin, Samuel P. Bridges, Anna Wiener, Jenna S. Pimenta, Sandra M. Holley, Carlos Cepeda, Michael S. Levine, Peter D. Balsam, H. Tad Blair, Andrew M. Wikenheiser, Kate M. Wassum

**Affiliations:** Dept. of Psychology, UCLA, Los Angeles, CA 90095; Brain Research Institute, UCLA, Los Angeles, CA 90095, USA; Intellectual and Developmental Disabilities Research Center, Semel Institute for Neuroscience and Human Behavior, David Geffen School of Medicine, University of California, Los Angeles, Los Angeles, United States; Department of Psychology, Barnard College, New York City, NY, USA; Columbia University and New York State Psychiatric Institute, New York City, NY, USA

**Keywords:** learning, decision making, instrumental conditioning, reward, striatum, habit, devaluation, calcium imaging, chemogenetics, direct and indirect pathway

## Abstract

Goal-directed decision making relies on learned action-outcome associations to allow actions to adapt when outcomes change. With repetition, behavioral control shifts toward habit, enabling more automatic action execution at the expense of flexibility. Much is unknown about how the brain learns and supports goal-directed behavior and what changes as habits form. Here, we longitudinally tracked and manipulated the activity of two major dorsomedial striatum populations, D1^+^ and D2/A2A^+^ neurons, to reveal their activity and function during goal-directed learning and habit formation. Across learning, DMS D1^+^ neurons stably encode instrumental actions, carry information about behavioral engagement and organization, and develop and maintain action-reward representational similarity and conjunctive activation. This persists even after behavioral control transitions to habit. In contrast, one ensemble of DMS A2A^+^ neurons transiently encodes actions during early learning and another slowly starts to encode actions as habits form. Both DMS D1^+^ and A2A^+^ neuronal activity mediates the learning processes underlying goal-directed control, but only DMS D1^+^ neurons maintain support of adaptive, goal-directed decision making. These data reveal a cell-type-specific dissociation between stable v. reorganizing striatal action representations and, independently, persistent v. transient contributions to goal-directed control.

---

When making a decision, we often use our knowledge of available actions and their consequences to prospectively evaluate our options and choose actions that will cause beneficial outcomes^1–3^. Learned action-outcome associations support the predictions and inferences needed for adaptive, goal-directed decision making^2–7^. Should an outcome become unbeneficial, we will reduce the actions that cause it. For well-practiced routines we have a more efficient strategy: habits. With repeated successful practice, behavioral control transitions from reliance on action-outcome knowledge to more automatic execution^2,8–11^. Habits are, therefore, resource efficient, but inflexible. We might continue a habit even when it does not serve our goals. Balance between goal-directed and habitual control allows behavior to be adaptive when needed, yet efficient when appropriate^12,13^. Disrupted goal-directed behavior and overreliance on habit can, however, cause inadequate consideration of consequences, disrupted decision making, inflexible behavior, and a lower threshold for compulsivity^14–17^. This can contribute to cognitive deficits seen in addiction^18–29^ and myriad other mental health conditions^16,25,30–39^. Yet, despite importance for understanding adaptive and maladaptive behavior, much remains unknown about how the brain learns and supports goal-directed behavior and what changes as habits form.

The dorsomedial striatum (DMS) is an evolutionarily conserved hub for goal-directed behavior^8,40–52^. In the striatum, experiential factors that drive learning converge onto basal ganglia outputs that drive motor execution^53^. There are two major subtypes of these outputs: one expressing dopamine D1 receptors and another expressing dopamine D2 and adenosine Adora2A (A2A) receptors. Whereas D1^+^ neurons are more dominant in the direct pathway to basal ganglia output nuclei, D2/A2A^+^ neurons are associated with the multisynaptic, indirect output pathway^54–56^, though this organization is likely more complex than originally thought^57,58^. The activity of these neurons has been well characterized during locomotion and previously learned actions^8,40,59^, but little is known of how their activity is shaped during action-outcome learning or transitions in behavioral control. Such information is needed to understand how learning and repeated practice of an action shape striatal activity to support the learning underlying goal-directed control, adaptive decision making, and the transition to habit. Here we addressed this gap in knowledge by longitudinally tracking and manipulating the activity of DMS D1^+^ and A2A^+^ neurons during action-outcome learning and habit formation. Three questions guided our investigations: 1. To what extent do DMS D1^+^ and A2A^+^ neurons carry information about instrumental behavior and earned rewards? 2. How stable are these activity patterns as behavior transitions from goal-directed to habitual? 3. Do DMS D1^+^ and A2A^+^ neurons support the learning processes underlying goal-directed control and goal-directed decision making?

## RESULTS

### The DMS D1^+^ neuronal population conveys information about instrumental behavior and develops action-reward representational similarity across training

We first asked how DMS D1^+^ neuronal activity relates to instrumental behavior during action-outcome learning and the transition to habit. We used UCLA miniscopes^60^ with a gradient refractive index (GRIN) lens for longitudinal, cellular-resolution, microendoscopic imaging of the genetically encoded calcium indicator jGCaMP7s^61^ selectively expressed in DMS D1^+^ neurons of Drd1a-cre mice (Figure 1a-d). We recorded calcium activity as mice learned to lever press to earn food-pellet rewards in a self-initiated instrumental task (Figure 1e). We used a random-interval (RI) reinforcement schedule that with 4 training sessions enables goal-directed control and with overtraining (8 sessions) promotes habit formation (Extended Data Figure 1-1). All mice acquired the instrumental behavior (Figure 1f; see Extended Data Figure 1-2 for food-port entry data). To evaluate the extent of goal-directed v. habitual control, we used the outcome-specific devaluation test^2,62^. Mice were given 90-min, non-contingent access to the food pellet earned during training to induce a sensory-specific satiety rendering that pellet temporarily devalued. Lever pressing was then assessed in a 5-min, non-reinforced devaluation probe test. Performance was compared to that following satiation on an alternate food pellet to control for general satiety (Valued state). Goal-directed decisions involve the use of learned action-outcome associations to adaptively reduce action performance when the earned outcome is not valuable^1–3^. Thus, we operationally defined action-outcome-based, goal-directed behavior as sensitivity to outcome devaluation. Goal-directed behavior dominates after 4 training sessions, such that mice reduce lever-press actions when their outcome is devalued (Extended Data Figure 1-1). Habits are operationally defined by insensitivity to devaluation^2,8–11^. We observed this in mice following overtraining (Figure 1g-h; see also Extended Data Figure 1-1). To capture activity related to goal-directed behavior, we analyzed calcium activity at the beginning (session 1, RI-15-s training) and middle (session 4, RI-30s) of instrumental training. To evaluate activity related to habit, we analyzed calcium activity at the end of overtraining (session 8, RI-30s; see Supplemental Table 1 for neurons/mouse/session). We leveraged longitudinal tracking (Figure 1d; Extended Data Figure 1-3; Supplemental Table 1) to ask how individual DMS D1^+^ neurons change their activity across training phases.

**Figure 1:**
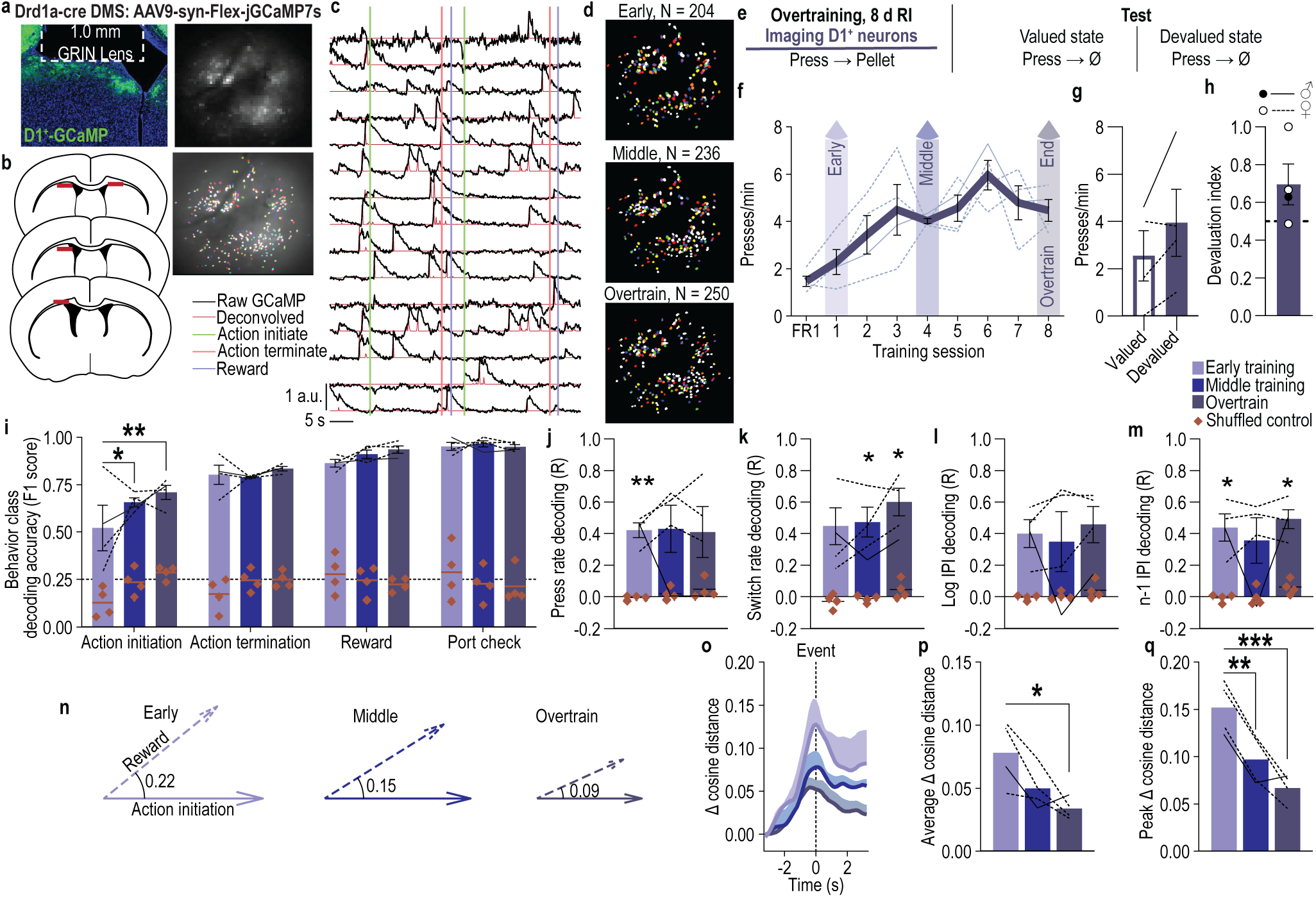
The DMS D1^+^ neuronal population encodes instrumental actions, reward outcomes, behavioral engagement and organizational information, and develops and maintains action-reward representational similarity across training. (a) Representative cre-dependent jGCaMP7s expression in DMS D1^+^ neurons (top left) and maximum intensity projection of the field-of-view with (top right) and without (bottom right) demarcation of cell ROIs for one session. (b) Map of DMS GRIN lens placements. (c) Representative raw fluorescence (*F* normalized to max, arbitrary units) and deconvolved calcium signal. (d) Representative recorded D1^+^ neuron spatial footprints during the first (early), 4^th^ (middle), and 8^th^ (overtrain) random-interval training sessions. Colored, coregistered neurons; white, non-coregistered neurons. (e) Procedure. RI, random-interval reinforcement schedule. (f) Training press rate. 1-way ANOVA, Training: F_1.81, 5.43_ = 4.84, *P* = 0.06. (g) Test press rate. 2-tailed t-test, t_3_ = 1.98, *P* = 0.14, 95% confidence interval (CI) –0.85 –3.65. (h) Devaluation index [(Devalued presses)/(Valued presses + Devalued presses)]. One-tailed Bayes factor, BF_10_ = 0.18. (i) Behavior class decoding accuracy from D1^+^ coregistered neuron activity in 0.4-s period prior to event compared to shuffled control. Line = chance. 3-way ANOVA, Neuron activity (v. shuffled): F_1, 3_ = 1092.83, *P* < 0.001. (j-m) Linear decoding accuracy from D1^+^ coregistered neuron activity of press rate (j; 2-way ANOVA, Neuron activity: F_1, 3_ = 12.16, *P* = 0.04), switch rate (k; 2-way ANOVA, Neuron activity: F_1, 3_ = 33.57, *P* = 0.01), log-transformed current interpress interval (l; 2-way ANOVA, Neuron activity: F_1, 3_ = 8.91, *P* = 0.05), and the prior press interpress interval (m; 2-way ANOVA, Neuron activity: F_1, 3_ = 22.10, *P* = 0.02). R = correlation between observed and decoded values. (n) Visualization of representative single-subject, single-timepoint example population vectors with cosine distance between action-initiation and earned-reward neuronal population vectors. (o-q) Change (Δ) in cosine angular distance, relative to pre-event baseline, between action-initiation and earned-reward D1^+^ neuronal population vectors across time and training (o), average (p; 1-way ANOVA, Training: F_1.60, 4.79_ = 8.46, *P* = 0.03), and maximum (q; peak; 1-way ANOVA, Training: F_1.02, 3.07_ = 29.20, *P* = 0.01) ± 2.5 s peri-event. D1-cre: *N* = 4 (1 male). Data presented as mean ± s.e.m. Males = closed circles/solid lines, Females = open circles/dashed lines. \**P* < 0.05, \*\**P* < 0.01, \*\*\**P*< 0.001. Full statistical reporting provided in Supplemental Table 2.

#### DMS D1^+^ neuron population activity conveys information about actions, outcomes, and behavioral engagement and organization

To establish the behavioral and task information available in DMS D1^+^ neuronal population activity, we trained a multi-class support vector machine (SVM) to classify behavioral events from the temporal fluorescence traces of all coregistered D1^+^ neurons. Behavioral events were *action initiation*: first lever press in the session, after earned reward collection, or after non-reinforced food-port check; *action termination*: last press before reward collection or non-reinforced food-port check; *check*: non-reinforced food-port entry; and *reward:* earned reward consumption. Across training, the decoder classified each behavior more accurately than one trained on shuffled data (Figure 1i). Decoding of action initiation improved during initial training, but did not improve further with overtraining. Thus, the D1^+^ neuronal population can convey information about actions and outcomes.

We next asked whether DMS D1^+^ neuronal population activity reflects features of behavioral engagement and organization associated with subsequent behavioral control. Using a linear regression model to predict sensitivity to outcome devaluation from training behavior, we found that 2 measures of behavioral engagement: press rate (lever presses/min) and behavior switch rate (rate of switching from pressing to checking the food port or vice versa) and 2 measures of behavioral organization: interpress interval variability/consistency (standard deviation of the log-transformed interpress intervals) and the extent to which current trial behavior relates to past trial behavior (correlation between current interpress interval, n, and interpress interval of the prior press, n-1^63,64^) jointly accounted for variation in subsequent sensitivity to devaluation (Extended Data Figure 1-4). Devaluation-sensitive, goal-directed behavior was associated with actively and flexibly engaging with pressing and checking for the reward and re-engagement in pressing (high press and switch rate) and behavior that is related to past performance (low interpress interval variability and high n:n-1 interpress interval correlation). Devaluation insensitivity was predicted by an absence of such press and switch rate coordination and behavior that was less related to past behavior (high interpress interval variability and low n:n-1 interpress interval correlation). Press rate, switch rate, log interpress interval, and past trial interpress interval could all be decoded across training beyond the temporally structured null model with linear regression models trained to decode these behavioral engagement and organizational variables from D1^+^ neuronal population activity (Figure 1j-m; see Extended Data Figure 1-5 for evidence that null model preserves internal temporal structure, accuracy of decoding as a function of time lag, and example decoding). Thus, DMS D1^+^ neuronal population activity contains information about local fluctuations in behavioral engagement and organization, variables that correlate with the extent of goal-directed v. habitual behavioral control.

#### The DMS D1^+^ neuronal population develops and maintains action-reward representational similarity with training

Having established that the D1^+^ neuronal population contains information about both instrumental actions and earned rewards, we next asked whether learning shapes the similarity of these representations. We found emergent learning-related convergence of the population representation of action and earned reward. We used a population vector analysis^65^ to examine the geometric relationship between the D1^+^ neuronal population representations of action initiation and earned reward during each phase of training (Figure 1n-q; see also Extended Data Figure 1-6). We defined the activity of each neuron as a dimension and the population response to each event as a vector in high-dimensional state space and used the cosine distance between the two vectors as a measure of the similarity in the pattern of population activity (Figure 1n). To control for any non-event-related change in population geometry, we subtracted the baseline cosine distance between the two vectors. With learning, the DMS D1⁺ neuronal population vectors representing action and earned reward outcome moved closer together in representational space. The distance between the action initiation and reward population vectors decreased with training (Figure 1o-p; see also Extended Data Figure 1-7 for concordant representational similarity analysis). Owing to the RI reinforcement schedule, this did not result from increased temporal predictability of reward. Rather, the time between action initiation and reward became slightly longer across training (Early action initiation – reward time: 18.05, s.e.m. = 4.09; Middle: 25.87, 7.94; Overtrain: 34.11, 7.94; 1-way ANOVA, Training: F_1.04, 3.12_ = 1.32, *P* = 0.33) and was similarly variable (Early action initiation – reward time average stdev = 27.81, s.e.m. = 11.46; Middle: 33.58, 9.27; Overtrain: 39.72, 6.99; 1-way ANOVA, Training: F_1.31, 3.93_ = 0.41, *P* = 0.61). Increased representational similarity did not occur for similar behavioral combinations in which the port-check was not reinforced or other behavior combinations, providing evidence against a global change in population geometry (Extended Data Figure 1-7). The action-reward representational similarity persisted after overtraining, suggesting habit formation does not disrupt DMS D1^+^ action-reward population geometry. Thus, the DMS D1^+^ neuronal population develops and maintains action-reward representational similarity.

### DMS D1^+^ neurons stably encode action initiation and develop action-reward conjunctive activation

#### DMS D1^+^ neurons are active around actions and earned rewards

We next examined how individual D1^+^ neurons encode task events during action-outcome learning and habit formation. A substantial proportion of D1^+^ neurons were significantly excited around, largely prior to, action initiation during each phase of training (Figure 2a-b; see Extended Data Figure 2-1 for encoding of neurons from the whole population, see Extended Data Figure 2-2a-c for example classification). Another ensemble was inhibited by action initiation (Figure 2c; Extended Data Figure 2-3). Subpopulations of D1^+^ neurons also encoded action termination (Extended Data Figure 2-4) and earned reward (Extended Data Figure 2-5). Thus, individual DMS D1^+^ neurons encode instrumental actions and reward outcomes during learning and habit formation.

**Figure 2:**
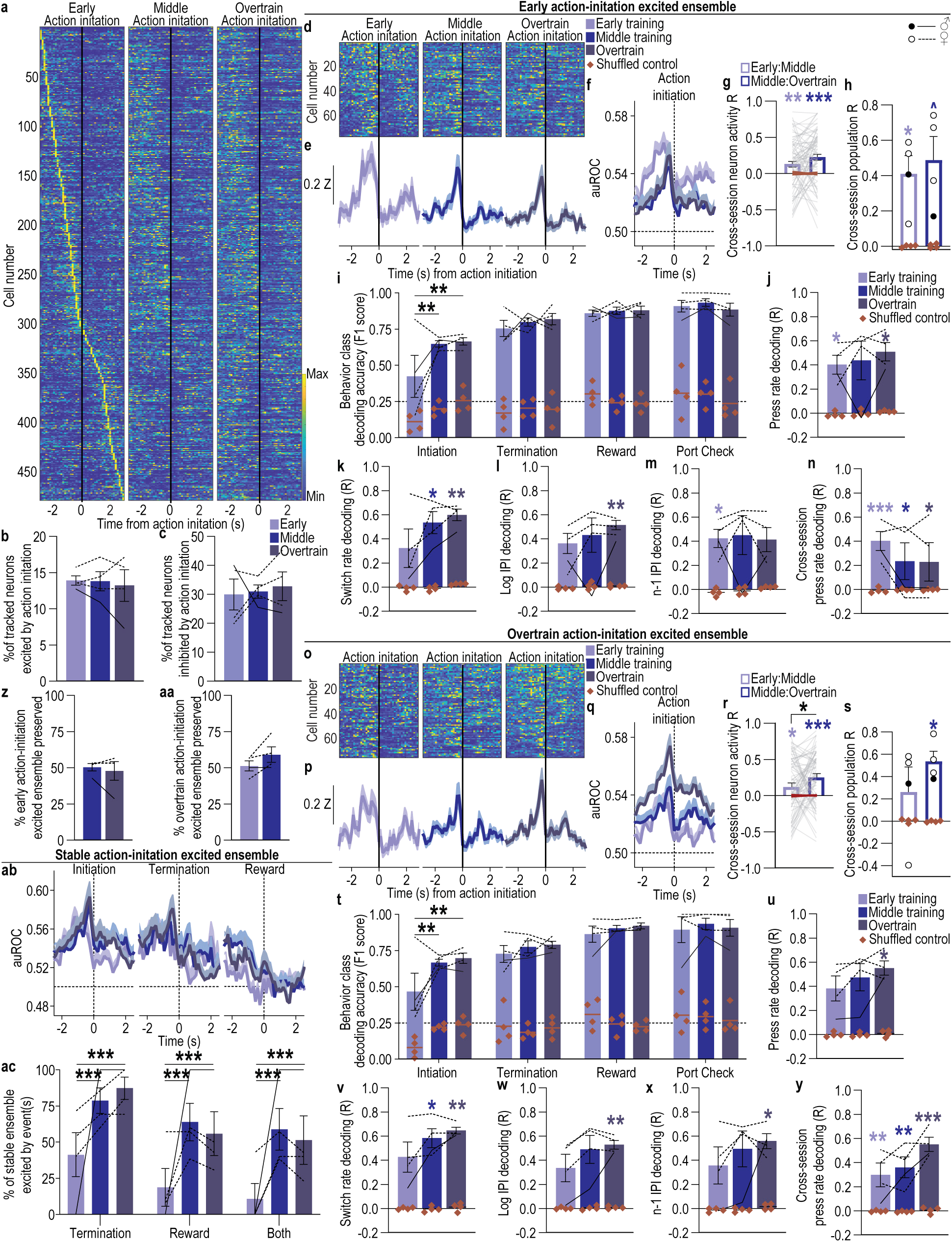
A subpopulation of DMS D1^+^ neurons stably encodes action initiation, contains behavioral engagement and organizational information, and develops conjunctive activation by earned outcome. **(a)** Heat map of minimum to maximum deconvolved activity (sorted by total activity) of each coregistered DMS D1^+^ neuron around lever-press action initiation. **(b-c)** Percent of coregistered neurons (Average 137 coregistered neurons/mouse, s.e.m. 37.73) significantly excited (b; 1-way ANOVA, F_1.13, 3.63_ = 0.14, *P* = 0.76), or inhibited (c; 1-way ANOVA, F_1.29, 3.86_ = 0.14, *P* = 0.79) around action initiation. **(d-f)** Activity and modulation across training of D1^+^ early action-initiation excited neurons. Heat map (d), Z-scored activity (e), and area under the receiver operating characteristic curve (auROC) modulation index (f) of early action-initiation excited neurons around action initiation across training. **(g-h)** Cross-session correlation of the activity around action initiation of each early action-initiation excited neuron (g; 2-way ANCOVA, Neuron activity: F_1, 75_ = 22.92, *P* < 0.001) or the population activity of these neurons (h; 2-way ANOVA, Neuron activity: F_1, 3_ = 39.10, *P* = 0.008). **(i)** Behavior class decoding accuracy from D1^+^ early action-initiation excited neuron activity in 0.4-s period prior to event compared to shuffled control. Line = chance. 3-way ANOVA, Neuron activity (v. shuffled): F_1, 3_ = 1118.27, *P* < 0.001. **(j-m)** Linear decoding accuracy from D1^+^ early action-initiation excited neuron activity of press rate (j; 2-way ANOVA, Neuron activity: F_1, 3_ = 21.00, *P* = 0.02), switch rate (k; 2-way ANOVA, Neuron activity: F_1, 3_ = 30.27, *P* = 0.01), log-transformed current interpress interval (l, 2-way ANOVA, Neuron activity: F_1, 3_ = 24.74, *P* = 0.01), and the prior press interpress interval (m; 2-way ANOVA, Neuron activity: F_1, 3_ = 21.31, *P* = 0.02). R = correlation between observed and decoded values. **(n)** Cross-session decoding accuracy of press rate from the activity of D1^+^ early action-initiation-excited neuron population activity on the 1^st^ training session. Planned, Bonferroni corrected, 2-tailed t-tests, Early: t_6_ = 7.77, *P* = 0.0007, 95% CI 0.24 – 0.59; Middle: t_6_ = 4.47, *P* = 0.01, 95% CI 0.06 – 0.41; Overtrain: t_6_ = 4.37, *P* = 0.01, 95% CI 0.06 – 0.41. **(o-q)** Activity and modulation across training of D1^+^ overtrain action-initiation excited neurons. Heat map (o), Z-scored activity (p), and auROC modulation index (q) of overtrain action-initiation excited neurons around action initiation across training. **(r-s)** Cross-session correlation of the activity around action initiation of each overtrain action-initiation excited neuron (r; 2-way ANCOVA, Neuron activity: F_1, 74_ = 21.06, *P* < 0.001) or the population activity of these neurons (s; 2-way ANOVA, Neuron activity: F_1, 3_ = 8.25, *P* = 0.06). **(t)** Behavior class decoding accuracy from D1^+^ overtrain action-initiation excited neuron activity. 3-way ANOVA, Neuron activity: F_1, 3_ = 262.75, *P* < 0.001. **(u-x)** Linear decoding accuracy from D1^+^ overtrain action-initiation excited neuron activity of press rate (u; 2-way ANOVA, Neuron activity: F_1, 3_ = 21.00, *P* = 0.02), switch rate (v; 2-way ANOVA, Neuron activity: F_1, 3_ = 52.93, *P* = 0.005), log-transformed current interpress interval (w; 2-way ANOVA, Neuron activity: F_1, 3_ = 24.70, *P* = 0.02), and the prior press interpress interval (x; 2-way ANOVA, Neuron activity: F_1, 3_ = 20.44, *P* = 0.02). **(y)** Cross-session decoding accuracy of lever-press rate from the activity of D1^+^ overtrain action-initiation-excited neuron population activity on the 8^th^ training session. Planned, Bonferroni corrected, 2-tailed t-tests, Early: t_6_ = 5.70, *P* = 0.004, 95% CI 0.13 to 0.48; Middle: t_6_ = 6.95, *P* = 0.001, 95% CI 0.19 – 0.54; Overtrain: t_6_ = 10.33, *P* = 0.0001, 95% CI 0.38 – 0.72. **(z)** Percent of D1^+^ early action-initiation excited neurons that continued to be significantly excited by action initiation on the 4^th^ and 8^th^ training sessions. 2-tailed Wilcoxon signed rank test, W = –1.00, P > 0.99. **(aa)** Percent of D1^+^ overtrain action-initiation excited neurons that were also significantly excited by action initiation on 1^st^ and 4^th^ training sessions. 2-tailed t-test, t_3_ = 2.02, *P* = 0.14, 95% CI –4.52 – 20.15. **(ab)** Modulation around action initiation, termination, and earned reward across training of D1^+^ stable action-initiation excited neurons. **(ac)** Percent of coregistered D1^+^ neurons stably excited by action initiation across training that were also significantly modulated by action termination and reward collection. 2-way ANOVA, Event: F_2, 6_ = 7.81, *P* = 0.02; Training: F_2, 6_ = 4.37, *P* = 0.07; Event x Training: F_1.86, 5.59_ = 1.76, *P* = 0.20. D1-cre: *N* = 4 (1 male). Data presented as mean ± s.e.m. Males = closed circles/solid lines, Females = open circles/dashed lines. \**P* < 0.05, \*\**P* < 0.01, \*\*\**P* < 0.001.

#### A subpopulation of DMS D1^+^ neurons stably encodes action initiation

We next investigated whether action encoding in individual DMS D1^+^ neurons shifts or persists as actions are learned and behavioral control transitions from goal-directed to habitual. Stability would suggest habit formation does not require the loss or replacement of previously established action representations. Using both prospective and retrospective approaches, we found remarkable stability of action encoding in individual DMS D1^+^ neurons across action-outcome learning and habit formation. We classified neurons as significantly excited by action initiation either during initial training (*‘early action-initiation excited’*) or during overtraining (*‘overtrain action-initiation excited’)*. Both populations of action-initiation D1^+^ neurons were more excited (Figure 2d-e, o-p) and modulated (Figure 2f, q; Extended Data Figure 2-2d-e) prior to the initiating lever press and encoded action initiation with high fidelity (Extended Data Figure 2-6). Looking prospectively, we found the early action-initiation excited ensemble continued to be active and modulated prior to action initiation throughout training and overtraining (Figure 2d-f; Extended Data Figure 2-2d). Looking retrospectively, we found the overtrain action-initiation excited ensemble was also active (Figure 2o-p) and modulated (Figure 2q) prior to action initiation during early and middle training. Further demonstrating ensemble stability, the activity around action initiation of both individual (Figure 2g, r) and the population (Figure 2h, s) of action-initiation excited D1^+^ neurons was significantly correlated across training phases. About half (Figure 2z-aa) the DMS D1^+^ action-initiation excited neurons stably encoded action initiation, significantly more than would be expected by chance (2-tailed t-test, t_3_ = 4.53, *P* = 0.02, 95% CI –8.27 – –1.45). The D1^+^ neuron ensemble that was inhibited around action initiation was similarly stable (Extended Data Figure 2-3). Thus, an ensemble of DMS D1^+^ neurons stably encodes action initiation throughout action-outcome learning and habit formation.

#### DMS D1^+^ action-initiation neurons stably convey information about actions, outcomes, and behavioral engagement and organization

We next asked whether the DMS D1^+^ neurons that stably encode action initiation also carry a broader and persistent representation of instrumental performance. We found D1^+^ action-initiation excited neurons contained and maintained information about ongoing actions, outcomes, and local fluctuations in behavioral engagement and organization. Actions, checking for, and receiving reward, could all be decoded with a multi-class SVM trained on the activity of either the D1^+^ early (Figure 2i) or overtrain (Figure 2t) action-initiation excited ensemble. Decoding of action initiation improved with training. Behavioral engagement (press rate and switch rate; Figure 2j-k, u-v), and organization (current relative interpress interval and past trial interpress interval; Figure 2l-m, w-x) could also be decoded across training using linear regression models trained on the activity of D1^+^ early action-initiation excited neurons. We also found coding-property stability. Using a linear regression model trained to decode action rate from the activity of the early action-initiation excited D1^+^ ensemble on the 1^st^ training session, we could decode action rate not only for the early session, but also from the middle and overtraining sessions (Figure 2n). Similarly, using a linear regression model to decode action rate from the activity of overtrain action-initiation excited D1^+^ neurons on the last training session, we could reliably decode action rate not only from this session, but also the preceding training sessions (Figure 2y). Thus, DMS D1^+^ neurons that stably encode action initiation maintain a broader representation of instrumental behavior, indicating that information about ongoing actions, outcomes, and behavioral organization remains available within the same neuronal population even as behavioral control shifts from goal-directed to habitual.

#### Stable DMS D1^+^ action-initiation neurons are conjunctively activated by action and reward outcome with learning

Having established that DMS D1^+^ neurons maintain stable action representations, we next asked whether these neurons come to encode other events with learning. We were particularly interested in whether learning recruits earned-reward responses into the stable D1^+^ neuronal population representing the action. Indeed, we found that stable action-initiation excited neurons also came to be excited by action termination and, critically, earned reward with learning (Figure 2ab-ac; Extended Data Figure 2-7). The stable action-initiation excited D1^+^ neurons showed action bracketing^66–68^, being excited before the start and end of an action bout. The percentage of stable action-initiation neurons excited by both action initiation and termination grew from 41.3% to 87.03% with training. The proportion of the stable action-initiation neurons that were also excited by earned reward grew from 18.7% to 63.9%, significantly more than would be expected by chance (2-way ANOVA, Neuron activity v. chance: F_1, 3_ = 22.42, *P* = 0.02; 2-tailed, Bonferroni corrected t-tests actual v. chance, Early: t_6_ = 2.02, *P* = 0.27, 95% CI –1.31 – 5.45; Middle: t_6_ = 4.07, *P* = 0.02, 95% CI 0.81 – 7.56; Overtrain: t_6_ = 3.63, *P* = 0.03, 95% CI 0.35 – 7.10). This action and reward conjunction activation was not due to coding of the press-check motor behavior. Stable action-initiation excited neurons were not activated by similar behavioral events in which the port-check was not reinforced (one-way ANOVA, Training: F_1.15, 3.46_ = 1.10, *P* = 0.38). Interestingly, the D1^+^ subpopulation encoding earned reward was not stable across training (Extended Data Figure 2-5). Most of the D1^+^ neurons that maintained reward encoding and all of those that gained it with training also encoded action initiation (Extended Data Figure 2-5m-p). Those neurons that lost reward encoding with training were less likely to encode action initiation. Therefore, learning recruits earned-reward responses into the stable D1^+^ action-encoding ensemble, producing conjunctive action-reward activation.

### DMS D1^+^ neurons support goal-directed learning and decision making

#### DMS D1^+^ neuronal activity is necessary for the learning underlying goal-directed behavior

Having identified that DMS D1^+^ neurons encode instrumental behavior and its reward outcome, we next asked whether DMS D1^+^ neuronal activity supports the learning underlying goal-directed behavior. We found it does. To test this, we chemogenetically inhibited DMS D1^+^ neurons during learning. We chose chemogenetic manipulation because DMS D1^+^ neurons were endogenously active at multiple time points and this would avoid artificially synchronizing subpopulations of neurons with different activity profiles and the reinforcing/punishing^69,70^ and locomotor^71–73^ effects that can occur with optogenetic manipulations of these cells. We expressed the inhibitory designer receptor human M4 muscarinic receptor (hM4Di) selectively in DMS D1^+^ neurons of DRD1a-cre mice (Figure 3a-b). Mice were trained to lever press to earn food-pellet rewards on a random-interval reinforcement schedule (Figure 3c). We gave mice limited training (4 sessions) to preserve dominance of goal-directed behavior in wildtype controls. Prior to each training session, all mice received the hM4Di ligand clozapine-N-oxide (CNO; 2.0 mg/kg^74–76^ i.p.) to, in hM4Di-expressing DRD1a-cre subjects, inactivate DMS D1^+^ neurons (see Extended Data Figure 3-1 for validation). Chemogenetic inactivation of DMS D1^+^ neurons did not significantly alter acquisition of the instrumental behavior (Figure 3d; see Extended Data Figure 3-2 for food-port entry and presses/reward). Thus, DMS D1^+^ neuronal activity is not necessary for general acquisition or execution of instrumental behavior. Inhibition of D1^+^ neurons did slightly decrease behavioral switch rate and significantly decreased interpress interval consistency (Extended Data Figure 3-3). To evaluate goal-directed v. habitual behavioral control, training was followed by the outcome-specific devaluation test. No CNO was given on test. Whereas controls reduced lever pressing when the outcome was devalued, indicating goal-directed control, D1^+^ neuron inhibition during learning caused subsequent insensitivity to outcome devaluation (Figure 3e-f). This is consistent with impaired learning, updating, and/or strengthening of action-outcome association, but could also indicate an altered influence of goal-directed v. habitual control. Regardless, DMS D1^+^ neuronal activity during instrumental learning mediates the learning that supports flexible, goal-directed behavior.

**Figure 3:**
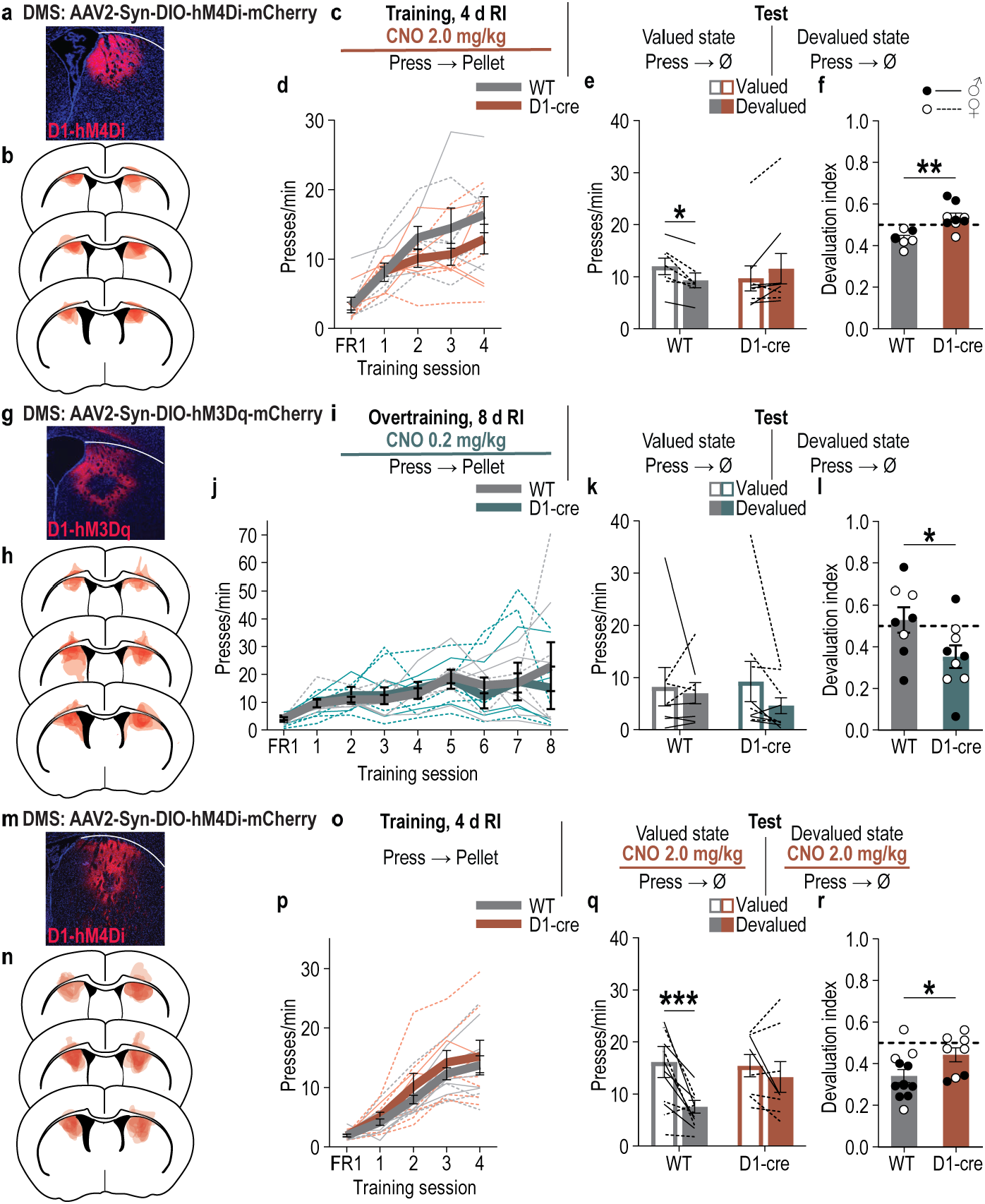
DMS D1^+^ neurons support goal-directed learning and decision making. (a-f) Chemogenetic inactivation of DMS D1^+^ neurons during learning. WT: *N* = 7 (2 males); D1-cre: *N* = 9 (5 males). **(a)** Representative immunofluorescent image of cre-dependent hM4Di expression in DMS. **(b)** Map of DMS cre-dependent hM4Di expression for all subjects. **(c)** Procedure. RI, random-interval reinforcement schedule; CNO, clozapine-N-oxide. **(d)** Training press rate. 2-way ANOVA, Training: F_2.22, 31.01_ = 27.65, *P* < 0.0001. **(e)** Test press rate. 2-way ANOVA, Value x Genotype: F_1, 14_ = 14.30, *P* = 0.002. **(f)** Devaluation index [(Devalued condition presses)/(Valued condition presses + Devalued presses)]. 2-tailed t-test, t_14_ = 4.01; *P* = 0.001, 95% CI –0.16 – –0.05. **(g-l)** Chemogenetic activation of D1^+^ neurons during overtraining. WT: *N* = 8 (5 males); D1-cre: *N* = 9 (4 males). **(g)** Representative immunofluorescent image of cre-dependent hM3Dq expression in DMS. **(h)** Map of DMS cre-dependent hM3Dq expression. **(i)** Procedure. **(j)** Training press rate. 2-way ANOVA, Training: F_2.44, 24.43_ = 3.23, *P* = 0.048. **(k)** Test press rate. 2-way ANOVA, Value x Genotype: F_1, 10_ = 3.37, *P* = 0.10. **(l)** Devaluation index. 2-tailed t-test, t_10_ = 3.07; *P* = 0.01, 95% CI –0.45 – –0.07. **(m-r)** Chemogenetic inactivation of D1^+^ neurons during test after learning. WT: *N* = 12 (7 males); D1-cre: *N* = 8 (2 males). **(m)** Representative immunofluorescent image of cre-dependent hM4Di expression in DMS. **(n)** Map of DMS cre-dependent hM4Di expression. **(o)** Procedure. **(p)** Training press rate. 2-way ANOVA, Training: F_1.59, 28.63_ = 68.95, *P* < 0.0001. **(q)** Test press rate. 2-way ANOVA, Value x Genotype: F_1, 18_ = 4.23, *P* = 0.05. **(r)** Devaluation index. 2-tailed t-test, t_18_ = 2.15; *P* = 0.045, 95% CI 0.003 – 0.20. Data presented as mean ± s.e.m. Males = closed circles/solid lines, Females = open circles/dashed lines. \**P* < 0.05, \*\**P* < 0.01, \*\*\**P* < 0.001.

#### DMS D1^+^ neuronal activity is sufficient to promote goal-directed behavioral control

Because DMS D1^+^ neurons stably encode instrumental actions and maintain action-reward representational similarity and conjunctive activation after habits have formed, we reasoned that DMS D1^+^ neurons might help maintain goal-directed control. If this is true, then augmenting D1^+^ neuron activity should promote the dominance of goal-directed behavioral control even after overtraining. We found it does. To test this, we chemogenetically activated D1^+^ neurons during learning and overtraining and then probed behavioral control using the devaluation test. We expressed the excitatory designer receptor human M3 muscarinic receptor (hM3Dq) selectively in DMS D1^+^ neurons of DRD1a-cre mice (Figure 3g-h). Mice were trained to lever press to earn food-pellet rewards (Figure 3i) and were overtrained (8 sessions) to promote habit formation. Prior to each training session, mice received the hM3Dq ligand CNO (0.2 mg/kg^77–80^ i.p.) to, in hM3Dq-expressing DRD1a-cre subjects, activate D1^+^ neurons (Extended Data Figure 3-1). Chemogenetic activation of DMS D1^+^ neurons did not alter acquisition of the instrumental behavior (Figure 3j; see also Extended Data Figure 3-2d-f and Extended Data 3-3d-f), but did promote goal-directed behavioral control. Whereas controls showed evidence of habit formation via insensitivity to devaluation, subjects for which we activated D1^+^ neurons during learning were sensitive to devaluation, indicating the dominance of goal-directed behavior (Figure 3k-l). Thus, enhanced DMS D1^+^ neuronal activity during learning promotes dominance of flexible, goal-directed behavior.

#### DMS D1^+^ neuronal activity mediates goal-directed decision making

We next reasoned that DMS D1^+^ neurons may function online to support flexible, goal-directed decision making. If this is true, then D1^+^ neuronal activity should be required to express goal-directed behavior by adapting when the outcome value changes. We found it is. To test this, we chemogenetically inhibited D1^+^ neurons at test, rather than during learning. We expressed hM4Di selectively in DMS D1^+^ neurons of DRD1a-cre mice (Figure 3m-n). Mice received limited training to lever press to earn food-pellet rewards (Figure 3o). All mice acquired the instrumental behavior (Figure 3p). Mice then received the outcome-specific devaluation test. After sensory-specific satiety, prior to the lever-pressing probe test mice received CNO (2.0 mg/kg i.p.) to inactivate D1^+^ neurons. Controls showed flexible, goal-directed decision making, reducing action performance when the outcome was devalued (Figure 3q-r). Chemogenetic D1^+^ neuron inhibition caused insensitivity to devaluation (Figure 3q-r). Inhibition of D1^+^ neurons, therefore, prevented adaptive goal-directed control. Thus, DMS D1^+^ neurons mediate both the learning underlying goal-directed control and flexible, goal-directed decision making.

### The DMS A2A^+^ neuronal population conveys information about instrumental behavior, but does not develop action-reward representational similarity

We next investigated how DMS A2A^+^ neuronal activity relates to instrumental behavior during action-outcome learning and habit formation using cellular-resolution, microendoscopic imaging of jGCaMP7s selectively expressed in A2A^+^ neurons of A2A-cre mice (Figure 4a-d). As with D1^+^ neurons, we longitudinally recorded calcium activity in A2A^+^ neurons as mice learned to lever press to earn food-pellet rewards on a random-interval reinforcement schedule and then probed goal-directed v. habitual behavioral control with an outcome-specific devaluation test (Figure 4e). All mice acquired the instrumental behavior (Figure 4f; see Extended Data Figure 4-1 for food-port entry data). Following overtraining, half the mice were insensitive to devaluation (Figure 4g-h), indicating routine habit formation. Unexpectedly, half the A2A-cre mice did not form habits with overtraining. To capture activity related to goal-directed behavior and habit formation, we focused on the subjects that formed habits and analyzed and coregistered (Figure 4d; Extended Data Figure 4-2) calcium activity across the beginning (session 1, RI-15s), middle (session 4, RI-30s), and end of training (session 8, RI-30s; Supplemental Table 1).

**Figure 4:**
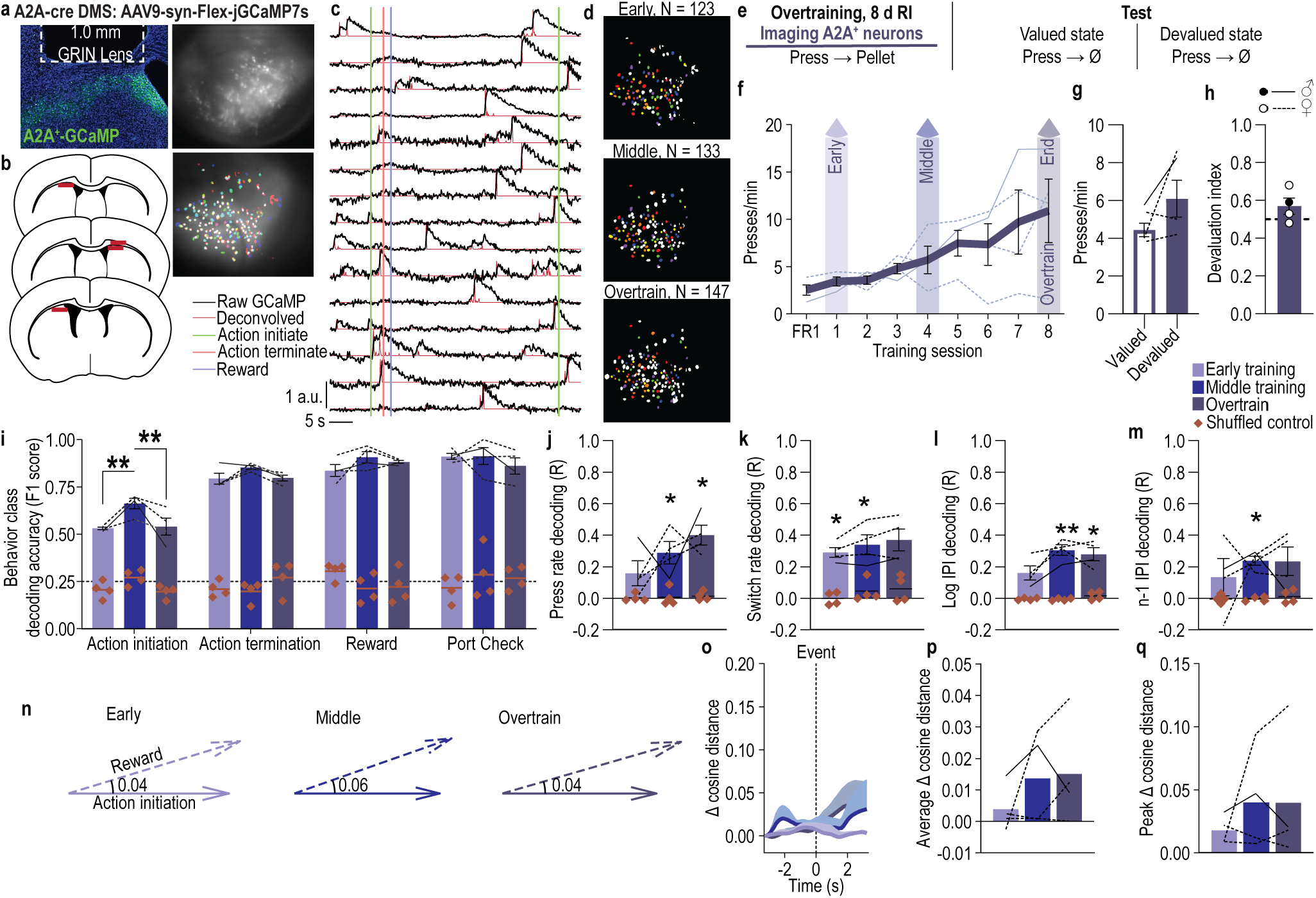
The DMS A2A^+^ neuronal population encodes instrumental behavior. **(a)** Representative image of cre-dependent jGCaMP7s expression in DMS A2A^+^ neurons (top left) and maximum intensity projection of the field-of-view with (top right) and without (bottom right) demarcation of cell ROIs for one session. **(b)** Map of DMS GRIN lens placements. **(c)** Representative raw fluorescence (*F* normalized to max) and deconvolved calcium signal. **(d)** Representative recorded A2A^+^ neuron spatial footprints during the first (early), 4^th^ (middle), and 8^th^ (overtrain) random-interval training sessions. Colored, coregistered neurons; white, non-coregistered neurons. **(e)** Procedure. RI, random-interval reinforcement schedule. **(f)** Training press rate. 1-way ANOVA, Training: F_1.39, 4.18_ = 3.56, *P* = 0.12. **(g)** Test press rate. 2-tailed t-test, t_3_ = 1.50, *P* = 0.23, 95% CI –1.86 – 5.16. **(h)** Devaluation index [(Devalued condition presses)/(Valued condition presses + Devalued presses)]. One-tailed Bayes factor, BF_10_ = 0.19. **(i)** Behavior class decoding accuracy from A2A^+^ coregistered neuron activity in 0.4-s period prior to event compared to shuffled control. Line = chance. 3-way ANOVA, Neuron activity (v. shuffled): F_1, 3_ = 1570.57, *P* < 0.001. **(j-m)** Linear decoding accuracy from A2A^+^ coregistered neuron activity of press rate (j; 2-way ANOVA, Neuron activity: F_1, 3_ = 65.01, *P* = 0.004), switch rate (k; 2-way ANOVA, Neuron activity: F_1, 3_ = 42.56, *P* = 0.007), log-transformed current interpress interval (l; 2-way ANOVA, Neuron activity: F_1, 3_ = 70.36, *P* = 0.004), and the prior press interpress interval (m; 2-way ANOVA, Neuron activity: F_1, 3_ = 82.59, *P* = 0.003). R = correlation between observed and decoded values. **(n)** Visualization of representative single-subject, single-timepoint example population vectors with cosine distance between action-initiation (“action”) and earned-reward (“reward”) neuronal population vectors. **(o-q)** Change (Δ) in cosine angular distance, relative to pre-event baseline, between action-initiation and earned-reward A2A^+^ neuronal population vectors across time and training centered around each event (o), average (p; 1-way ANOVA, Training: F_1.42, 4.26_ = 1.09, *P* = 0.39), and maximum (q; peak; 1-way ANOVA, Training: F_1.12, 3.35_ = 0.67, *P* = 0.48) ± 2.5 s around each event. Data presented as mean ± s.e.m. A2A-cre: *N* = 4 (1 male). Males = closed circles/solid lines, Females = open circles/dashed lines. \**P* < 0.05, \*\**P* < 0.01.

**Figure 5:**
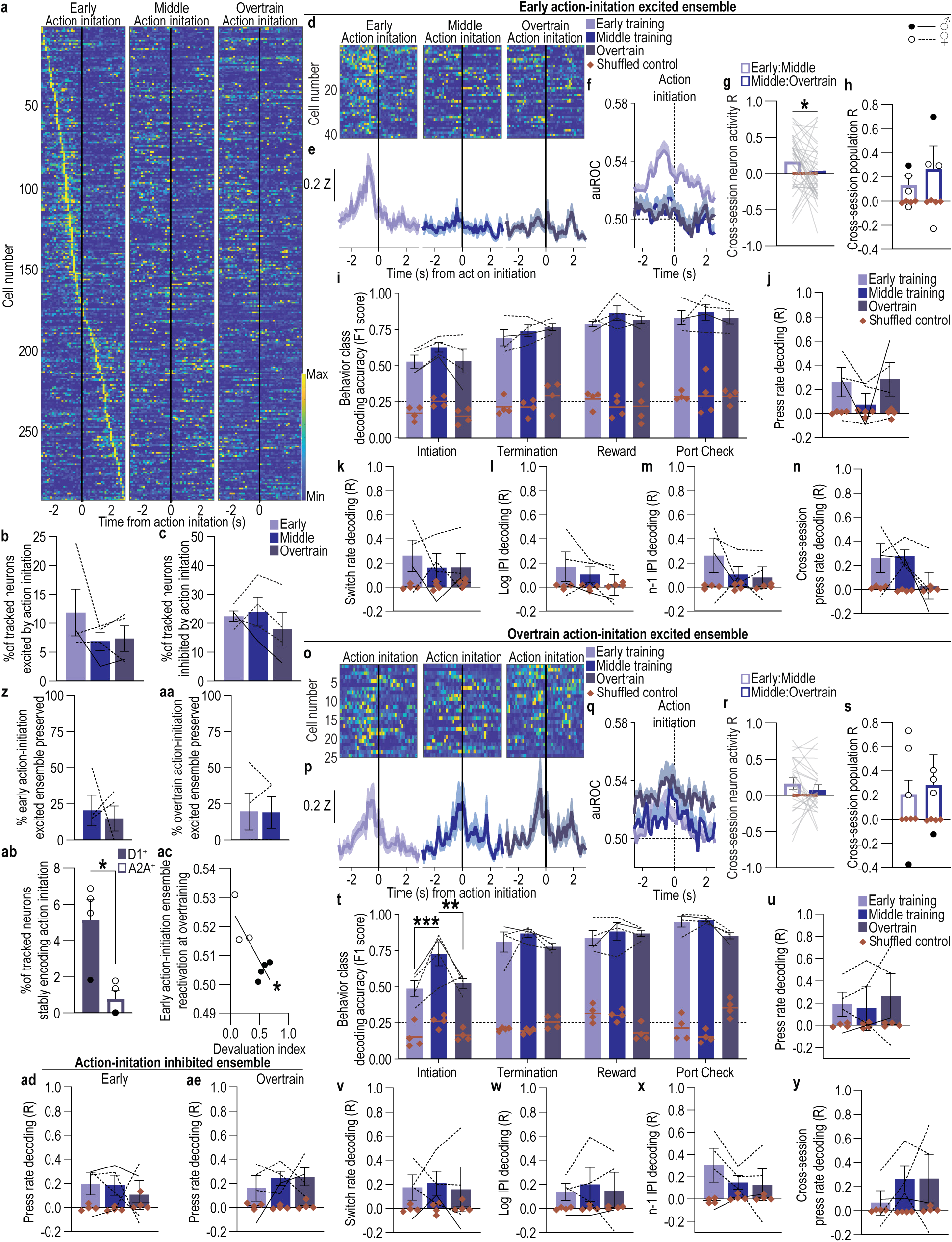
The subpopulation of DMS A2A^+^ neurons that encodes action initiation reorganizes with habit formation. **(a)** Heat map of minimum to maximum deconvolved activity (sorted by total activity) of each coregistered DMS A2A^+^ neuron around lever-press action initiation. **(b-c)** Percent of coregistered neurons (Average 73.25 coregistered neurons/mouse, s.e.m. 17.09) significantly excited (b; 1-way ANOVA, F_1.16, 3.48_ = 1.79, *P* = 0.27) or inhibited (c; 1-way ANOVA, F_1.05, 3.16_ = 1.14, *P* = 0.37) around action initiation. **(d-f)** Activity and modulation across training of A2A^+^ early action-initiation excited neurons. Heat map (d), Z-scored activity (e), and area under the receiver operating characteristic curve (auROC) modulation index (f) of early action-initiation excited neurons around action initiation across training. **(g-h)** Cross-session correlation of the activity around action initiation of each early action-initiation excited neuron (g; 2-way ANCOVA, Neuron activity: F_1, 38_ = 1.12, *P* = 0.30) or the population activity of these neurons (h; 2-way ANOVA, Neuron activity: F_1, 3_ = 3.07, *P* = 0.18). **(i)** Behavior class decoding accuracy from A2A^+^ early action-initiation excited neuron activity in the 0.4-s period prior to event compared to shuffled control. Line = chance. 3-way ANOVA, Neuron activity (v. shuffled): F_1, 3_ = 169.03, *P* < 0.001. **(j-m)** Linear decoding accuracy from A2A^+^ early action-initiation excited neuron activity of lever-press rate (j; 2-way ANOVA, Neuron activity: F_1, 3_ = 4.80, *P* = 0.12), switch rate (k; 2-way ANOVA, Neuron activity: F_1, 3_ = 3.74, *P* = 0.15), log-transformed current interpress interval (l; 2-way ANOVA, Neuron activity: F_1, 3_ = 1.10, *P* = 0.37), and the prior press interpress interval (m; 2-way ANOVA, Neuron activity: F_1, 3_ = 2.53, *P* = 0.21). R = correlation between observed and decoded values. **(n)** Cross-session decoding accuracy of lever-press rate from the activity of A2A^+^ early action-initiation-excited neuron population activity on the 1^st^ training session. Planned, Bonferroni corrected, 2-tailed t-tests, Early: t_6_ = 2.59, *P* = 0.12, 95% CI –0.07 – 0.55; Middle: t_6_ = 3.02, *P* = 0.07, 95% CI –0.02 – 0.59; Overtrain: t_6_ = 0.29, *P* > 0.9999, 95% CI –0.28 – 0.34. **(o-q)** Activity and modulation across training of A2A^+^ overtrain action-initiation excited neurons. Heat map (o), Z-scored activity (p), and auROC modulation index (q) of overtrain action-initiation excited neurons around action initiation across training. **(r-s)** Cross-session correlation of the activity around action initiation of each overtrain action-initiation excited neuron (r; 2-way ANCOVA, Neuron activity: F_1, 23_ = 0.18, *P* = 0.68) or the population activity of these neurons (s; 2-way ANOVA, Neuron activity: F_1, 3_ = 2.01, *P* = 0.25). **(t)** Behavior class decoding accuracy from A2A^+^ overtrain action-initiation excited neuron activity. 3-way ANOVA, Neuron activity: F_1, 3_ = 1296.08, *P* < 0.001. **(u-x)** Linear decoding accuracy from A2A^+^ overtrain action-initiation excited neuron activity of lever-press rate (u; 2-way ANOVA, Neuron activity: F1, 3 = 1.74, *P* = 0.28; Training: F_1.67, 5.01_ = 0.31, *P* = 0.71), switch rate (v; 2-way ANOVA, Neuron activity: F_1, 3_ = 2.19, *P* = 0.24), log-transformed current interpress interval (w; 2-way ANOVA, Neuron activity: F_1, 3_ = 1.93, *P* = 0.26), and the prior press interpress interval (x; 2-way ANOVA, Neuron activity: F1, 3 = 2.59, *P* = 0.21). **(y)** Cross-session decoding accuracy of lever-press rate from the activity of A2A^+^ overtrain action-initiation-excited neuron population activity on the 8^th^ training session. Planned, Bonferroni corrected, 2-tailed t-tests, Early: t_6_ = 0.59, *P* > 0.999, 95% CI –0.29 to 0.41; Middle: t_6_ = 2.58, *P* = 0.13, 95% CI –0.08 – 0.62; Overtrain: t_6_ = 2.39, *P* = 0.16, 95% CI –0.10 – 0.60. **(z)** Percent of A2A^+^ early action-initiation excited neurons that continued to be significantly excited by action initiation on the 4^th^ and 8^th^ training sessions. 2-tailed t-test, t_3_ = 0.37, *P* = 0.74, 95% CI –53.97 – 42.86. **(aa)** Percent of A2A^+^ overtrain action-initiation excited neurons that were also significantly excited by action initiation on 1^st^ and 4^th^ training sessions. 2-tailed t-test, t_3_ = 0.13, *P* = 0.91, 95% CI –18.88 – 17.44. **(ab)** Percent of all coregistered D1^+^ and A2A^+^ neurons that were significantly modulated by action initiation across all phases of training (‘stable action-initiation ensemble’). 2-tailed t-test, t_6_ = 3.55, *P* = 0.01, 95% CI –7.34 – –1.35. **(ac)** Correlation between the stability of the early action-initiation excited A2A^+^ ensemble (average auROC modulation index of this ensemble prior to action initiation during overtraining) and devaluation index for all subjects (A2A-cre: *N* = 7; 4 habitual, 3 goal-directed, 1 excluded due to having no early action-initiation excited neurons). Pearson r_7_ = –0.82, *P* = 0.02. **(ad-ae)** Accuracy with which lever-press rate can be decoded from the activity of A2A^+^ early (ad, 2-way ANOVA, Neuron activity: F_1, 3_ = 9.03, *P* = 0.05) or overtrain (ae, 2-way ANOVA, Neuron activity: F_1, 3_ = 31.44, *P* = 0.01) action-initiation inhibited neurons. A2A-cre: *N* = 4 (1 male). Data presented as mean ± s.e.m. Males = closed circles/solid lines, Females = open circles/dashed lines. \**P* < 0.05, \*\**P* < 0.01.

#### DMS A2A^+^ neuronal population activity conveys information about actions, outcomes, and behavioral engagement and organization

We used decoding to establish the behavioral and task information available in DMS A2A^+^ neuronal population activity. Behavioral events, engagement, and organization could be decoded from DMS A2A^+^ neuron population activity. Using a multi-class SVM algorithm, action initiation, action termination, non-reinforced food-port checks, and earned reward could all be reliably decoded from A2A^+^ neuronal activity (Figure 4i). Decoding of action initiation initially improved with learning, but decreased with overtraining, suggesting fluctuation in action coding with action-outcome learning and habit formation. Using linear regression models, both behavioral engagement (press rate and switch rate) and organizational (log interpress interval and past trial interpress interval) variables that correlate with the extent of goal-directed v. habitual behavioral control (sensitivity to devaluation) could be decoded from the A2A^+^ neuronal population activity (Figure 4j-m; see also Extended Data Figure 4-3). Thus, the DMS A2A^+^ neuronal population can contain information about actions, outcomes, and local fluctuations in behavioral engagement and organization.

#### The DMS A2A^+^ neuronal population does not develop action-reward representational similarity

We next asked whether the DMS A2A^+^ neuronal population representation of action and earned reward became more similar with learning. We found it did not. The cosine distance between the population vectors for action initiation and reward did not significantly change with training (Figure 4n-q; see also Extended Data Figure 4-4 for concordant representational similarity analysis). Thus, the DMS A2A^+^ population representation of action and outcome does not become more similar with learning.

### DMS A2A^+^ neuronal encoding of action initiation reorganizes with habit formation

#### DMS A2A^+^ neurons are active around actions

We next examined how individual A2A^+^ neurons encode task events during action-outcome learning and the transition to habit. A substantial proportion of A2A^+^ neurons were significantly excited around, largely prior to, action initiation during each phase of training (Figure 5a-b; see Extended Data Figure 5-1 for encoding in whole population; see Extended Data Figure 5-2a-c for an example classification). Another ensemble of A2A^+^ neurons was inhibited by action initiation (Figure 5c; Extended Data Figure 5-3). Few tracked A2A^+^ neurons were excited by action termination (Extended Data Figure 5-4) or reward (Extended Data Figure 5-5), though substantial proportions were inhibited by these events (Extended Data Figure 5-1). Indeed, A2A^+^ neurons appeared to quiet their activity during consumption of the earned reward (Extended Data Figure 5-1). Thus, DMS A2A^+^ neurons encode actions during learning and habit formation.

#### The subpopulation of DMS A2A^+^ neurons that encodes action initiation reorganizes with habit formation

We next investigated whether action encoding in individual DMS A2A^+^ neurons shifts or persists as actions are learned and behavioral control transitions from goal-directed to habitual. We found it shifts. One subpopulation of A2A^+^ neurons transiently encodes actions during early action-outcome learning and another slowly starts to encode actions as habits form. Thus, shift in behavioral control is associated with reorganization of A2A^+^ neuronal action representations. We classified A2A^+^ neurons that were significantly excited by action initiation early in training (early action-initiation excited) or after overtraining (overtrain action-initiation excited). Both A2A^+^ neuron populations were more excited (Figure 5d-e, o-p) and modulated (Figure 5f, q; Extended Data Figure 5-2d-e) prior to action initiation than after and encoded action initiation with moderate fidelity (Extended Data Figure 5-6). The early A2A^+^ action-initiation excited neurons lost their encoding of action initiation with training (Figure 5f; Extended Data Figure 5-2d). Neither the activity around action initiation of individual (Figure 5g), nor the population (Figure 5h) of early action-initiation excited A2A^+^ neurons was significantly correlated across training. After overtraining only about 15% of the early action-initiation excited A2A^+^ neuron ensemble continued to significantly encode action initiation (Figure 5z). These neurons did not retune to become excited by other task events, although some became inhibited by actions or rewards (Extended Data Figure 5-7). The activity of the overtrain action-initiation excited A2A^+^ neurons was also not significantly correlated across training sessions (Figure 5r-s) and only ∼19% of the overtrain action-initiation A2A^+^ ensemble significantly encoded action initiation at earlier training phases (Figure 5aa). Rather, the activity around and modulation by action initiation of the overtrain action-initiation excited A2A^+^ ensemble grew with training (Figure 5o-q; Extended Data Figure 5-2e). This did not result from retuning. Overtrain action-initiation A2A^+^ neurons were largely not excited by other events earlier in training, although, a small proportion were inhibited by actions at earlier task phases (Extended Data Figure 5-7). Therefore, one subpopulation of A2A^+^ neurons transiently encodes actions during early action-outcome learning and another slowly starts to encode actions as habits form. This reorganization contrasts with the stable encoding of D1^+^ neurons (Figure 5ab).

#### DMS A2A^+^ action-initiation excited neurons only convey information about behavioral events not engagement or organization

We investigated the extent to which the early and overtrain A2A^+^ action-initiation neurons carry broader information about instrumental behavior and outcomes. We found that A2A^+^ action-initiation excited neurons contained information about ongoing actions and outcomes, but not the local fluctuations in behavioral engagement and organization that relate to goal-directed v. habitual behavioral control. Actions, checking for, and receiving reward could all be decoded with a multi-class SVM trained on the A2A^+^ early (Figure 5i) or overtrain (Figure 5t) action-initiation excited neuron activity. But none of the behavioral engagement or organizational variables could be decoded, beyond the temporally structured null model, with linear regression models trained on the activity of either A2A^+^ action-initiation excited population (Figure 5j-m, u-x). Demonstrating coding-property instability, we were unable to decode action rate across sessions (Figure 5n, y). Thus, the shifting DMS A2A^+^ neuronal representation of action conveys behavioral events, but not engagement or organization.

The DMS A2A^+^ ensemble that was inhibited around action initiation was similarly unstable (Extended Data Figure 5-3), but did contain information about instrumental behavior, including action rate (Figure 5ad-ae) and other behavioral engagement and organizational variables (Extended Data Figure 5-3). Thus, although not stable in their encoding, DMS A2A^+^ neurons inhibited by action initiation contain information about local fluctuations in behavioral engagement and organizational information.

#### Reorganization of DMS A2A^+^ neuronal action encoding is associated with habit formation

If reorganization of DMS A2A^+^ neuronal action-initiation encoding is associated with habit formation, it should not occur if habits do not form. To test this, we exploited the serendipity that half the A2A-cre subjects did not form habits with overtraining and found they showed more stable A2A^+^ neuronal action representation. Like subjects that formed habits, a substantial proportion of A2A^+^ neurons were activated around, largely prior to, action initiation in subjects that did not form habits (Extended Data Figure 5-8). In these mice, the A2A⁺ neurons maintained elevated responses to action initiation across training (Extended Data Figure 5-8). Critically, the stability of the early action-initiation excited A2A^+^ neurons, assessed by the extent to which it continued to be excited prior to action initiation after overtraining, negatively correlated with devaluation index (Figure 5ac). More stability was associated with greater sensitivity to devaluation and, thus, stronger goal-directed behavioral control. Therefore, habit formation is associated with DMS A2A^+^ neuronal reorganization of action encoding.

### DMS A2A^+^ neurons support the learning underlying goal-directed control

#### DMS A2A^+^ neuronal activity is necessary for the learning underlying goal-directed behavior

Given that a subpopulation of DMS A2A^+^ neurons encodes actions during early learning, we reasoned DMS A2A^+^ neuronal activity might enable the learning that supports goal-directed behavioral control. We found it does. To test this, we chemogenetically inhibited DMS A2A^+^ neurons during learning and then probed goal-directed v. habitual behavioral control using the devaluation test. We expressed hM4Di selectively in DMS A2A^+^ neurons of A2A-cre mice (Figure 6a-b). Mice were trained to lever press to earn food-pellet rewards (Figure 6c). Prior to each of the 4 random-interval training sessions, mice received CNO (2.0 mg/kg, i.p.) to, in hM4Di-expressing A2A-cre subjects, inactivate A2A^+^ neurons (Extended Data Figure 3-1). Chemogenetic inactivation of A2A^+^ neurons did not alter acquisition of the instrumental lever-press behavior (Figure 6d; see also Extended Data Figure 6-1). Thus, DMS A2A^+^ neurons are not necessary for general acquisition or execution of instrumental behavior. A2A^+^ neuron inhibition did cause more entries into the food-delivery port, suggesting an inability to suppress this competing behavior (Extended Data Figure 6-1), but did not affect other behavioral engagement or organizational variables (Extended Data Figure 6-2). Inhibition of A2A^+^ neurons during learning resulted in subsequent insensitivity to outcome devaluation (Figure 6e-f). Thus, DMS A2A^+^ neuronal activity enables the learning processes that support flexible, goal-directed decision making.

**Figure 6:**
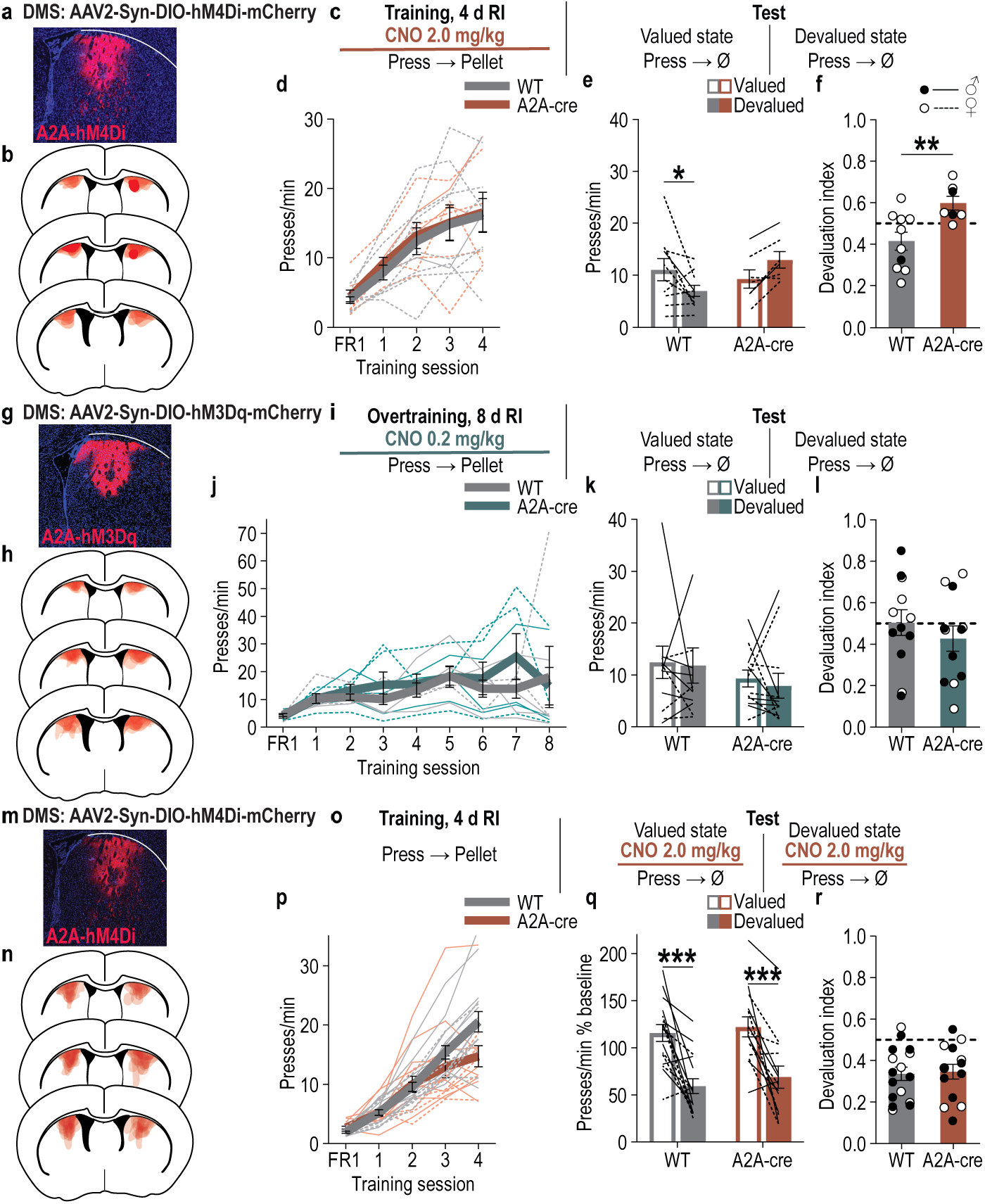
DMS A2A^+^ neurons support the learning underlying goal-directed control, but not goal-directed decision making. (**a-f**) Chemogenetic inactivation of DMS A2A^+^ neurons during learning. WT: *N* = 10 (1 male); A2A-cre: *N* = 7 (2 males). **(a)** Representative immunofluorescent image of cre-dependent hM4Di expression in DMS. **(b)** Map of DMS cre-dependent hM4Di expression for all subjects. **(c)** Procedure. RI, random-interval reinforcement schedule; CNO, clozapine-N-oxide. **(d)** Training press rate. 2-way ANOVA, Training: F_2.39, 35.84_ = 30.54, *P* < 0.0001. **(e)** Test press rate. 2-way ANOVA, Value x Genotype: F_1, 15_ = 9.67, *P* = 0.007. **(f)** Devaluation index [(Devalued condition presses)/(Valued condition presses ^+^ Devalued presses)]. 2-tailed t-test, t_15_ = 3.10; *P* = 0.007, 95% CI 0.057 – 0.31. **(g-l)** Chemogenetic activation of A2A^+^ neurons during overtraining. WT: *N* = 12 (7 males); A2A-cre: *N* = 12 (6 males). **(g)** Representative immunofluorescent image of DMS cre-dependent hM3Dq expression. **(h)** Map of DMS cre-dependent hM3Dq expression. **(i)** Procedure. **(j)** Training press rate. 2-way ANOVA, Training: F_1.87, 41.22_ = 15.50, *P* < 0.0001. **(k)** Test press rate. 2-way ANOVA, Value x Genotype: F_1,22_ = 0.03, *P* = 0.87. **(l)** Devaluation index. 2-tailed t-test, t_22_ = 0.89; *P* = 0.38, 95% CI –0.26 – 0.10. **(m-r)** Chemogenetic inactivation of A2A^+^ neurons during test after learning. WT: *N* = 16 (9 males); A2A-cre: *N* = 14 (8 males). **(m)** Representative immunofluorescent image of cre-dependent hM4Di expression in DMS. **(n)** Map of DMS cre-dependent hM4Di expression. **(o)** Procedure. **(p)** Training press rate. 2-way ANOVA, Training: F1.78, 49.82 = 107.5, *P* < 0.0001. **(q)** Test press rate normalized to pre-test training baseline. 2-way ANOVA, Value x Genotype: F_1, 28_ = 0.04, *P* = 0.84. **(r)** Devaluation index. 2-tailed t-test, t_28_ = 0.22; *P* = 0.83, 95% CI –0.09 to 0.11. Data presented as mean ± s.e.m. Males = closed circles/solid lines, Females = open circles/dashed lines. \**P* < 0.05, \*\**P* < 0.01, \*\*\**P* < 0.001.

#### DMS A2A^+^ neuronal activity is not sufficient to promote goal-directed behavioral control

We next asked whether augmenting A2A^+^ neuronal activity would promote the dominance of goal-directed behavioral control after overtraining. We reasoned that it would not because habit formation is associated with reorganization of DMS A2A^+^ neuronal action encoding. This prediction was borne out in the results. To test this, we chemogenetically activated A2A^+^ neurons during learning and then probed goal-directed v. habitual behavioral control using the devaluation test. We expressed hM3Dq selectively in DMS A2A^+^ neurons of A2A-cre mice (Figure 6g-h). Mice were trained to lever press to earn food-pellet rewards (Figure 6i) and were overtrained. Prior to each training session, mice received CNO (0.2 mg/kg, i.p.) to, in hM3Dq-expressing A2A-cre subjects, activate A2A^+^ neurons (Extended Data Figure 3-1). Chemogenetic activation of A2A^+^ neurons did not alter acquisition of the instrumental lever-press behavior (Figure 6j). It also did not prevent habit formation (Figure 6k-l). Both controls and subjects for which we activated A2A^+^ neurons during learning were insensitive to devaluation. Thus, DMS A2A^+^ neuronal activity is not sufficient to drive dominance of goal-directed behavioral control or disrupt habit formation.

#### DMS A2A^+^ neuronal activity is not necessary for goal-directed decision making

We next asked whether DMS A2A^+^ neurons function online to support goal-directed decision making by chemogenetically inhibiting DMS A2A^+^ neurons at test, rather than during learning. We expressed hM4Di selectively in DMS A2A^+^ neurons of A2A-cre mice (Figure 6m-n). Mice received limited training to lever press to earn food-pellet rewards (Figure 6o). All mice acquired the instrumental behavior (Figure 6p). Mice then received the outcome-specific devaluation test and were given CNO (2.0 mg/kg, i.p.) after sensory-specific satiety, prior to the lever-pressing probe test. Presses were expressed as a percent of baseline (training session prior to test) to normalize the pre-existing differences in press rate between groups (see Extended Data Figure 6-3 for press rate). Inactivation of A2A^+^ neurons did not affect flexible, goal-directed decision making, as evidenced by sensitivity to devaluation in both groups (Figure 6q-r). Thus, A2A^+^ neurons support the learning underlying goal-directed control, but become unnecessary for flexible, goal-directed decision making. This pattern of function differs from that of the DMS D1^+^ neurons, which support both goal-directed learning and decision making (2-way ANOVA on Devaluation index, Genotype x Inhibition period: F_2,78_ = 3.02, *P* = 0.05; Genotype (WT, D1-cre, A2A-cre): F_2,78_ = 5.55, *P* = 0.005; Inhibition period (training v. test): F_1,78_ = 22.22, *P* < 0.0001).

## DISCUSSION

Here we discovered a cell-type-specific dissociation between stable v. reorganizing DMS action representations and persistent v. transient contributions to goal-directed control. Early in learning both DMS neuron subtypes encode instrumental actions. D1^+^ neurons continue to stably encode instrumental actions and develop and maintain action-outcome representational similarity and conjunctive activation. Conversely, A2A^+^ neurons do not develop action-reward representational similarity and reorganize their action encoding with habit formation. Both DMS D1^+^ and A2A^+^ neurons support the learning underlying goal-directed behavior, but, whereas D1^+^ neurons enable flexible goal-directed decisions, A2A^+^ neurons become dispensable for such decisions.

DMS D1^+^ neurons stably encode instrumental actions, behavioral engagement and organizational information, and with learning come to conjunctively represent both the action and its earned outcome. Individual DMS D1^+^ neurons maintained encoding of instrumental action initiation across training. This stability likely results from post-synaptic plasticity in these neurons following action-outcome learning^81^. These neurons also stably conveyed behavioral engagement and organization variables that reflect task engagement, the structure of ongoing movements, behavioral stereotypy, and are predictive of goal-directed v. habitual control. With learning, neurons that stably represented the action also became activated by its earned reward, accompanied by increasing population-level action and reward representational similarity. This learning-dependent action-reward representational convergence is consistent with conjunctive action-outcome activation and extends previous evidence that striatal projection neurons multiplex specific actions and their reinforcing outcomes^82^ by revealing that learning causes the action and its consequence to be represented within a common, stable DMS D1^+^ neuronal ensemble.

The encoding stability and convergence of action and reward encoding could indicate DMS D1^+^ neurons form a lasting action-outcome code for flexible, goal-directed decision making. It could also reflect encoding of a learned behavioral chunk^66–68^. The absence of comparable coactivation and representational similarity between action and non-reinforced check behavior argues against a purely movement-based organizational chunk. However, firm evidence for an action-outcome code will require future experiments examining the extent to which D1^+^ neuronal action representations, action-reward representational similarity, and conjunctive coding are sensitive to action-outcome contingency, outcome value, or identity manipulations. Such experiments will also answer open questions about whether activity at action prospectively represents the expected reward and/or whether activity at reward retrospectively represents the action that produced it.

DMS D1^+^ neurons maintain encoding of instrumental behavior and conjunctive action-reward activation even after habits form. The transition from goal-directed to habitual control is well established to be associated with a shift in control away from DMS to the dorsolateral striatum^8,41,52^. Within this framework, it is tempting to predict that DMS representations of instrumental actions would weaken as habits form. To the contrary, DMS D1^+^ neuronal representations remain remarkably stable across learning, maintaining encoding of instrumental actions and action-reward representational similarity even once behavioral control has transitioned to a habit system likely independent of these neurons. That these representations persist even when behavior has become less dependent on goal-directed control suggests a lasting representation of actions and outcomes may remain available in DMS D1^+^ neurons should contingency or outcome value changes require renewed goal-directed control. This speculation is ripe for future investigation.

We found that D1^+^ neurons support both goal-directed learning and decision making. Inactivation of D1^+^ neurons during learning disrupted subsequent adaptive goal-directed control. This could have resulted from disruption to action-outcome learning, strengthening, refinement, or maintenance of this relationship, the ability of this relationship to adapt to the changed reinforcement schedule, or other features that would enable its influence relative to habitual control. Prior evidence that disruption to either DMS^43^ or DMS direct-pathway striatonigral projection activity prevents acquisition of new action-outcome associations argues that action-outcome learning is at least one likely function. Activating DMS D1^+^ neurons promoted the dominance of flexible, goal-directed decision making, similar to prior reports^83^. Inactivation of DMS D1^+^ neurons at test also disrupted expression of goal-directed decision making. This finding contrasts with prior evidence of preserved sensitivity to devaluation during optogenetic inactivation of DMS striatonigral projections^84^. This discrepancy may reflect differences in species, behavioral demands (single-lever v. 2-lever choice), the neuronal populations targeted (D1^+^ versus striatonigral neurons), or simply the extent and duration of inhibition achieved by chemogenetic v. optogenetic inhibition. Further studies are needed to clarify the precise contribution of DMS D1^+^ neurons to goal-directed decision making and the conditions under which their activity is necessary. Regardless, our results show that DMS D1^+^ neurons can support both the learning processes underlying goal-directed control and goal-directed decision making. Whether action-reward encoding neurons are critical to D1^+^ neuronal function in goal-directed control is an important question for future study.

DMS A2A^+^ neurons transiently encode actions and support the learning processes underlying goal-directed control. A subpopulation of A2A^+^ neurons transiently encodes action initiation early in learning. A2A^+^ neuronal activity during learning is necessary for subsequent goal-directed control. This finding is inconsistent with evidence that inhibiting DMS indirect striatopallidal projections does not disrupt acquisition of new action-outcome associations^84^. This discrepancy raises two intriguing possibilities. First, DMS A2A^+^ neurons may not contribute to initial action-outcome learning, but rather to the strengthening, refinement, and/or maintenance of this association across training and changes in reinforcement schedule, or otherwise enable goal-directed control. Second, rather than canonical indirect-pathway, striatopallidal projections, the DMS A2A^+^ neurons we find that function in goal-directed learning may be a distinct subtype^57,85–87^, perhaps a non-canonical projection. Both possibilities warrant further investigation.

DMS A2A^+^ neuronal encoding of instrumental behavior is distinct from that of D1^+^ neurons. A2A^+^ neurons show little encoding of reward outcomes, consistent with prior reports^88^ and do not show action-reward representational similarity or conjunctive activation. The A2A^+^ action-initiation excited neuronal ensemble does not convey behavioral engagement or organizational information predictive of the extent of goal-directed v. habitual control. Rather, this information is carried in A2A^+^ neurons inhibited around action initiation. Inhibition of DMS A2A^+^ neurons caused more transitions to the competing food-port check behavior. Thus, by inhibiting competing behaviors, DMS A2A^+^ neurons could help support goal-directed learning. This is consistent with the coordinated function view^71,89,90^ in which D1^+^ neurons drive selected actions and D2/A2A^+^ neurons permit such actions by inhibiting competing or unrewarded actions^68,69,71–73,89–95^. This coordinated D1^+^ and D2/A2A^+^ function model may, therefore, extend beyond motor control to learning.

DMS A2A^+^ neurons reorganize their action encoding with habit formation. This reorganization of action encoding may contribute to a function shift. In support of this, realignment was incomplete if habits did not form and instability of the A2A^+^ neuronal encoding of action initiation was associated with stronger habit formation (greater insensitivity to devaluation). Although A2A^+^ neurons initially support the learning processes underlying goal-directed control, they become unnecessary for goal-directed decision making and their activation is not sufficient to promote goal-directed control. Moreover, DMS A2A^+^ neurons have been implicated in habit^96–98^ and other behavioral strategy shifts including action-outcome reversal^84,99^ and extinction^99^. Thus, the DMS A2A^+^ neuronal action representation reorganizes as the behavioral control mode generating those actions shifts.

These discoveries open the door to many important future questions. The stability of D1^+^ and reorganization of A2A^+^ neuronal encoding likely require a combination of molecular^100–102^, cellular^81^, and circuit-level^49,102–104^ mechanisms. These may include cortical^49,102,103,105^, amygdala^104^, and/or midbrain dopamine^106–108^ input, and possibly interneurons^109^, astrocytes^110^, and collateral interactions^97,111,112^. Future studies will be crucial to define these mechanisms. The evidence here cannot resolve whether the observed stability of D1^+^ and reorganization of A2A^+^ neuronal encoding precedes, accompanies, or follows the transition to habit. This will require future studies combining longitudinal, cell-type-specific recordings with repeated behavioral assessments^113^ or behavioral state modeling^114^ approaches, independently validated against devaluation sensitivity, to provide trial-by-trial assessments of behavioral control across training. The task here was necessarily simple to allow neurons to be tracked from the initial learning through habit formation, but limits the depth to which we could explore action and outcome encoding. Thus, the features of the action, outcome, and action-outcome association encoded in DMS D1^+^ neurons will be important to investigate in the future. Being critical for goal-directed decision making, action value^115^ and outcome identity^116^ are both possible. Relatedly, the computations that allow behavior to flexibly adapt to outcome changes and extent to which they occur in the stable DMS D1^+^ action ensemble are important future questions. An intriguing larger question is whether stable D1^+^ neuronal encoding and A2A^+^ encoding realignment reflect broader principles of striatal contributions to learning and behavioral control. Our findings reveal that transitions in behavioral control need not erase established DMS D1^+^ neural action representations, raising the possibility that these persistent representations could contribute to renewed behavioral flexibility when circumstances change. A2A^+^ neuronal reorganization may enable shifts in behavioral control. Together, these findings provide a foundation for understanding how stress^104,105^, exposure to drugs and alcohol^26,50,117^, neurodevelopmental factors^93,118^, and aging^119^ may disrupt adaptive decision making and promote maladaptive habits.

## AUTHOR CONTRIBUTIONS

MM and KMW conceptualized and designed the experiments, interpreted the data, and wrote the paper. MM executed the experiments and analyzed the data. AL, NP, JYG, MDM, WG, Alicia W, MS, Anna W, JSP assisted with experiments. BSH conducted the on-devaluation-test chemogenetic inactivation experiments. SPB conducted the behavioral pilot experiment. JRG conducted the electrophysiological validation experiments with support from SMH, CC, and MSL. NKG managed the breeding colonies and assisted with histological verification. GJB, ACS, HTB, and AMW provided guidance and assistance with data analysis. PDB contributed guidance on behavioral modeling. AMW, HTB, and PDB provided guidance on data interpretation.

## ACKNOWLEDGEMENTS

This research was supported by NIH R01DA046679 (KMW), NIH R01DA058374 (KMW), BBRF 32148 (KMW), NIH T32DA024635 (JRG), NIH F32DA056201 (JRG), A.P. Giannini Fellowship (JRG), NIH K99MH135177 (JRG), NIH TL4GM118977 (NP), NIH F32MH135680 (GJB), NSF NeuroNex 1707408 (HTB), and the Staglin Center for Behavior and Brain Sciences. The authors thank Dr. Sean Ostlund for helpful comments on the behavioral analysis.

## METHODS

### Subjects

Male wildtype C57/Bl6J mice (Jackson Laboratories, Bar Harbor, ME) were used for the behavioral pilot experiment. All other experiments were conducted with male and female Drd1a-Cre^120^ (D1-cre) and Adora2A-Cre^121^ (A2A-cre) transgenic mice and their wildtype littermates bred in house. Mice were between 9 – 16 weeks old at the time of experiment onset/surgery. Mice were housed in a temperature (68 – 79 °F) and humidity (30 – 70%) regulated vivarium on a 12:12 hr reverse dark/light cycle (lights off at 7 AM). Mice were initially housed in same-sex groups of 3 – 4 mice/cage. Prior to experiment onset/surgery mice were single-housed to facilitate food deprivation and preserve implants. Unless noted below, mice were provided with food (standard rodent chow, Lab Diet, St. Louis, MO) and water *ad libitum* in the home cage. Mice were handled for 3 – 5 days prior to the start of behavioral training for each experiment. All procedures were conducted in accordance with the NIH Guide for the Care and Use of Laboratory Animals and were approved by the UCLA Institutional Animal Care and Use Committee.

### Surgery

Mice were anesthetized with isoflurane (3% induction, 1% maintenance), and positioned in a digital stereotaxic frame (Kopf, Tujunga, CA). Subcutaneous Rimadyl (Carprofen; 5 mg/kg; Zoetis, Parsippany, NJ) was given pre-operatively for analgesia and anti-inflammatory purposes. An incision was made along the midline to expose the skull. For viral infusions, after performing a small craniotomy, virus was injected using a 28-g infusion needle (PlasticsOne, Roanoke, VA) connected to a syringe (Hamilton Company, Reno, NV) by intramedic polyethylene tubing (BD; Franklin Lakes, NJ) and controlled by a syringe pump (Harvard Apparatus, Holliston, MA). Virus was injected at a rate of 0.1 µl/min and the needle was left in place for 10 min post-injection. Further experiment-specific surgical details are provided below. After surgery, mice were kept on a heating pad maintained at 35 °C for 1 hr and then single-housed in a clean homecage for recovery and monitoring. Mice received chow containing the antibiotic TMS for 7 days following surgery to prevent infection, after which they were returned to standard rodent chow.

### Behavioral procedures

#### Apparatus

Training took place in Med Associates (East Fairfield, VT) wide mouse operant chambers housed within sound– and light-attenuating boxes. Each chamber had metal grid floors and contained a retractable lever to the left of a recessed food-delivery port (magazine) on the front wall. A photobeam entry detector was positioned at the entry to the food port. Each chamber was equipped with 2 pellet dispensers to deliver either 20-mg grain or chocolate-flavored purified pellets (Bio-Serv, Frenchtown, NJ) into the food port when activated. A fan mounted to the outer chamber provided ventilation and external noise reduction. A 3-watt, 24-volt house light mounted on the top of the back wall opposite the food port provided illumination. To monitor subject behavior, monochrome digital cameras (Med Associates) were positioned with a top-down view of the conditioning chambers.

#### Food deprivation

3 – 5 days prior to the start of behavioral training, mice were food-deprived to maintain 85% – 90% of their free-feeding body weight. Mice were given 1.5 – 3.0 g of their home chow at the same time daily at least 2 hrs after training.

#### Outcome pre-exposure

To familiarize subjects with the food pellet that would become the instrumental outcome, mice were given 1 session of outcome pre-exposure. Mice were placed in a clean, empty cage and allowed to consume 30 of the food pellets from a metal cup. If any pellets remained, they were placed in the home cage overnight for consumption.

#### Magazine conditioning

Mice received 1 session of training in the operant chamber to learn where to receive the food pellets (20-mg grain or chocolate-purified pellets). Mice received 30 non-contingent pellet deliveries from the food port with a fixed 60-s intertrial interval.

#### Instrumental Training

Mice next received 1 session/day consecutively of instrumental conditioning in which lever presses earned delivery of a single food pellet. The same experimenter conducted each session of training and subsequent tests. Earned pellet type (grain or chocolate) was counterbalanced across subjects within each group of each experiment. Each session began with the illumination of the house light and insertion of the lever and ended with the retraction of the lever and turning off of the house light. Each training session ended after 30 outcomes had been earned or 30 min elapsed. In all cases, instrumental training began on a fixed-ratio 1 schedule (FR-1), in which each action was reinforced with one food pellet outcome. Once 90% of the maximum session outcomes were earned, the reinforcement schedule was shifted to random-interval (RI) in which a variable interval must elapse following a reinforcer for another press to be reinforced. Mice received either 4 or 8 (overtrained) consecutive sessions of RI training starting first with 1 session on a RI-15s schedule, and then the remaining sessions on a RI-30s schedule.

#### Alternate outcome exposures

To equate exposure of the non-trained pellet, all mice were given non-contingent access to the same number of the alternate food pellets as the earned pellet type (e.g., chocolate pellets if grain pellets served as the training outcome) in a different context (clear plexiglass cage).

#### Sensory-specific satiety outcome devaluation test

Testing began 24 hr after the final instrumental conditioning session. Mice were given 1 – 1.5 hr access to either 4 g of the food pellets previously earned by lever pressing (Devalued condition) or 4 g of the non-trained pellets to control for general satiety (Valued condition). The remaining pellets were weighed following prefeeding to measure total consumption. Consumption did not significantly differ between the Devalued v. Valued conditions or between the control and experimental group for any experiment (Supplemental Table 3). After prefeeding lever pressing was assessed during a brief, 5-min, non-reinforced probe test. To assess the efficacy of the sensory-specific satiety, following the probe test, mice were given a 10-min consumption choice test with simultaneous access to 1 g of both pellet types. To ensure that lever-press test data was not confounded by incomplete devaluation, subjects that failed to fully reject the devalued food type (i.e., consumed more than our margin of measurement error, 0.09 g, of the pre-fed pellet type during post-test consumption) were not included in the analysis (see exclusions noted for each experiment below). The remaining mice consumed less of the prefed pellet than non-prefed pellet, indicating successful sensory-specific satiety devaluation (Supplemental Table 4). 24 – 48 hr after the first devaluation test, mice received 1 session of instrumental retraining (RI-30s), followed the next day by a second devaluation test in which they were prefed the opposite food pellet. Thus, each mouse was tested in both the Valued and Devalued conditions, with test order counterbalanced across subjects within each group for each experiment.

### Establishing goal-directed behavior following limited instrumental random-interval training and habit following overtraining

Naïve, male C57BL/6J mice (8 weeks old, Jackson Laboratory) were used to establish the training protocols for action-outcome learning and goal-directed decision making (*N* = 8) or habit formation with overtraining (*N* = 8). No subjects were excluded. Mice were food restricted and then began instrumental training, as described above. Mice were habituated to intraperitoneal (i.p.) injections during the final day of FR-1 training. Then, all subjects received an i.p. injection of 0.9% saline following each of the RI training sessions. Following training, mice received a counterbalanced pair of sensory-specific satiety outcome-specific devaluation tests, as above. No injection was given on test.

### Microendoscopic calcium imaging

Naïve, male and female Drd1a-cre and A2A-cre mice (D1-cre: Final *N* = 4, 1 male; A2A-cre habitual *N* = 4, 1 male; A2A-cre goal-directed *N* = 4, 3 male) were used in this experiment to monitor calcium activity in individual D1^+^ or A2A^+^ DMS striatal projection neurons during instrumental conditioning. 18 (D1-cre: 9, A2A-cre: 9) subjects with lack of viral expression or without detectable individual neurons were excluded from the experiment. We collected data from 2 D1-cre mice that did not form habits, but these mice did not have sufficient detectable individual neurons (< 35 neurons) and so were excluded from analysis to avoid overfitting during population level analyses. This threshold was established prior to the onset of testing and was applied to all subjects in the experiment. Three (D1-cre: 2, A2A-cre: 1) subjects with viral overexpression were excluded from the dataset. At surgery, mice were infused with an adeno-associated virus (AAV) encoding the cre-dependent genetically encoded calcium indicator jGCaMP7s (0.40 µl; AAV9-syn-Flex-jGCaMP7s) into the DMS (AP +0.15, ML ±1.9, DV −2.65 mm from bregma). In a separate surgery, 4 – 7 days later, a unilateral 1.1-mm craniotomy (left/right hemisphere counterbalanced across subjects) was performed, centered above the DMS (AP +0.20, ML ±1.70 mm from bregma). Cortex was aspirated with a 27 – 30-g needle to a depth of approximately 1.8 mm. A 1-mm diameter, 4-mm long gradient refractive index (GRIN) lens (Inscopix, Palo Alto, CA) was implanted above the DMS (DV –2.25 mm). The lens was fixed to the skull with cyanoacrylate glue and secured with C&B Metabond quick adhesive cement system (Parkell Inc., Edgewood, NY), followed by opaque dental cement (Lang Dental Manufacturing, Wheeling, IL). Approximately 3 – 4 weeks following implant, a baseplate was attached to the previously formed headcap, ensuring proper alignment between the GRIN lens and the miniscope.

Mice were habituated to restraint and fitted with the miniscope for 2 days prior to behavioral training. Mice received a second magazine training session to become accustomed to retrieving pellets with the head-mounted miniscope. After the first FR-1 instrumental session, mice were subsequently fitted with the miniscope prior to each of 8 RI training sessions. Following training, mice received a pair of sensory-specific satiety outcome-specific devaluation tests, as above. We noticed that not all of the A2A-cre subjects developed habits (insensitivity to devaluation) with overtraining. We, therefore, separated these subjects into those that formed habits (Devaluation index ≥ 0.473; *N* = 4) and those that remained sensitive to devaluation showing goal-directed control (Devaluation index < 0.473; *N* = 4). These criteria are based on the mean ± s.e.m. of the devaluation index of all overtrained mice in our lab’s library of data (0.517, s.e.m. ±0.044 = 0.473 minimum). These groups did not differ in their acquisition of lever pressing during training (Training session: F_1.51, 9.04_ = 8.89, *P* = 0.01; Group: F_1, 6_ = 0.48, *P* = 0.52; Training x Group: F_8, 48_ = 1.02, *P* = 0.43). They also did not differ in the efficacy of devaluation (see Supplemental Tables 3 – 4).

#### Predicting devaluation index

To reveal behavioral variables that relate to the extent of goal-directed v. habitual control we used a linear regression model to ask whether and how behavior during the final training session predicts subsequent sensitivity to outcome devaluation. We pooled behavioral data from all the D1-cre and A2A-cre subjects (N = 12) so we could exploit the variability in sensitivity to devaluation for this model. We included measures of behavioral engagement: lever-press rate and switch rate (rate at which subjects switch from pressing to checking the food port or vice versa; a switch was defined as any time the animal switched from pressing to entering the food port or from entering the food port to press). We also included measures of behavioral patterning. On the random-interval reinforcement schedule a variable amount of time must elapse after an earned reward before another press will earn a reward. Thus, we focused our pattern measures on the time subjects waited in between presses (interpress interval, IPI). As a measure of overall pattern, we included the variability/consistency (standard deviation) of the log-transformed interpress intervals. As a measure of short-term patterning and the extent to which current trial behavior relates to past trial behavior, we included the correlation between the current interpress interval, n, and the interpress interval of the prior press, n-1^63,64^. In preliminary investigations, we also evaluated whether the influence of the interpress interval variables on devaluation sensitivity depended on whether there was a food-port check during the interval and found it did not and so focused on interpress interval. Press rate, switch rate, standard deviation of the log interpress intervals, and n:n-1 interpress interval did not significantly covary (*P* = 0.06 – 0.51). Using the MATLAB function *fitlm*, we built a linear regression model to measure the extent to which press rate, switch rate, standard deviation of the log interpress intervals, and n:n-1 interpress interval correlation predict subsequent sensitivity to devaluation, measured by the devaluation index. The model included the standard deviation of log-transformed interpress-intervals and n:n-1 interpress interval correlation as fixed effects, along with an interaction term between press rate and switch rate. Model performance was evaluated using MATLAB function *predict* in a 5-fold cross-validation procedure repeated 5 times. The average adjusted R-squared was calculated for each cross-validation and averaged across repetitions to provide a robust estimate of the model’s predictive power.

#### Miniscope system

Calcium imaging videos were recorded using an open-source, head-mounted UCLA Miniscope imaging device, which interfaces with the UCLA open-source Miniscope DAQ hardware and software (http://miniscope.org/) to capture the neuronal calcium dynamics with corresponding frame timestamps. A webcam (Logitech, San Jose, CA) recorded the Med Associates interface LEDs, which display light pulses coincident with task events (lever presses, food-port entry, pellet delivery). This video was simultaneously saved on the same computer alongside the calcium imaging video via the miniscope software with synchronized frame timestamps.

#### Calcium trace extraction and spike inference

Videos were initially processed using custom Python code^122^ to concatenate all the videos from a session into one tiff stack, spatially downsample by a factor of 2, and temporally downsample to an effective frame rate of ∼7.5 frames/s to accelerate image processing. The Python implementation of the Calcium Imaging Analysis package, ‘CaImAn’^123^, was then used to process the videos. We applied the nonrigid motion correction algorithm to remove tissue motion artifacts and then applied constrained non-negative matrix factorization for microendoscopic data to extract individual spatial contours and demix temporal fluorescence traces for each detected neuron. Individual fluorescent signals were deconvolved to estimate discrete spikes for each neuron^124^. The deconvolved traces, the estimation of discrete spikes that generate the calcium traces, were derived from the denoised fluorescence traces using CaImAn’s ‘*deconvolveCa*’ function. Neurons were coregistered across sessions using the open source package CellReg^125^. Activity for each neuron was Z-scored to the whole session to allow comparison of baseline and phasic activity across cells and subjects with different levels of GCaMP expression and/or lens proximity.

#### Behavior extraction

All behavior-event videos from a session were concatenated and downsampled as above. Using custom MATLAB code, the brightness of each LED on the interface was measured for each frame. A task event was considered to have occurred if the LED brightness was >2x the background brightness threshold (determined individually per recording), thus binarizing task events in each frame of the video. The extracted frame-by-frame event information was then interpolated to the timestamps of the calcium imaging video.

#### Cell classification

We used an analysis based on the area under the receiver operating characteristic curve to identify neurons tuned to key behavioral events: ‘action initiation’, first press in session, first press after earned reward, and first press after a non-reinforced food-port check; ‘action termination’, last press before earned reward collection and last press before a non-reinforced food-port check; ‘food-port check’, entries into the food delivery port, for those in which there was no reward present (a.k.a. non-reinforced entries); ‘reward consumption’, consumption of the earned food pellet occurring at the time of exit from the food-delivery port. The auROC provides a non-parametric measure of the separability between two distributions of data. To increase the temporal precision in linking neural activity to behavior, we used the deconvolved traces to distinguish between neuronal activity preceding v. in response to an event. For each neuron, we defined two distributions of activity: one from the activity aligned to the ±2.9 s window around repeated occurrences of a specific behavioral event and one from equated baseline periods in which subjects were not engaged in the task. We compared these two distributions of neuronal activity using the MATLAB *perfcurve* function to generate receiver operating characteristic curves of the true positive rate and the false positive rate and measured the area under the curve (auROC). To determine if a neuron significantly encoded an event, we generated a null distribution of auROC values by circularly shifting the activity trace for each neuron and recalculating the auROC 1000 times. Neurons with true auROC values >97.5% of the auROCs in the null distribution were considered significantly excited by an event and those with true auROC values <2.5% of the auROCs in the null distribution were considered significantly suppressed. The 2.9-s window before and after the event of interest was selected to ensure that the cell classification was not biased toward neurons modulated exclusively before or after the event, thereby capturing the range of event-related dynamics, including both anticipatory and reactive responses. To validate the robustness of the classification and characterize the temporal dynamics of these responses, the time at which neurons were significantly modulated within the ±2.9 s window, the auROC was further calculated at each individual frame within the ±2.9 s window around each event. To test whether action-initiation neuron identity is preserved across sessions above chance, we ran a 10000-fold Monte Carlo permutation simulation. For each individual subject, we simulated the null distribution of overlapping neurons based on that subject’s specific daily proportions of neurons modulated by action initiation and total tracked neurons. To test if daily neuronal overlap between action initiation and reward was above chance, we ran a Monte Carlo permutation simulation as described above. In this case, chance overlap was simulated using the single-session proportions of action– and reward-modulated neurons. In all simulations, differences in total neuron yield across animals was controlled by normalizing the observed overlap and the simulated chance overlap as a percentage of the total tracked neuron pool for that subject.

#### Behavior decoding

We used support vector machines (SVM) to decode key behavioral events (action initiation, action termination, food-port check, and reward consumption) using the activity of specific neuron populations. We used the temporal fluorescence trace because it is a continuous signal that can reflect recent neural activity that contains a more integrated signal important for changes between behaviors. To predict behavior events prior to onset, observation data used to train and test the classifier consisted of the 0.4-s epoch preceding each of the 4 distinct event labels. Since there were large variations in the number of each type of behavioral event, we controlled for the number of each event label. The ¼ of the data was used as the test set by randomly selecting a similar number of each event label. The remaining ¾ data was used as the training set and was balanced to have similar representation of each event class. First, for the behavior event labels that were overrepresented (usually food-port entries) in the training set, we calculated the median number for all event labels, and limited the overrepresented event labels to that value. Then, for the behavior event labels that were underrepresented in the training set, we used the synthetic minority oversampling technique (SMOTE)^130^ on the training set to generate synthetic activity samples to match the event label with the highest number of events in the reaminig train set. For each session, the classifier was trained (MATLAB function *fitcecoc*) to generate a model and behavior was predicted (MATLAB function *predict*) with the generated model on the remaining data. Decoding performance was averaged across 20 different random data splits. Decoding performance was assessed by computing the F-score, a standard measure of binary classification accuracy that weighs recall (proportion of actual positives that are correctly predicted) against precision (the proportion of positive predictions that are actually positive) of the classifier. It is calculated as the harmonic mean of precision and recall. We performed a 1000-fold shuffling procedure to confirm that the average decoding of randomized data is at the theoretical chance level of ¼ correct.

To examine whether the ongoing lever-press rate, switch rate, and interpress-interval fluctuation could be explained by the neuronal activity, we fit a linear model (MATLAB function: *fitlm*) using the temporal fluorescence traces as predictors and behavior as the response. The discrete lever presses were transformed into an ongoing behavior rate by convolving the lever presses with a half-gaussian-post event kernel width of 30 s and a standard deviation of the distribution of 6 s (see Extended Data Figure 1-5f for example). Switch points were identified as a shift from a lever press to a food-port entry or vice versa and were convolved with a gaussian kernel width of 30 s and standard deviation of 6. As a measure of how long the subject waited before pressing relative to overall session press wait times, we used log interpress intervals. Interpress intervals were assigned to the frame in which the current press occurred and were defined as current press (n) time ꟷ the time of the last press (n-1) before being log transformed. These were also convolved with a half-gaussian-post event kernel width of 30 s and a standard deviation of 6. As a measure of past trial action performance, we also calculated the n-1 interpress intervals. These were assigned to the frame in which the current press (n) occurred and were defined as the time of the last press (n-1) – the time of the press before that (n-2). These were also convolved with a gaussian-post event kernel width of 30 s and a standard deviation of 6. To evaluate the temporal specificity of the decoding model, we evaluated decoding performance as a function of the population circular shift magnitude (±240 s in 10 s steps). We observed a sharp, narrow peak in decoding accuracy centered at the true alignment (Δt = 0 s), with performance dropping to a low, stable baseline beyond shifts of ±30 s (Extended Data Figure 1-5c-e and 4-3c-e). Thus, for our primary analyses we chose a kernel window of 30 s, which also aligned with the RI-30s schedule of reinforcement. The standard deviation of 6 was used to account for the slow decay time constant of GCaMP7s, accommodating potential fluorescence accumulation across behavior events^127^. The predictor and response data were Z-scored. A model was trained using the initial ¾ of the data and behavior rates were predicted (MATLAB function *predict*) with the generated model on the remaining test data. Decoding performance was assessed by calculating the correlation between the predicted behavior rate and the actual behavior rate. We performed a 1000-fold circular shuffling procedure to compare decoding accuracy to that of a null model. We used a minimum shift window of ±60 s (2x the shift magnitude where decoding accuracy remains elevated) to ensure local co-fluctuations between behavior and neural activity were decoupled. The circular shift was applied to the session data on the population as a whole prior to the train-test split to preserve the internal temporal structure and autocorrelations of the neural and behavior data. Preservation of the internal temporal structure was confirmed for an example neuron and for the population by calculating the autocorrelation function for the raw and circularly shifted neural traces using the MATLAB function *xcorr* (Extended Data Figures 1-5a-b and 4-3a-b).

#### Modulation fidelity

To determine whether neural activity in response to a behavior was consistent across sessions at the neuron level, we averaged the deconvolved activity of a single neuron across all trials within a session to get a single response trace for a session, and compared it to its average trace for each session using a Spearman correlation. For the whole population, we averaged the activity of all of the neurons to get a population-level average response trace for each session and correlated these population-level traces across sessions. To determine whether individual neuron activity was consistent within each session, we calculated the Spearman correlation for the activity of a single neuron between all trials and averaged across correlations to get an overall within-session reliability measure for each neuron. We assessed the whole population within-session reliability by calculating the average activity trace for all neurons for each individual trial, calculated the Spearman correlation between trial activity, and calculated the average correlation as a measure of the whole population within-session reliability. In all cases, we performed a 1000-fold circular shuffling procedure to compare modulation fidelity to that of randomized data. To further assess the reliability of the neural population on a trial-by-trial basis, we quantified the percentage of trials in which each neuron responded, using a response threshold of 2x standard deviation of the trace activity during the 2 s prior to behavior event onset. The distribution of percent of trial response for all neurons was plotted to assess the response heterogeneity of the neural population across sessions.

#### High-dimensional population vector analysis

Neural activity from individual neurons was aligned to two key behavioral events: action initiation and reward. For each event, temporal fluorescence traces were averaged across trials. We analyzed the temporal dynamics of the population activity using a sliding window to create a time series of population vectors for each event. The neural activity within a 4-frame (∼ 0.5-s) time window was averaged to form a population vector representing the population’s activity at a specific time point. This was repeated with a 1-frame timestep size to generate population vectors for each event over the entire ±2.9-s window around the events. The similarity between the population representations of action initiation and reward at each timepoint was quantified by calculating the cosine distance between the two population vectors (MATLAB function *pdist*) at each time point. The cosine distance is defined as 1−cos(θ), where θ is the angle between the two vectors. This metric was chosen because it is sensitive to the pattern of neural activity and is insensitive to the overall firing rate or vector magnitude. The distance was calculated for each sliding window, resulting in a trajectory of real distance values over time. The average and peak cosine distance were quantified across the ±2.5-s window around the events. To control for global population geometry shifts, the change in cosine distance (Δ cosine distance) was quantified by computing a baseline during a 0.5 s window 3 s prior to the events and subtracted this from the peri-event cosine distance. To visualize representative examples, representative vector distances were selected at the timepoint where the average distance was greatest in the first training session. The magnitude of each vector was calculated using the Euclidean norm^65^ as the square root of the sum of the squared activity of all neurons. The angle in degrees between the two vectors was calculated as the inverse cosine of 1-cosine distance. We used an identical approach to compare population activity around action initiation to that around action termination, rewarded food-port checks to non-reward checks, and action initiation to non-reward checks.

#### Representational Similarity Analysis

We used a time-resolved Representational Similarity analysis to examine the geometry of the neural activity within the entire neural population for each occurrence of action initiation and reward across training sessions. For each session, an empirical Representational Dissimilarity Matrix (RDM) was constructed from the neural activity for all of the Action initiation and Reward occurrences. Each entry in the RDM represents the pairwise dissimilarity between the activity patterns of two events, calculated as 1 – Pearson’s correlation coefficient between the population activity for each event. The resulting matrix is symmetric, with a diagonal of zeros. A theoretical model RDM was generated to hypothesize the representational geometry between the same event types (e.g., Action initiation-Action initiation or Reward-Reward) as highly similar, represented as a pairwise dissimilarity of 0, while the representational geometry between different event types (e.g., Action initiation-Reward) as distinct, represented as a pairwise dissimilarity of 1. We then used Spearman’s rank correlation to compare the lower-triangular portion of the empirical RDM with the model RDM, using a sliding window, for each session. A high correlation between the empirical RDM and model RDM indicates the neural representation between action initiation and reward is dissimilar. A lower correlation indicates similar neural representations between events. A visualization of the empirical RDM’s across sessions for a representative subject was generated using multidimensional scaling (MDS; MATLAB function *mdscale*), to apply dimensionality reduction to the matrix of pairwise dissimilarities, capturing the relative dissimilarities between all occurrences of action initiation and reward. The representative RDMs were selected at the timepoint where the correlation between the empirical and model RDMs had the highest correlation (a.k.a. highest dissimilarity). To determine the correlation expected by chance, we performed a 1000-fold permutation of the event labels and recalculated the correlation. The observed correlation was considered significant if it exceeded the 95^th^ percentile of the null distribution.

### Chemogenetic inhibition of DMS D1^+^ neurons during instrumental learning

Naïve, male and female D1-cre mice (Final *N* = 9, 5 males) and wildtype littermate controls (*N* = 7, 2 males) served as subjects in this experiment to assess the necessity of DMS D1^+^ neuron activity for the learning that supports goal-directed decision making. 3 D1-cre subjects with off-target viral expression were excluded from the dataset. 3 (D1-cre: 2, WT: 1) subjects were excluded for ineffective sensory-specific satiety. At surgery, all mice received bilateral infusion of AAV encoding a cre-inducible inhibitory designer receptor human M4 muscarinic receptor (hM4Di; AAV2-Syn-DIO-hM4Di-mCherry; 0.3 µl) into the DMS (AP +0.2, ML ±1.8, DV −2.65 mm from bregma). After 2 weeks of recovery, mice were food restricted and then began instrumental training, as described above. Mice were habituated to i.p. injections during the final day of FR-1 training. Then, all subjects received an i.p. injection of water-soluble clozapine-n-oxide (CNO; 2.0 mg/kg; Hello Bio, Princeton, NJ) 30 min prior to each of the 4 RI training sessions. Following training, mice received a pair of sensory-specific satiety outcome-specific devaluation tests, as above. No CNO was given on test. CNO was given prior to the retraining session (RI-30s) in between tests.

### Chemogenetic activation of DMS D1^+^ neurons during instrumental overtraining

Naïve, male and female D1-cre mice (Final *N* = 9, 4 males) and wildtype littermate controls (*N* = 8, 5 males) served as subjects in this experiment to assess whether DMS D1^+^ neuron activity is sufficient to promote goal-directed behavioral control and prevent the habit formation that normally occurs with overtraining. 6 D1-cre subjects with off-target viral expression were excluded from the dataset. 4 (D1-cre: 1, WT: 3) subjects were excluded for ineffective sensory-specific satiety. 1 WT subject that ate twice as much prior to one test than the other was excluded from the dataset to prevent the confound of differential general satiety. At surgery, all mice received bilateral infusion of AAV encoding a cre-inducible excitatory designer receptor human M3 muscarinic receptor (hM3Dq; AAV2-Syn-DIO-hM3Dq-mCherry; 0.3 µl) into the DMS (AP +0.2, ML ±1.8, DV −2.65 mm from bregma). After 2 weeks of recovery, mice were food restricted and then began instrumental training, as described above. Mice were habituated to i.p. injections during the final day of FR-1 training. Then, all subjects received CNO (0.2 mg/kg i.p.; Hello Bio, Princeton, NJ) 30 min prior to each of 8 RI training sessions. Following training, mice received a pair of sensory-specific satiety outcome-specific devaluation tests, as above. No CNO was given on test days. CNO was given prior to the retraining session (RI-30s) in between tests.

### Chemogenetic inhibition of DMS D1^+^ neurons during goal-directed decision making

Naïve, male and female D1-cre mice (Final *N* = 8, 2 males) and wildtype littermate controls (*N* = 12, 7 males) served as subjects in this experiment to assess whether DMS D1^+^ neuron activity is necessary for the expression of goal-directed decision making. 7 D1-cre subjects with off-target viral expression were excluded from the dataset. At surgery, all mice received bilateral infusion of AAV encoding cre-inducible hM4Di (AAV2-Syn-DIO-hM4Di-mCherry; 0.3 µl) into the DMS. After 2 weeks of recovery, mice were food restricted and then began instrumental training, as described above. Mice were habituated to i.p. injections during the final day of FR-1 training. Mice received an i.p. injection of 0.9% saline 30 min prior to each of 4 RI training sessions. Following training, mice received a pair of sensory-specific satiety outcome-specific devaluation tests, as above. Immediately after the pre-feeding, all mice received CNO (2.0 mg/kg i.p.) and commenced to the non-rewarded probe test 30 min later. Saline was given prior to the retraining session (RI-30s) in between tests.

### Chemogenetic inhibition of DMS A2A^+^ neurons during instrumental learning

Naïve, male and female A2A-cre mice (Final *N* = 7, 2 males) and wildtype littermate controls (*N* = 10, 1 male) served as subjects in this experiment to assess whether DMS A2A^+^ neuron activity is necessary for the learning that supports goal-directed decision making. 3 A2A-cre subjects with off-target viral expression were excluded from the dataset. 3 (A2A-cre: 1, WT: 2) subjects were excluded for ineffective sensory-specific satiety. At surgery, all mice received bilateral infusion of AAV encoding cre-inducible hM4Di (AAV2-Syn-DIO-hM4Di-mCherry; 0.3 µl) into the DMS. After 2 weeks of recovery, mice were food restricted and then began instrumental training, as described above. Mice were habituated to i.p. injections during the final day of FR-1 training. Then, all subjects received CNO (2.0 mg/kg i.p.) 30 min prior to each of 4 RI training sessions. Following training, mice received a pair of sensory-specific satiety outcome-specific devaluation tests, as above. No CNO was given on test. CNO was given prior to the retraining session (RI-30s) in between tests.

### Chemogenetic activation of DMS A2A^+^ neurons during instrumental overtraining

Naïve, male and female A2A-cre mice (Final *N* = 12, 6 males) and wildtype littermate controls (*N* = 12, 7 males) served as subjects in this experiment to assess whether DMS A2A^+^ neuron activity is sufficient to promote goal-directed behavioral control and prevent the habit formation that normally occurs with overtraining. 8 A2A-cre subjects with off-target viral expression were excluded from the dataset. 1 WT subject was excluded due to mechanical lesion. 11 (A2A: 6, WT: 5) subjects were excluded for ineffective sensory-specific satiety. At surgery, all mice received bilateral infusion of AAV encoding a cre-inducible hM3Dq (AAV2-Syn-DIO-hM3Dq-mCherry; 0.3 µl) into the DMS. After 2 weeks of recovery, mice were food restricted and then began instrumental training, as described above. Mice were habituated to i.p. injections during the final day of FR-1 training. Then, all subjects received CNO (0.2 mg/kg i.p.) 30 min prior to each of 8 RI training sessions. Following training, mice received a pair of sensory-specific satiety outcome-specific devaluation tests, as above. No CNO was given on test. CNO was given prior to the retraining session (RI-30s) in between tests.

### Chemogenetic inhibition of DMS A2A^+^ neurons during goal-directed decision making

Naïve, male and female A2A-cre mice (Final *N* = 14, 8 males) and wildtype littermate controls (*N* = 16, 9 males) served as subjects in this experiment to assess whether DMS A2A^+^ neuron activity is necessary for the expression of goal-directed decision making. 6 A2A-cre subjects with off-target viral expression were excluded from the dataset. At surgery, all mice received bilateral infusion of AAV encoding cre-inducible hM4Di (AAV2-Syn-DIO-hM4Di-mCherry; 0.3 µl) into the DMS. After 2 weeks of recovery, mice were food restricted and then began instrumental training, as described above. Mice were habituated to i.p. injections during the final day of FR-1 training. Then, all subjects received saline (0.9% i.p.) 30 min prior to each of 4 RI training sessions. Following training, mice received a pair of sensory-specific satiety outcome-specific devaluation tests, as above. Immediately after the pre-feeding, all mice received CNO (2.0 mg/kg i.p.) and commenced to the non-rewarded probe test 30 min later. Saline was given prior to the retraining session (RI-30s) in between tests. A2A-cre and control mice had slightly different press rates on the last day of training, and so test data were normalized to baseline press rates.

### Electrophysiological validation of chemogenetic manipulations

Whole-cell patch clamp recordings were used to validate the efficacy of chemogenetic manipulation of DMS D1^+^ and A2A^+^ neurons. Male and female naïve Drd1a-cre and A2A-cre mice served as subjects. At surgery, mice received bilateral infusion of AAV encoding cre-inducible hM4Di (AAV2-Syn-DIO-hM4Di-mCherry; 0.3 µl), cre-inducible hM3Dq (AAV2-Syn-DIO-hM3Dq-mCherry; 0.3 µl), or a fluorophore control (AAV2-Syn-DIO-mCherry; 0.3 µl) into the DMS. Recordings were performed from slices containing the DMS ∼2 – 3 weeks following infusion. Mice were ∼2 months old at the time slices were taken. Mice were deeply anesthetized with isoflurane and transcardially perfused with ice-cold, oxygenated NMDG-based slicing solution containing (in mM): 30 NaHCO_3_, 20 HEPES, 1.25 NaH_2_PO_4_, 102 NMDG, 40 glucose, 3 KCl, 0.5 CaCl2-2H_2_O, 10 MgSO_4_-H_2_O (pH adjusted to 7.3-7.35, osmolality 300-310 mOsm/L). Brains were extracted and placed in ice-cold, oxygenated NMDG slicing solution. Coronal slices (300 µm) were cut using a vibrating microtome (VT1000S; Leica Microsystems, Germany), transferred to an incubating chamber containing oxygenated NMDG slicing solution warmed to 32-34 °C, and allowed to recover for 15 min before being transferred to an artificial cerebrospinal spinal fluid (aCSF) solution containing (in mM): 130 NaCl, 3 KCl, 1.25 NaH_2_PO_4_, 26 NaHCO_3_, 2 MgCl_2_, 2 CaCl_2_, and 10 glucose) oxygenated with 95% O_2_, 5% CO_2_ (pH 7.2-7.4, osmolality 290-310 mOsm/L, 32-34°C). After 15 min, slices were moved to room temperature and allowed to recover for ∼30 additional min prior to recording. All recordings were performed using an upright microscope (Olympus BX51WI, Center Valley, PA) equipped with differential interference contrast optics and fluorescence imaging (QIACAM fast 1394 monochromatic camera 765 with Q-Capture Pro software, QImaging, Surrey, BC, Canada). Patch pipettes (3-5 MΩ resistance) contained a K-gluconate-based solution containing (in mM): 112.5 K-gluconate, 4 NaCl, 17.5 KCl, 0.5 CaCl_2_, 1 MgCl_2_, 5 ATP (K^+^ salt), 1 NaGTP, 5 EGTA, 10 HEPES, pH 7.2 (270–280 mOsm/L). Biocytin (0.2%, Sigma-Aldrich, St. Louis, MO) was included in the internal recording solution for subsequent verification of fluorescence labeling in recorded cells. Recordings were obtained using a MultiClamp 700B Amplifier (Molecular Devices, Sunnyvale, CA) and the pCLAMP 10.7 acquisition software. Fluorescence-guided whole-cell patch clamp recordings in current-clamp mode were obtained from DMS D1^+^ and A2A^+^ neurons (Control, *N* = 19 cells, 4 D1-cre mice, 1 male; D1-cre, hM4Di: *N* = 8 cells, 4 mice, 3 males, hM3Dq: *N* = 10 cells, 5 mice, 3 males; A2A-cre, hM4Di: *N* = 8 cells, 4 mice, 3 males, hM3Dq: *N* = 10 cells, 5 mice, 3 males). After breaking through the membrane, recordings were obtained from cells while injecting suprathreshold depolarizing current (1 s). Current injection intensities between 100 – 800 pA, resulting in 5 – 15 action potentials, were selected for recordings. Electrode access resistances were maintained at <30 MΩ throughout recordings. Number of action potentials generated was recorded both prior to and after bath application of CNO (1 or 100 µM) for >10 minutes. Percent change in the number of action potentials before and after CNO was calculated for both hM4Di and hM3Dq expressing cells.

### Histology

At the conclusion of instrumental training and testing, mice were euthanized and brain tissue was processed to assess viral expression location and spread and GRIN lens placement. Mice from miniscope experiments were anesthetized with isoflurane and transcardially perfused with ice-cold PBS followed by cold 4% paraformaldehyde. The brains were removed, post-fixed in 4% paraformaldehyde, then cryoprotected in 30% sucrose in PBS. 30-µm coronal slices were taken on a cryostat and collected in PBS. Sections were mounted on slides using ProLong Gold antifade reagent with DAPI (Invitrogen). Mice from chemogenetic experiments were anesthetized with isoflurane, brains were removed and rapidly frozen. 20-µm coronal slices were taken on a cryostat and dry mounted onto subbed slides. Slides were submerged in 4% paraformaldehyde, and coverlipped with ProLong Gold antifade reagent with DAPI (Invitrogen). All images were acquired using a Keyence (BZ-X710) microscope with 4X, 10X, and 20X objectives (CFI Plan Apo), CCD camera, and BZ-X Analyze software, to confirm viral expression and GRIN lens placement.

### Statistical Analysis

Datasets were analyzed by 2-tailed t-tests, one-tailed Bayes Factor, or 1-, 2-, or 3-way repeated-measures analysis of variance (ANOVA), or analysis of co-variance (ANCOVA), as appropriate (GraphPad Prism, GraphPad, San Diego, CA; MATLAB, SPSS, IBM, Chicago, IL). ANCOVA controlling for subject was used for all datasets in which cell rather than subject was the unit N. All data sets were checked for normality. Bonferroni post hoc tests corrected for multiple comparisons were performed to clarify statistical interactions. Chemogenetic validation data used uncorrected planned comparisons. Given single-session decoding results, corrected planned comparisons were used for cross-session decoding. Greenhouse-Geisser correction was applied to mitigate the influence of unequal variance between conditions. Alpha levels were set at *P* < 0.05.

### Sex as a biological variable

Male and female mice were used in approximately equal numbers, where possible based on breeding and mouse availability, but the *N* per sex was underpowered to examine sex differences. Sex was therefore not included as a factor in statistical analyses, though individual data points are visually disaggregated by sex.

### Rigor and reproducibility

See Supplemental Table 5 for key reagents. Group sizes were estimated based on prior work with this behavioral task^100^ and to ensure counterbalancing of virus, pellet type, and devaluation test order. Investigators were not blinded to viral group because they were required to administer virus. All behaviors were scored using automated software (Med Associates). Each experiment included at least 1 replication cohort and cohorts were balanced by Viral group or hemisphere (for imaging) prior to the start of the experiment. Investigators were blinded to group when performing histological validation and determining exclusions based on viral spread or mistargeted implant. Calcium analyses included cross-validation and comparison to shuffled data.

### Data and code availability

All data that support the findings of this study are available in the source data accompanying this paper and from the corresponding author upon request. Custom-written MATLAB code is accessible via Dryad repository https://doi.org/10.5061/dryad.5hqbzkhmx and available from the corresponding author upon request.

## EXTENDED DATA FIGURES

**Extended Data Figure 1-1:**
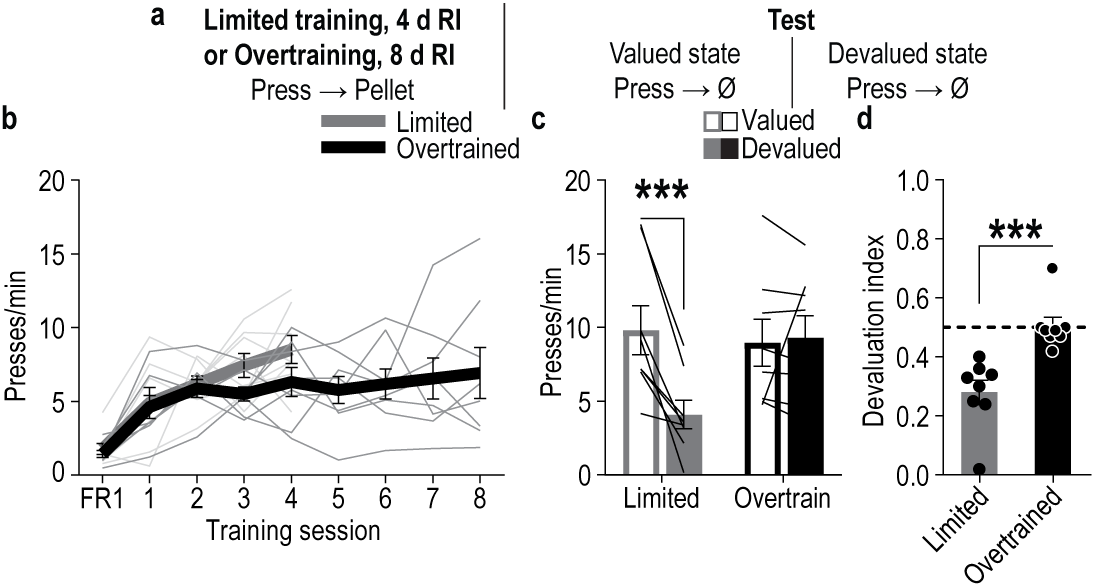
Behavior is goal-directed following limited random-interval instrumental training and habitual following overtraining. **(a)** Procedure. Lever presses earned food pellet rewards on a random-interval reinforcement schedule. Mice received either limited training (4 RI sessions) or overtraining (8 sessions). Mice were then given a lever-pressing probe test in the Valued state, prefed on untrained food-pellet type to control for general satiety and Devalued state prefed on trained food-pellet type to induce sensory-specific satiety devaluation. Test order was counterbalanced across subjects within each group, with a single intervening retraining session. **(b)** Training press rate. 1-way ANOVA, Limited training: Training: F_2.75, 19.27_ = 12.08, *P* = 0.0001. Overtraining: Training: F_2.24, 15.69_ = 4.56, *P* = 0.02. **(c)** Test press rate. 2-way ANOVA, Value x Training duration: F_1, 14_ = 14.69, *P* = 0.002; Value: F_1, 14_ = 11.69, *P* = 0.004; Training duration: F_1, 14_ = 1.31, *P* = 0.27. **(d)** Devaluation index [(Devalued presses)/(Valued presses + Devalued presses)]. 2-tailed Mann-Whitney U test, U = 0, P = 0.002. N = 8/group (all male). Data presented as mean ± s.e.m. \*\*\**P* < 0.001. Mice learn action-outcome relationships during instrumental conditioning on a random-interval schedule of reinforcement and use them for goal-directed decision making after limited training and form habits with overtraining.

**Extended Data Figure 1-2:**
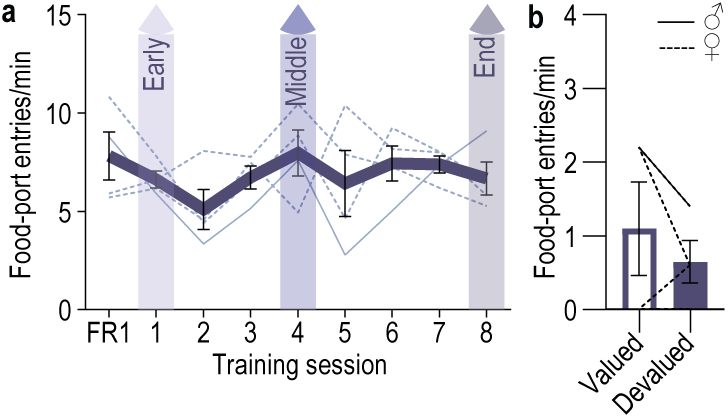
Entries into the food-delivery port during training and test for DMS D1^+^ imaging experiment. **(a)** Training entry rate. 1-way ANOVA, Training: F_2.11, 6.31_ = 0.75, *P* = 0.52 **(b)** Test entry rate. 2-tailed t-test, t_3_ = 0.94, *P* = 0.42, 95% CI –1.97 – 1.07. D1-cre: *N* = 4 (1 male). Data presented as mean ± s.e.m. Males = solid lines, Females = dashed lines.

**Extended Data Figure 1-3.**
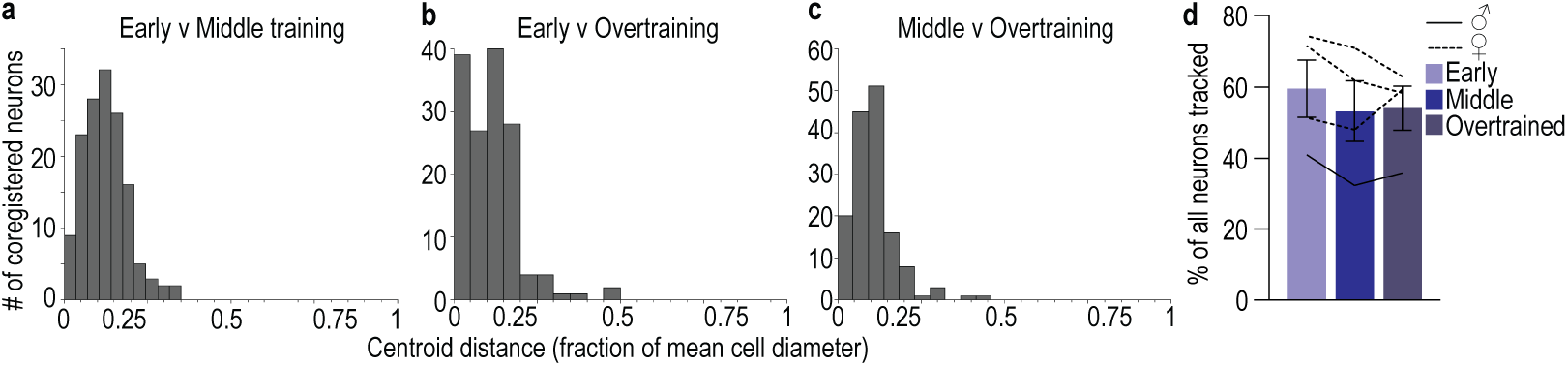
Coregistration of DMS D1^+^ neurons across training. (a-c) Representative example of coregistration of DMS D1^+^ neurons across training. Distribution of the distance between cell centroids (as a fraction of total cell diameter) of coregistered cell pairs in the 1^st^ (a, early) and 4^th^ (middle), 1^st^ and 8^th^ (b, overtrain), and the 4th and 8^th^ (c) training sessions. **(d)** Percent of all DMS D1^+^ neurons coregistered across training. 1-way ANOVA, F_1.19, 3.58_ = 1.62, *P* = 0.29. D1-cre: *N* = 4 (1 male). We were able to coregister on average 56% (s.e.m. = 3.90%) of the D1^+^ neurons across training. Data presented as mean ± s.e.m. Males = solid lines, Females = dashed lines.

**Extended Data Figure 1-4.**
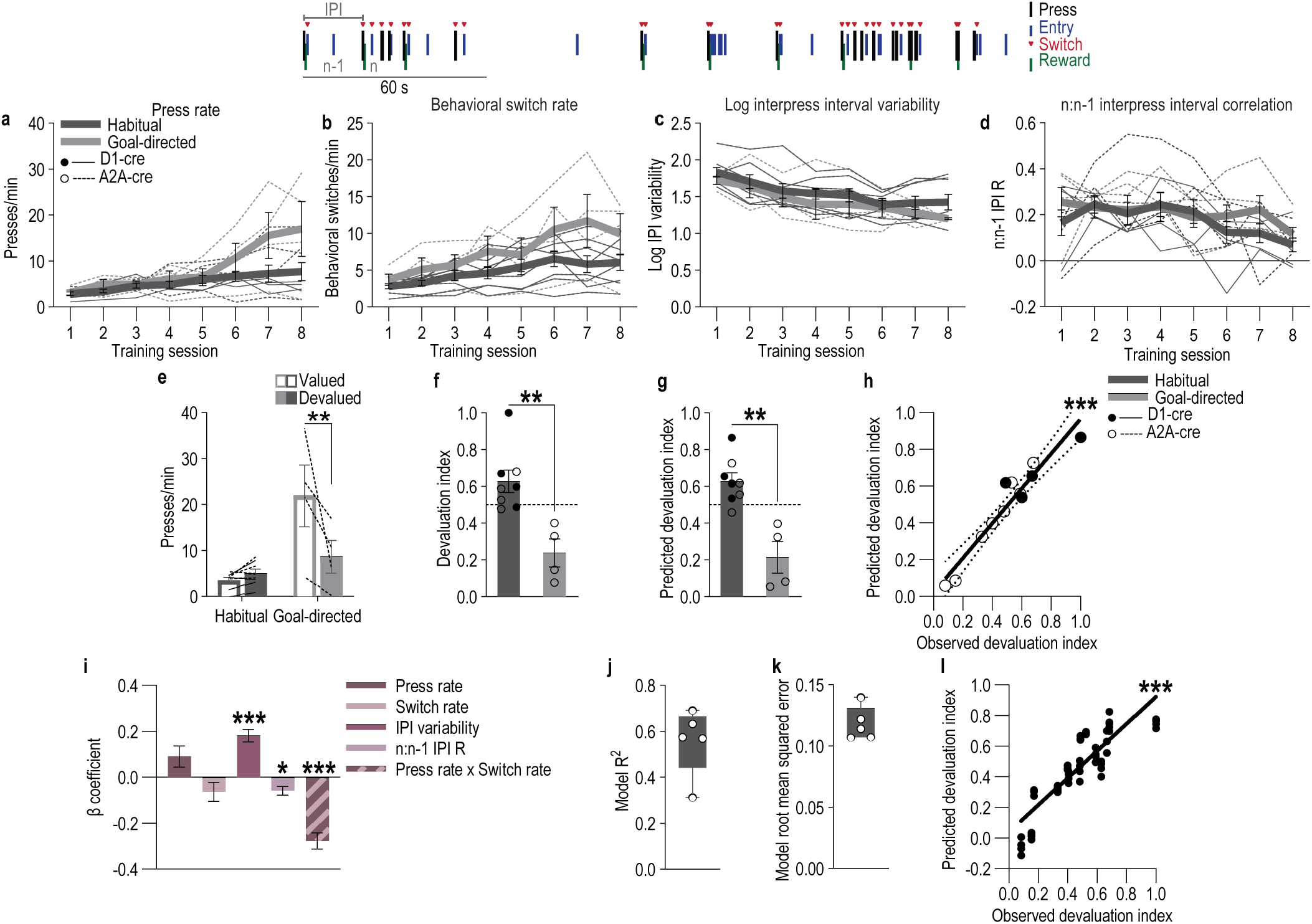
Linear regression model relating behavioral engagement and organizational variables to goal-directed v. habitual control. To reveal behavioral variables that relate to the extent of goal-directed v. habitual control we used a linear regression model to ask whether and how behavior during the final training session predicts subsequent sensitivity to outcome devaluation. We pooled behavioral data from all the D1-cre and A2A-cre subjects (*N* = 12) so we could exploit the variability in sensitivity to devaluation for this model. We included measures of behavioral engagement: lever-press rate and switch rate (rate at which subjects switch from pressing to checking the food port or vice versa). We also included measures of behavioral organization. On the random-interval reinforcement schedule a variable amount of time must elapse after an earned reward before another press will earn a reward. Thus, we focused our organization measures of the time subjects waited in between presses (interpress interval, IPI). As a measure of overall organization, we included the variability/consistency (standard deviation) of the log-transformed interpress intervals. As a measure of short-term patterning and the extent to which current trial behavior relates to past trial behavior, we included the correlation between the current interpress interval, n, and the interpress interval of the prior press, n-1^63,64^. The schematic shows a representative example of these behavioral events across time. **(a-d)** Model predictor variables across training. Data are divided by subjects that were sensitive to devaluation (‘Goal-directed’, devaluation index < group mean of 0.483) v. those that were insensitive (‘Habitual’, devaluation index > 0.483). Lever-press rate (a; 2-way ANOVA, Training: F_1.66, 16.55_ = 11.59, *P* = 0.001; Devaluation sensitivity group: F_1, 10_ = 2.72, *P* = 0.13; Training x Group: F_1.66, 16.55_ = 3.36, *P* = 0.07), behavioral switch rate (b; 2-way ANOVA, Training: F_2.89, 28.89_ = 12.99, *P* < 0.0001; Devaluation sensitivity group: F_1, 10_ = 2.68, *P* = 0.13; Training x Group: F_2.89, 28.89_ = 2.17, *P* = 0.11), variability/consistency (standard deviation) of log interpress intervals (c; 2-way ANOVA, Training: F_2.67, 26.70_ = 12.29, *P* < 0.0001; Devaluation sensitivity group: F_1, 10_ = 0.86, *P* = 0.38; Training x Group: F_2.67, 26.70_ = 0.37, *P* = 0.75), and relationship to past trial interpress interval (n:n-1 interpress interval correlation; d; 2-way ANOVA, Training: F_2.86, 28.61_ = 13.10, *P* < 0.0001; Devaluation sensitivity group: F_1, 10_ = 1.70, *P* = 0.22; Training x Group: F_2.86, 28.61_ = 1.13, *P* = 0.35). Behavioral engagement and transitions tended to increase with training. Interpress-interval variability decreased with training, indicating more consistent overall interpress interval organization. Short-term patterning, the relationship between current trial press wait times and past trial wait, tended to decrease with training, indicating reduced use of past behavior to inform current behavior. These effects did not significantly differ for subjects for which behavior was primarily goal directed (sensitive to devaluation) v. those for which it was primarily habitual (insensitive to devaluation). **(e-f)** Devaluation test press rate (e; 2-way ANOVA, Devaluation sensitivity group x Devaluation: F_1, 10_ = 8.74, *P* = 0.01; Group: F_1, 10_ = 12.18, *P* = 0.006; Devaluation: F_1, 10_ = 13.88, *P* = 0.004 and devaluation index (f; 2-tailed t-test with Welch’s correction, t_6.77_ = 4.09, *P* = 0.005, 95% CI –0.62 – –0.16). **(g-i)** Linear regression model of devaluation index. Using the Matlab stepwiselm function, we identified the model including press rate, switch rate, log interpress-interval variability, and n:n-1 interpress interval correlation that provided the best fit for devaluation index. In preliminary investigations, we also evaluated whether the influence of the interpress interval variables on devaluation sensitivity depended on whether there was a food-port check during the interval and found it did not. The best model (*DevaluationIndex ∼ 1 + StdevlogIPI + IPIn:IPIn-1corr + PressRate*SwitchRate*) provided a strong fit of the data (AIC = –20.59, BIC = –16.12, Log-Likelihood = 17.30, and Deviance = –34.59) and significantly predicted devaluation index (β = 0.70, SE = 0.03; t_8_ = 20.44, *P* < 0.0001, 95% CI 0.62 – 0.77). (g) Devaluation index predicted by the linear regression model (2-tailed t-test with Welch’s correction, t_4.67_ = 4.33, *P* = 0.009, 95% CI – 0.67 – –0.16). (h) Correlation between actual and model-predicted devaluation index (Pearson r_12_ = 0.96, *P* < 0.0001, 95% CI 0.86 – 0.99). (i) Individual variable β coefficients. Neither press rate (β = 0.09, SE = 0.04, t_8_ = 2.06, *P* = 0.07, 95% CI –0.01 – 0.19) nor behavioral switch rate (β = –0.06, SE = 0.04; t_8_ = –1.47, *P* = 0.18, 95% CI –0.16 – 0.03) alone significantly predicted devaluation index, but the interaction between these was a significant negative predictor (β = –0.28, SE = 0.04, t_8_ = –7.86, *P* < 0.0001, 95% CI –0.36 – –0.20), indicating that as switch rate increases, press rate negatively predicts devaluation index. High press rates together with flexibly switching between pressing and checking the food-delivery port was associated with goal-directed control (sensitivity to devaluation). Habit (insensitivity to devaluation was associated by an absence of such press and switch rate coordination. Interpress-interval variability was a significant positive predictor of devaluation index (β = 0.18, SE = 0.03, t_8_ = 7.08, *P* = 0.0001, 95% CI 0.12 – 0.24), indicating that lower variability in the time between presses (low standard deviation of interpress intervals) is associated with goal-directed control (sensitivity to devaluation) and higher inter-press time variability (high standard deviation of interpress intervals) predicted habitual behavioral control (insensitivity to devaluation). Intuitively, habits might be thought to be less variable. However, this result indicates that the extent of goal-directed v. habitual behavioral control is predicted by how strongly the time an animal waits between presses relates to overall organization of waiting. Indeed, the extent to which current trial press wait time relates to past trial behavior was a significant negative predictor of devaluation index (β = –0.06, SEM = 0.02, t_8_ = –2.78, *P* = 0.02, 95% CI –0.10 – –0.01). A stronger relationship between current interpress intervals and past interpress intervals predicted goal-directed behavioral control (sensitivity to devaluation), a weaker relationship was associated with habit (insensitivity to devaluation). **(j-l)** Model performance was evaluated with a 5-fold cross-validation procedure repeated 5 times. Average adjusted R-squared (j) and root mean square error (k) for each cross-validation. Correlation between predicted and actual devaluation index for each model cross validation (l; Pearson R, r_70_ = 0.87, *P* < 0.0001, 95% CI 0.79 – 0.92). D1-cre: *N* = 12 (D1-cre *N* = 4, A2A-cre *N* = 8). Data presented as mean ± s.e.m. Males = solid lines, Females = dashed lines. \*\**P* < 0.05, \*\**P* < 0.01, \*\*\**P* < 0.001. Together this model indicates that devaluation sensitivity (goal-directed behavior) is statistically associated with being on task, engaged in pressing and checking for the outcome and re-engagement in pressing that is planned based on, or at least related to, past performance. Devaluation insensitivity (habit), on the other hand, is statistically associated with less press and switch rate coordination and behavior that is less related to past behavior. Although they are predictive of goal-directed v. habitual control measured by sensitivity to devaluation, these variables linked to movement execution, behavioral engagement, and organization, likely relate to other features of behavior and cannot be taken as indicators of behavioral control mode per se.

**Extended Data Figure 1-5.**
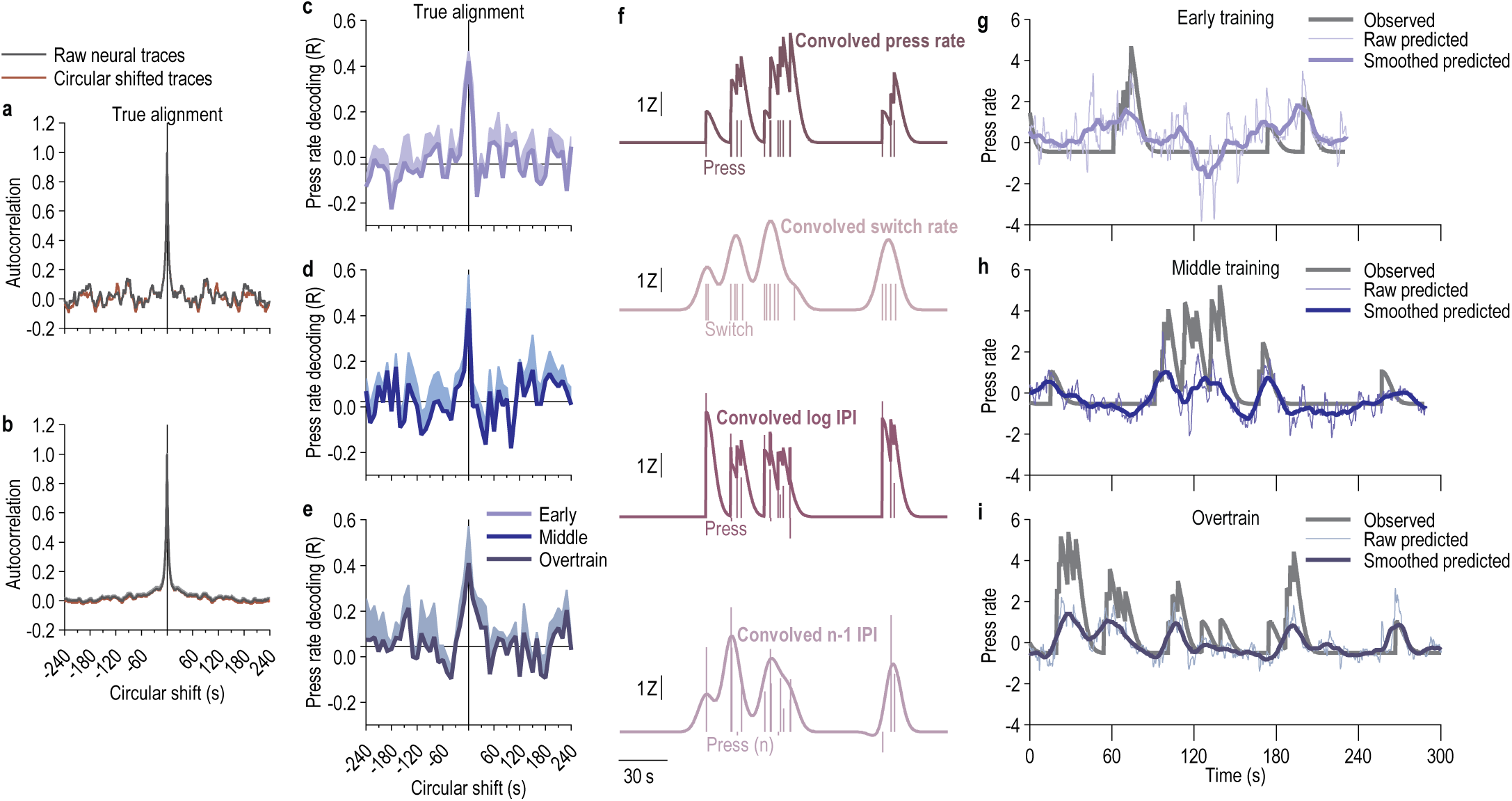
Representative linear decoding. (a-b) Circular shuffle null model preserves population autocorrelation structure. (a) Autocorrelation function of a representative DMS D1^+^ neuron temporal trace compared to its circularly shifted counterpart. (b) DMS D1^+^ neuronal population-averaged autocorrelation function for example subject. **(c-e)** Accuracy of press-rate decoding as a function of time lag. Linear press-rate decoding performance was evaluated as a continuous function of population circular shift magnitude (from –240 s to +240 s in 10-s increments) for the 1^st^ (c, early), 4^th^ (d, middle), and 8^th^ (e, overtrain) training session. Horizonal line indicates average absolute values beyond ±30s. Decoding performance peaks at zero lag and drops off within ±30 s. **(f)** Example convolved press rate, switch rate, current-trial log interpress interval (IPI), and prior trial interpress interval. Vertical lines: time of each event (press, switch, lever press for which the current log or prior interval is calculated), height represents duration of intervals. **(g-i)** Representative example of decoded press rate from the activity of coregistered DMS D1^+^ neurons across training. Actual observed and predicted lever-press rate from the 1^st^ (g, early), 4^th^ (h, middle), and 8^th^ (i, overtrain) training session for a single subject.

**Extended Data Figure 1-6.**
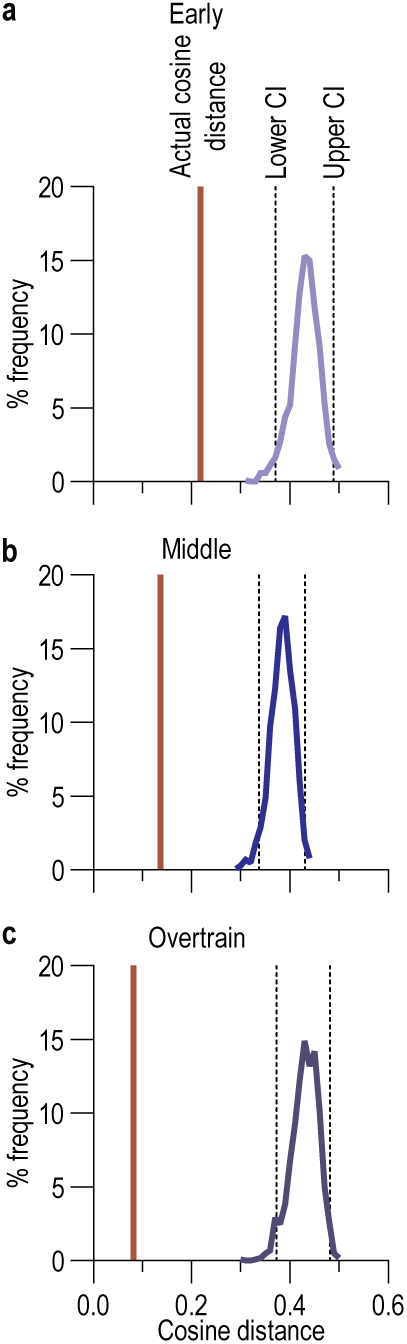
Representative DMS D1^+^ action and reward outcome population vector cosine distance compared to shuffled control. The distribution of cosine distances of a representative subject between action initiation and reward outcome D1^+^ neuronal population vectors for each phase of training (a, early; b, middle; c; overtrain) by bootstrap resampling. The angular distance of the paired group (red line) falls outside of the 95% confidence interval (dashed lines).

**Extended Data Figure 1-7.**
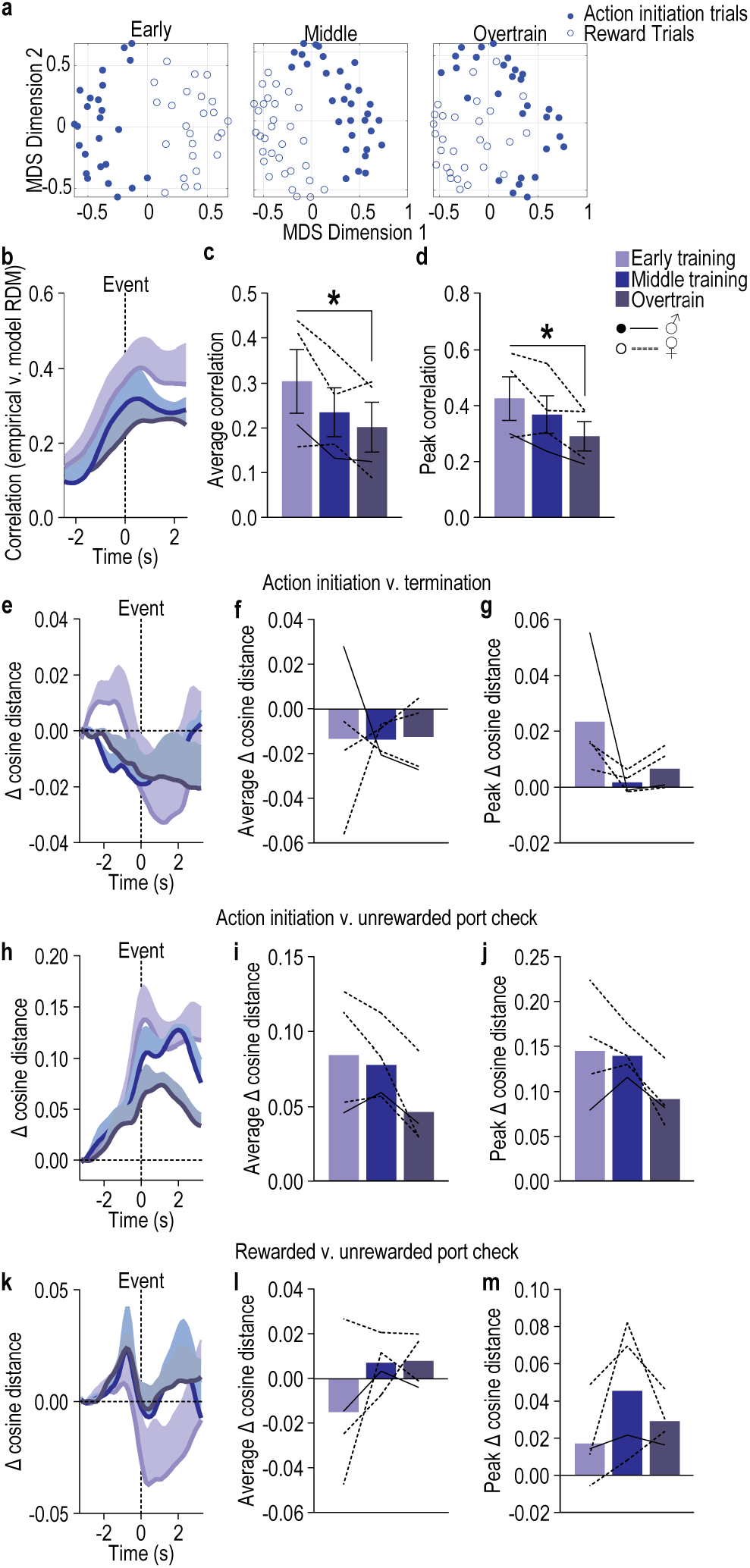
DMS D1^+^ population representation of action and reward becomes more similar with learning. To assess how the neural activity patterns for action initiation and reward evolve over the course of learning, we used a time-resolved representational similarity analysis that allows us to examine the geometry of the neural activity within the entire neural population for each occurrence of action initiation and reward across training sessions. Whereas the population vector analysis considers mean activity across trials, this approach includes trial x trial consistency and, therefore, provides a complementary analysis to the population vector approach. **(a)** Representative single-subject example multidimensional scaling (MDS) plots of the Representational Dissimilarity Matrix (RDM) of pairwise dissimilarities between the DMS D1^+^ population activity patterns around action initiation and earned reward from the 1^st^ (early), 4^th^ (middle), and 8^th^ (overtrain) training session. **(b-d)** Spearman’s rank correlation values between the empirical RDM of the D1^+^ population action initiation and reward activity patterns and those of a theoretical RDM generated to hypothesize complete dissimilarity between action initiation and reward (lower correlation indicates similar neural representations between action initiation and reward) across time centered around each event (b), average (c; 1-way RM ANOVA, F_1.54, 4.63_ = 8.72, *P* = 0.03), or peak (d; 1-way RM ANOVA, F_1.88, 5.64_ = 8.63, *P* = 0.02) ± 2.5 s around the events. With learning and continued training the DMS D1^+^ neuronal population representation of action initiation and earned reward becomes more similar. To evaluate whether the increased representational similarity between action and earned reward is selective to this association or instead reflects a global change in population geometry or task state, we replicated the population vector analysis to examine the geometric relationship between the D1^+^ neuronal population representations of other task events: action initiation and action termination, action initiation and non-reinforced food-delivery port check, reward collection and non-reinforced port check. **(e-g)** Change (Δ) in cosine angular distance, relative to pre-event baseline, between action-initiation and action-termination D1^+^ neuronal population vectors across time and training (e), average (f; 1-way ANOVA, Training: F_1.03, 3.00_ = 0.002, *P* = 0.97), and maximum (g; peak; 1-way ANOVA, Training: F_1.02, 3.05_ = 2.41, *P* = 0.22) ± 2.5 s peri-event. **(h-j)** Change (Δ) in cosine angular distance, relative to pre-event baseline, between the D1^+^ neuronal population vectors of action initiation and non-reward checks of the food-delivery port across time and training (h), average (i; 1-way ANOVA, Training: F_1.06, 3.17_ = 5.94, *P* = 0.09), and maximum (j; peak; 1-way ANOVA, Training: F_1.24, 3.71_ = 5.32, *P* = 0.09) ± 2.5 s peri-event. **(k-m)** Change (Δ) in cosine angular distance, relative to pre-event baseline, between the D1^+^ neuronal population vectors of rewarded and non-reward checks of the food-delivery port across time and training (k), average (l; 1-way ANOVA, Training: F_1.57, 4.71_ = 2.47, *P* = 0.18), and maximum (m; peak; 1-way ANOVA, Training: F_1.40, 4.21_ = 2.57, *P* = 0.19) ± 2.5 s peri-event. Only the population representations of action initiation and earned reward became more similar with training. The population representations of action initiation and termination, action initiation and non-reinforced food-port checks, and rewarded and unrewarded port checks did not become significantly more similar with training [Peak Δ cosine distance; 2-way ANOVA, Comparison type (action initiation:reward, action initiation:action termination, action initiation:non-reinforced check, rewarded check:non-reinforced check) x Training: F_6,18_ = 5.90, *P* = 0.002; Comparison Type: F_3,9_ = 37.68, *P* < 0.0001; Training: F_2,6_ = 20.33, *P* = 0.002; Average Δ cosine distance; 2-way ANOVA, Comparison type x Training: F_6,18_ = 2.54, *P* = 0.05; Comparison Type: F_3,9_ = 31.43, *P* < 0.0001; Training: F_2,6_ = 6.80, *P* = 0.03]. D1-cre: *N* = 4 (1 male). Data presented as mean ± s.e.m. Males = closed circles/solid lines, Females = open circles/dashed lines. \**P* < 0.05.

**Extended Data Figure 2-1.**
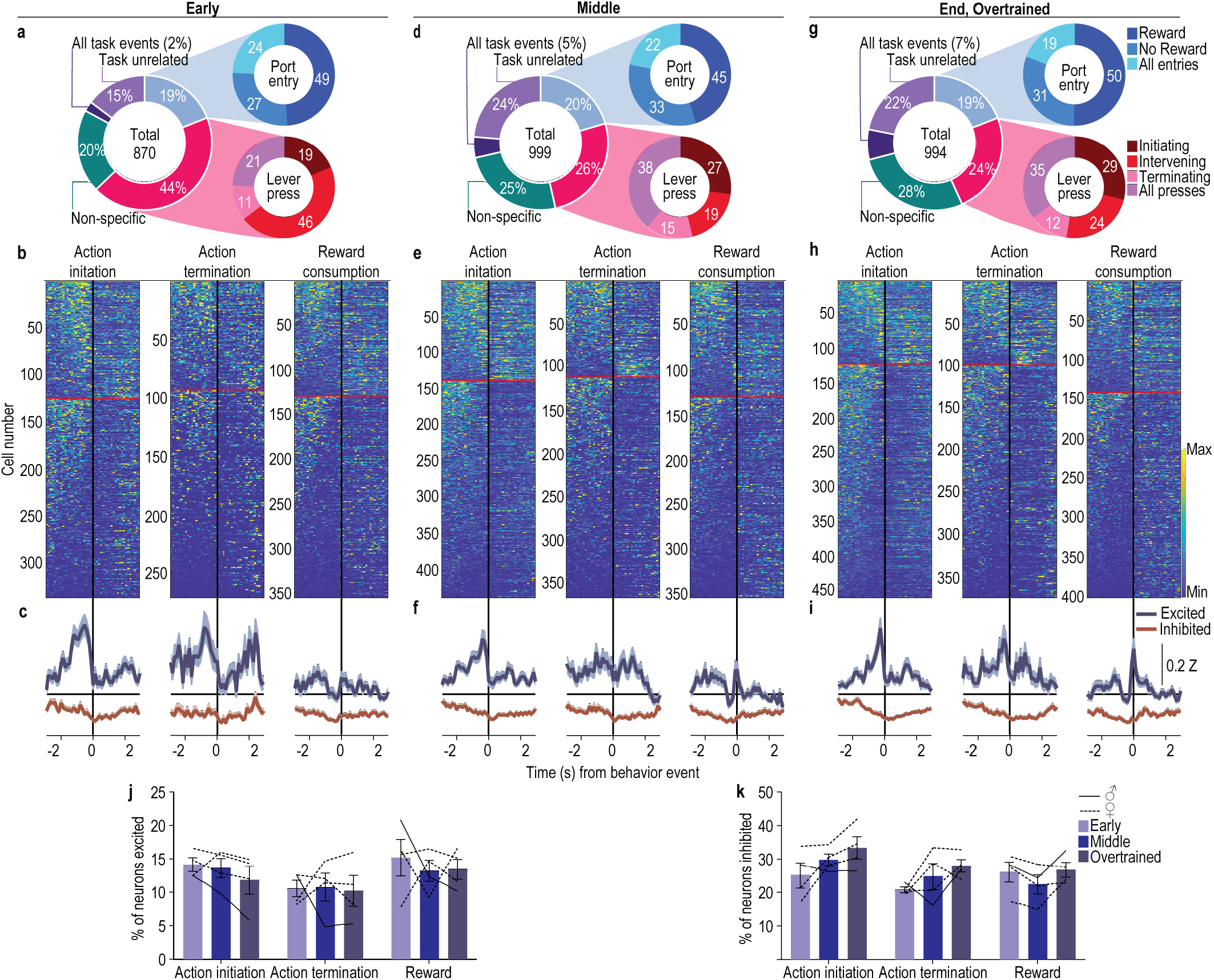
DMS D1^+^ neurons are modulated by actions and rewards during instrumental learning and overtraining. To ask how DMS D1^+^ neurons encode task events during action-outcome learning and the transition to habit, we computed area under the receiver operating characteristic (auROC) values and classified neurons as significantly modulated by critical behavioral events: lever-press action initiation (first press in the session, after earned reward collection, or after a non-reinforced food-port check), action termination (last press before reward collection or non-reinforced food-port check), intervening lever presses (all other presses), non-reinforced food-port checks, and reward collection. 76 – 85% of the neurons significantly increased or decreased their activity within ±2.9 s around one or more of these events. **(a-c)** Activity of DMS D1^+^ neurons on the 1^st^ (Early) session of RI training. (a) Percent of all recorded neurons (*N* = 870) significantly modulated by lever presses, food-delivery port checks, and reward. (b) Heat map of minimum to maximum deconvolved activity (sorted by total activity) of each D1^+^ neuron significantly modulated around lever-press action initiation (left), action termination (middle), or reward consumption (right). Above red line = excited above cutoff criterion, below = inhibited. (c) Z-scored activity of each population of modulated neurons. **(d-f)** Activity of DMS D1^+^ neurons on the 4^th^ (Middle) training session. (d) Percent of all recorded neurons (*N* = 999) significantly modulated by lever presses, food-delivery port checks, and reward. (e) Heat map of each D1^+^ neuron significantly modulated around lever-press action initiation, action termination, or reward consumption. (f) Z-scored activity of each population of modulated neurons. **(g-i)** Activity of DMS D1^+^ neurons on the 8^th^ (End, Overtrain) training session. (g) Percent of all recorded neurons (*N* = 994) significantly modulated by lever presses, food-delivery port checks, and reward. (h) Heat map of each DMS D1^+^ neuron significantly modulated around lever-press action initiation, action termination, or reward consumption. (i) Z-scored activity of each population of modulated neurons. **(j-k)** Percent of all recorded neurons (Early: *N* = 870, Middle: *N* = 999; End *N* = 994) classified (auROC values >95th percentile of the distribution of shuffled auROCs within 2 s before or after event) as significantly excited (j; 2-way ANOVA, Training session: F_1.07, 3.21_ = 0.24, *P* = 0.67; Event: F_1.34, 4.03_ = 2.76, *P* = 0.17; Training x Event: F_1.71, 5.14_ = 0.48, *P* = 0.62) or inhibited (k; –way ANOVA, Training: F_1.75, 5.26_ = 5.34, *P* = 0.06; Event: F_1.47, 4.40_ = 4.32, *P* = 0.10; Training x Event: F_1.66, 4.98_ = 1.56, *P* = 0.29) around initiating lever presses, terminating lever presses, or reward consumption. D1-cre: *N* = 4 (1 male). Data presented as mean ± s.e.m. Males = solid lines, Females = dashed lines. 76 – 85% of DMS D1^+^ neurons significantly increased or decreased their activity within ±2.9 s around one or more of task events. 11 – 14% of DMS D1^+^ neurons were activated around, largely immediately prior to, action initiation. Approximately 10% of neurons were activated around, mostly prior to, action termination. 13 – 15% of neurons responded to the earned reward. Similar proportions of neurons were activated by each task event across training, though there was a slight trend towards more neurons being inhibited around actions with overtraining. Thus, we find that DMS D1^+^ neurons encode actions and outcomes during learning and as habits form with overtraining.

**Extended Data Figure 2-2.**
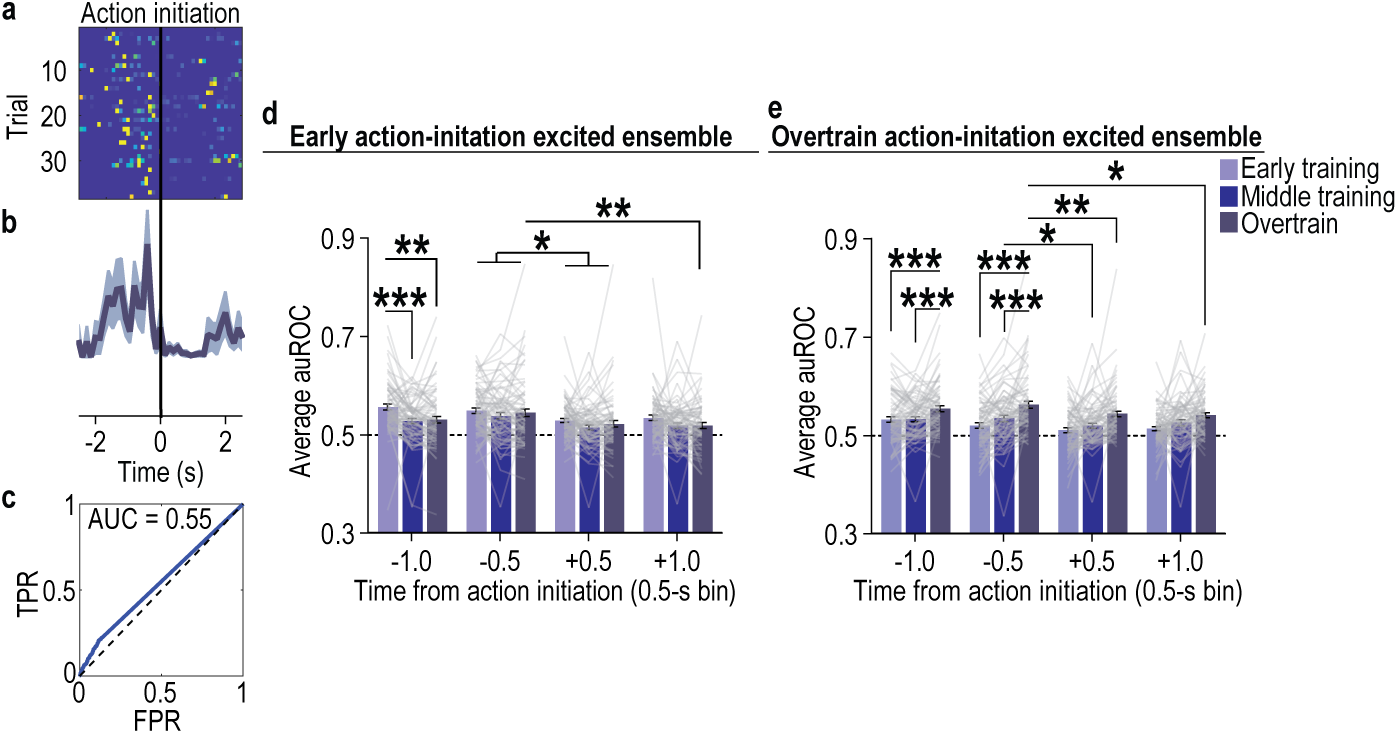
Example and quantification of the modulation of DMS D1^+^ action-initiation excited neurons. (a-c) Representative single-subject/single-session example of the trial x trial activity of a single action-initiation excited DMS D1^+^ neuron. (a) Heat map of minimum to maximum deconvolved activity sorted by trial of a single D1^+^ action-initiation excited DMS neuron. (b) Z-scored, trial-averaged activity of this neuron. (c) ROC plot with true positive rate (TPR) and false positive rate (FPR) for this neuron compared to chance level discrimination (dashed line), with the area under the ROC curve indicated. **(d)** Modulation across training of DMS D1^+^ early action-initiation excited neurons. Modulation index averaged across 0.5-s bins around action initiation. 2-way ANCOVA, Training: F_1.77, 133.28_ = 1.24, *P* = 0.29; Time bin: F_1.79, 133.96_ = 13.96, *P* < 0.001; Training x Time: F_3.40, 254.94_ = 2.94, *P* = 0.028. **(e)** Modulation across training of D1^+^ overtrain action-initiation excited neurons. Modulation index around action initiation. 2-way ANCOVA, Training: F_1.8, 79.90_ = 7.18, *P* = 0.002; Time bin: F_1.64, 65.47_ = 1.50, *P* = 0.23; Training x Time: F_4.28, 171.34_ = 1.50, *P* = 0.20. Data presented as mean ± s.e.m. \**P* < 0.05, \*\**P* < 0.01, \*\*\**P* < 0.001.

**Extended Data Figure 2-3.**
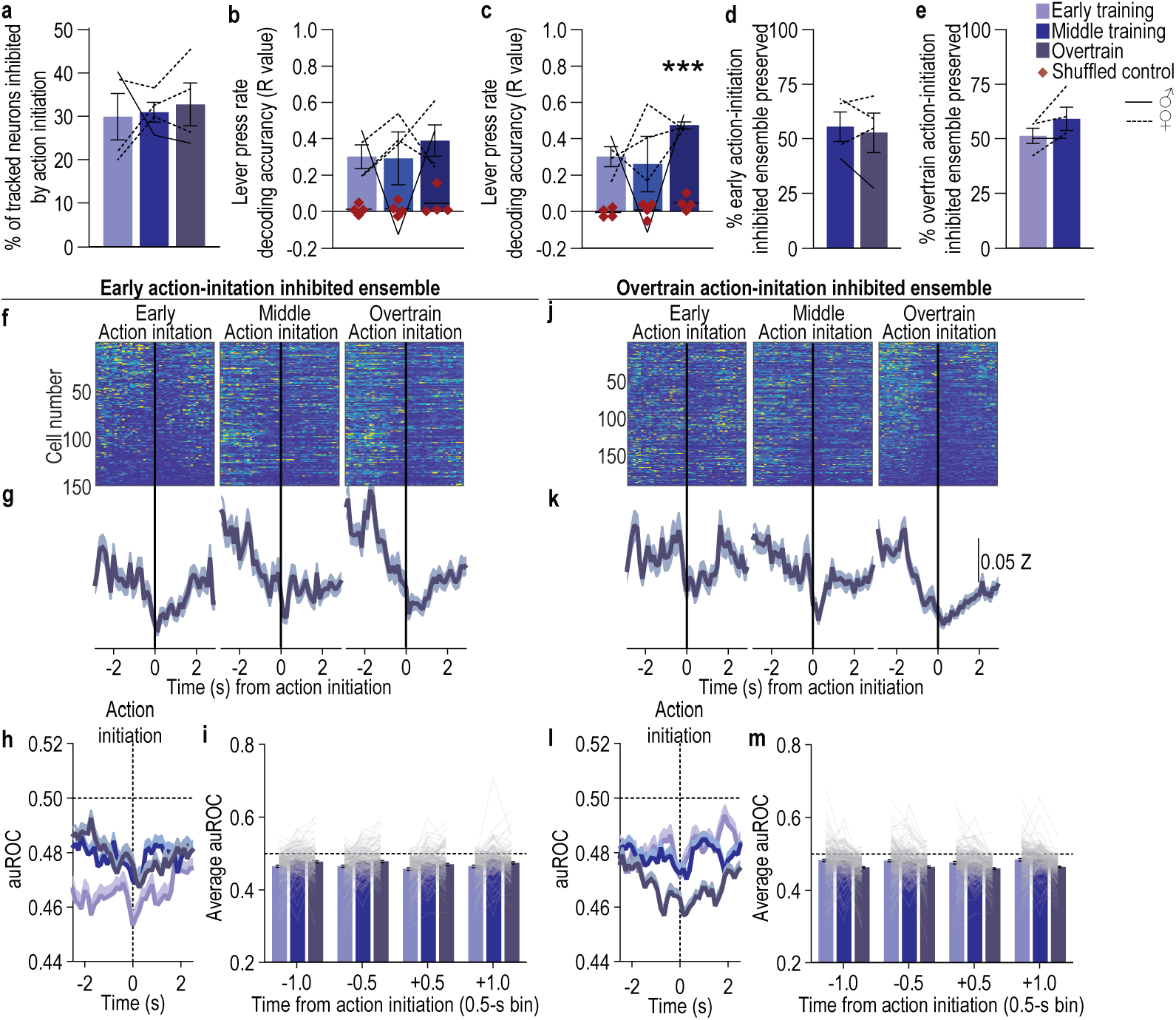
An ensemble of DMS D1^+^ neurons is inhibited during action initiation across learning and as habits form. **(a)** Percent of all recorded coregistered DMS D1^+^ neurons (Average 137 coregistered neurons/mouse, s.e.m. 37.73) significantly inhibited by action initiation. Approximately 30 – 32% of DMS D1^+^ neurons were inhibited around action initiation and this did not change with training. 1-way ANOVA, F_1.29, 3.86_ = 0.14, *P* = 0.79. **(b-c)** Accuracy with which lever-press rate can be can be decoded from the activity of DMS D1^+^ early (b; 2-way ANOVA, Neuron activity: F_1, 3_ = 47.56, *P* = 0.006; Training: F_1.36, 4.07_ = 0.37, *P* = 0.64; Neuron activity x Training: F_1.67, 5.02_ = 0.08, *P* = 0.90) or overtrain (c; 2-way ANOVA, Neuron activity: F_1, 3_ = 84.35, *P* = 0.003; Training: F_1.02, 3.06_ = 1.74, *P* = 0.28; Neuron activity x Training: F_1.02, 3.06_ = 0.57, *P* = 0.51) action-initiation inhibited neurons. Lever-press rate can be reliably decoded from the activity of the population of DMS D1^+^ neurons inhibited by action initiation. **(d)** Percent of D1^+^ early action-initiation inhibited neurons that continued to be significantly inhibited by action initiation on the 4^th^ and 8^th^ training sessions. Approximately 52 – 55% of the early action-initiation inhibited ensemble continued to be inhibited around action initiation during the middle and overtraining phases of training. The proportion preserved did not change with training. 2-tailed t-test, t_3_ = 0.56, *P* = 0.62, 95% CI –19.03 – 13.37. **(e)** Percent of D1^+^ overtrain action-initiation inhibited neurons that were also significantly inhibited by action initiation on 1st and 4th training sessions. Approximately 51 – 59% of the overtraining action-initiation inhibited ensemble was also inhibited around action initiation during the preceding early and middle training phases. The proportion preserved did not change with training. 2-tailed t-test, t_3_ = 2.02, *P* = 0.14, 95% CI –4.52 – 20.15. **(f-i)** Activity and modulation across training of DMS D1^+^ early action-initiation inhibited neurons. Heat map of minimum to maximum deconvolved activity (sorted by total activity) (f), Z-scored activity (g), and area under the receiver operating characteristic curve (auROC) modulation index (h) of these cells around action initiation across training. (i) auROC modulation index averaged across 0.5-s bins around action initiation. 2-way ANCOVA, Training: F_1.65, 246.27_ = 1.46, *P* = 0.24; Time bin: F_2.16, 321.48_ = 2.80, *P* = 0.06; Training x Time: F_4.70, 699.66_ = 0.27, *P* = 0.92. The modulation of the early action-initiation inhibited ensemble did not significantly change with training. **(j-m)** Activity and modulation across training of DMS D1^+^ overtrain action-initiation inhibited neurons. Heat map (h), Z-scored activity (i), and auROC modulation index (j) of these cells around action initiation across training. (m) Modulation index around action initiation. 2-way ANCOVA, Training: F_1.82, 347.95_ = 7.25, *P* = 0.001; Time bin: F_1.91, 365.08_ = 2.37, *P* = 0.10; Training x Time: F_4.35, 831.34_ = 0.67, *P* = 0.62. This overtrain action-initiation inhibited ensemble became more inhibited by action initiation as training progressed. The D1^+^ neuron ensemble that was inhibited around action initiation was stable, similar to the D1^+^ neurons that were excited by action initiation. D1-cre: *N* = 4 (1 male). Data presented as mean ± s.e.m. Males = solid lines, Females = dashed lines.

**Extended Data Figure 2-4.**
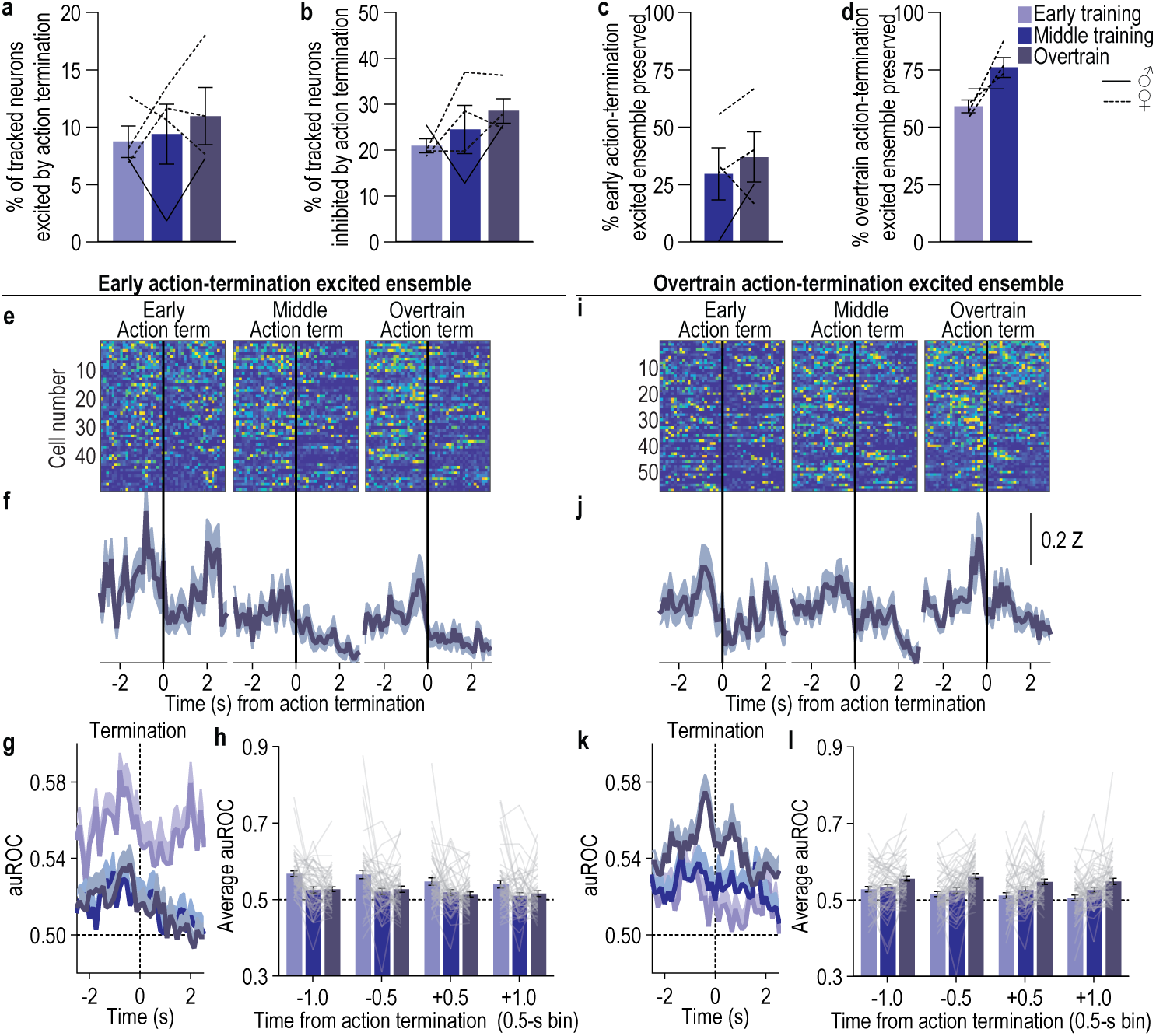
A subensemble of DMS D1^+^ neurons is modulated by action termination during learning and as habits form. (a-b) Percent of all recorded coregistered DMS D1^+^ neurons (Average 137 coregistered neurons/mouse, s.e.m. 37.73) significantly excited (a; 1-way ANOVA, F_1.62, 4.85_ = 0.37, *P* = 0.67) or inhibited (b; 1-way ANOVA, F_1.17, 3.50_ = 1.29, *P* = 0.34) by action termination. Approximately 8 – 10% of coregistered D1^+^ neurons were excited by action termination, and 21 – 29% were inhibited. This did not change with training. **(c)** Percent of DMS D1^+^ early action-termination excited neurons that were then also significantly excited by action termination on the 4^th^ (middle) and 8^th^ (overtrain) training sessions. Approximately 30 – 37% of the early action-termination ensemble continued to be excited by action termination across training. The proportion preserved did not change with training. 2-tailed t-test, t_3_ = 0.85, *P* = 0.46, 95% CI –20.34 – 35.07. **(d)** Percent of DMS D1^+^ overtrain action-termination excited neurons that were also significantly excited by action termination on the 1^st^ (early) and middle training sessions. Approximately 59 – 76% of the overtraining action-termination ensemble was also excited by action termination during the preceding training phases. The proportion preserved did not significantly change with training. 2-tailed t-test, t_3_ = 2.52, *P* = 0.09, 95% CI –4.48 to 38.31. **(e-h)** Activity and modulation across training of DMS D1^+^ early action-termination excited neurons. Heat map of minimum to maximum deconvolved activity (sorted by total activity) (d), Z-scored activity (e), and area under the receiver operating characteristic curve (auROC) modulation index (f) of these neurons around action termination across training. (h) auROC modulation index averaged across 0.5-s bins around action termination. 2-way ANCOVA, Training: F_1.92, 97.81_ = 1.08, *P* = 0.34; Time bin: F_1.65, 84.19_ = 0.48, *P* = 0.59; Training x Time: F_4.20, 214.10_ = 1.33, *P* = 0.26. Modulation of this early action-termination ensemble did not significantly change with training. **(i-l)** Activity and modulation across training of DMS D1^+^ overtrain action-termination excited neurons. Heat map (h), Z-scored activity (i), and auROC modulation index (j) of these cells around action termination across training. (l) Modulation index around action termination. 2-way ANOVA, Training: F_1.77, 99.05_ = 1.14, *P* = 0.32; Time bin: F_1.78, 99.88_ = 0.03, *P* = 0.96; Training x Time: F_4.20, 235.08_ = 0.71, *P* = 0.60. Modulation of the overtrain action-termination ensemble did not significantly change as training progressed. D1-cre: *N* = 4 (1 male). Data presented as mean ± s.e.m. Males = solid lines, Females = dashed lines.

**Extended Data Figure 2-5.**
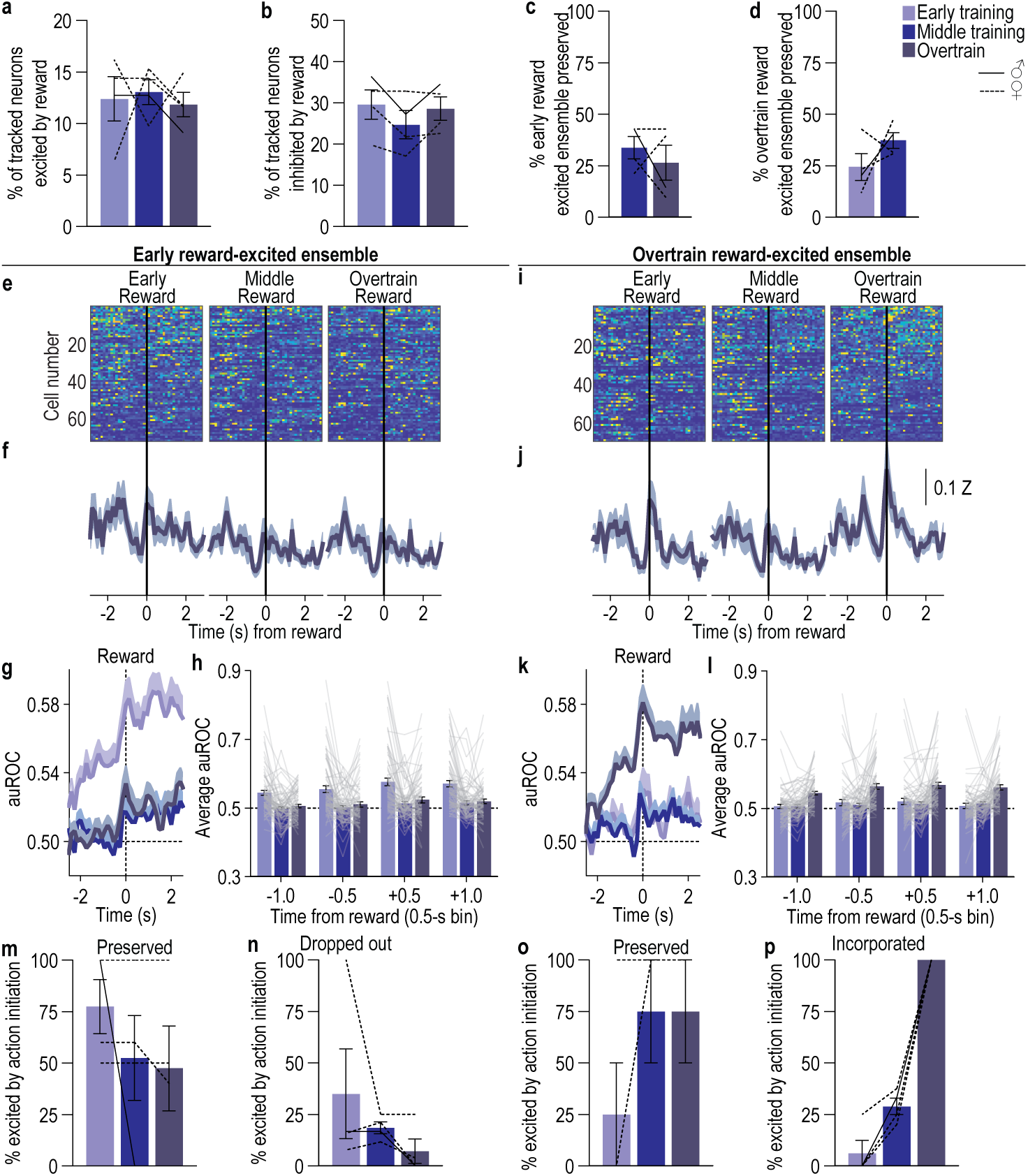
An ensemble of DMS D1^+^ neurons is modulated by reward during learning and as habits form. (a-b) Percent of all recorded coregistered DMS D1^+^ neurons (Average 137 coregistered neurons/mouse, s.e.m. 37.73) significantly excited (a; 1-way ANOVA, F_1.46, 4.38_ = 0.12, *P* = 0.83) or inhibited (b; 1-way ANOVA, F_1.91, 5.76_ = 2.52, *P* = 0.16) by earned reward. Approximately 12 – 13% of coregistered D1^+^ neurons were excited by reward and 25 – 30% were inhibited. This did not change with training. **(c)** Percent of D1^+^ early reward excited neurons that continued to be significantly excited by reward during the middle and overtraining phases. Approximately 26 – 34% of the early reward ensemble continued to be excited by reward across training. The proportion preserved did not change with training. 2-tailed t-test, t_3_ = 0.70, *P* = 0.53, 95% CI –40.49 – 25.89. **(d)** Percent of DMS D1^+^ overtrain reward excited neurons that were significantly excited by reward on the early and middle training sessions. Approximately 24 – 37% of the overtraining reward ensemble was also excited by reward during the preceding training phases. The proportion preserved did not change with training. 2-tailed t-test, t_3_ = 1.31, *P* = 0.28, 95% CI –18.55 – 44.33. **(e-h)** Activity and modulation across training of DMS D1^+^ early reward excited neurons. Heat map (e), Z-scored activity (f), and auROC modulation index (g) of these cells around reward across training. (h) Modulation index around reward. 2-way ANCOVA, Training: F_1.74, 123.26_ = 0.82, *P* = 0.42; Time bin: F_2.45, 175.73_ = 6.07, *P* = 0.01; Training x Time: F_4.82, 342.52_ = 1.59, *P* = 0.16. This early reward ensemble was more excited after earned reward than before, suggesting modulation by reward experience. This did not significantly change as training progressed. **(i-l)** Activity and modulation across training of D1^+^ overtrain reward excited neurons. Heat map (i), Z-scored activity (j), and auROC modulation index (k) of these cells around reward across training. (l) Modulation index around reward. 2-way ANCOVA Training: F_1.77, 120.87_ = 2.26, *P* = 0.12; Time bin: F_2.36, 160.42_ = 1.30, *P* = 0.28; Training x Time: F_5.37, 365.26_ = 0.28, *P* = 0.93. Modulation of the overtrain reward ensemble did not significantly change with training. **(m)** Percentage of the preserved early reward-excited neurons (those that continued to encode reward across training) that were also significantly excited by action initiation. 1-way ANOVA, F_1.06, 3.18_ = 1.27, *P* = 0.34. A substantial proportion of the neurons that continue to encode reward across training also encode action initiation. **(n)** Percentage of the neurons that dropped out of the early reward-excited ensemble (those that stopped being modulated by reward with training) that were significantly excited by action initiation. 1-way ANOVA, F_1.04, 3.13_ = 1.78, *P* = 0.27. Neurons that dropped out of the reward ensemble tended to be less likely to also encode action initiation. **(o)** Percentage of the preserved overtrain reward-excited neurons (those that encoded reward across training) that were also significantly excited by action initiation. 1-way ANOVA, F_1.00, 3.00_ = 3.00, *P* = 0.18. A substantial proportion of the neurons that encoded reward across training also encode action initiation. **(p)** Percentage of the neurons that were incorporated into the overtrain reward-excited ensemble (those that became modulated by reward with training) that were significantly excited by action initiation. 1-way ANOVA, F_1.48, 4.44_ = 194.3, *P* < 0.0001. Strikingly, all of the neurons that picked up encoding of the reward with training also encoded action initiation. Those neurons that either stably encode or pick up encoding of reward are also those that encode action initiation. Whereas, those neurons that stop encoding the reward were less likely to encode action initiation. Consistent with the emergent similarity of the population representation of action and outcome, this data suggests that with training individual neurons encode both the instrumental action and its reward outcome. D1-cre: *N* = 4 (1 male). Data presented as mean ± s.e.m. Males = solid lines, Females = dashed lines.

**Extended Data Figure 2-6.**
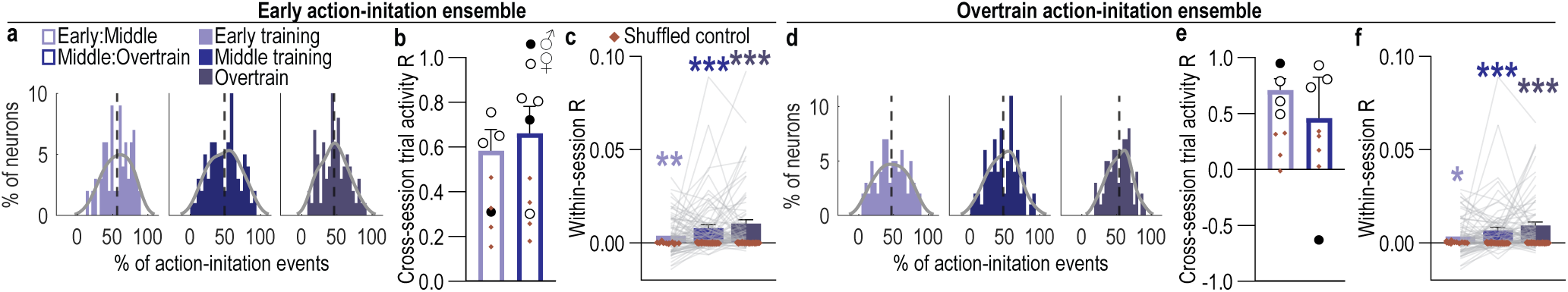
DMS D1^+^ neurons encode action initiation with high fidelity. (a-c) Fidelity with which D1^+^ early action-initiation excited neurons encode action initiation. **(a)** Distribution of the percentage of early action-initiation excited neurons as a function of the percentage of action-initiation events to which they respond for each training phase. **(b)** Cross-session correlation of the response distributions. 2-way ANOVA, Neuron activity distribution (v. shuffled): F_1, 3_ = 16.06, *P* = 0.03; Training session: F_1, 3_ = 0.25, *P* = 0.65; Distribution x Training: F_1, 3_ = 0.11, *P* = 0.76. Early action-initiation D1^+^ neurons tend to respond on more than half the action-initiation events across training and this is consistent across training. **(c)** Within-session correlation of the activity around action initiation of each early action-initiation excited neuron. 2-way ANCOVA, Neuron activity: F_1, 74_ = 27.35, *P* < 0.001; Training: F_2, 148_ = 5.84, *P* = 0.004; Activity x Time: F_2, 148_ = 6.28, *P* = 0.002. The activity of early action-initiation excited D1^+^ neurons around action initiation is correlated above shuffled control within a training session and becomes more correlated with training. **(d-e)** Fidelity with which D1^+^ overtrain action-initiation excited neurons encode action initiation. **(d)** Distribution of the percentage of overtrain action-initiation excited neurons as a function of the percentage of action-initiation events to which they respond for each training phase. **(e)** Cross-session correlation of the response distributions. 2-way ANOVA, Neuron activity distribution: F_1, 3_ = 19.36, *P* = 0.02; Training: F_1, 3_ = 0.27, *P* = 0.64; Distribution x Training: F_1, 3_ = 0.35, *P* = 0.60. Overtrain action-initiation D1^+^ neurons tend to respond on more than half the action-initiation events across training and this is consistent across training. **(f)** Within-session correlation of the activity around action initiation of each overtrain action-initiation excited neuron. 2-way ANCOVA, Neuron activity: F_1, 73_ = 19.32, *P* < 0.001; Training: F_2, 146_ = 1.87, *P* = 0.16; Activity x Time: F_2, 146_ = 1.96, *P* = 0.15. The activity of overtrain action-initiation excited D1^+^ neurons around action initiation is correlated above shuffled control within a training session and becomes more correlated with training. D1-cre: *N* = 4 (1 male). Data presented as mean ± s.e.m. Males = closed circles, Females = open circles.

**Extended Data Figure 2-7.**
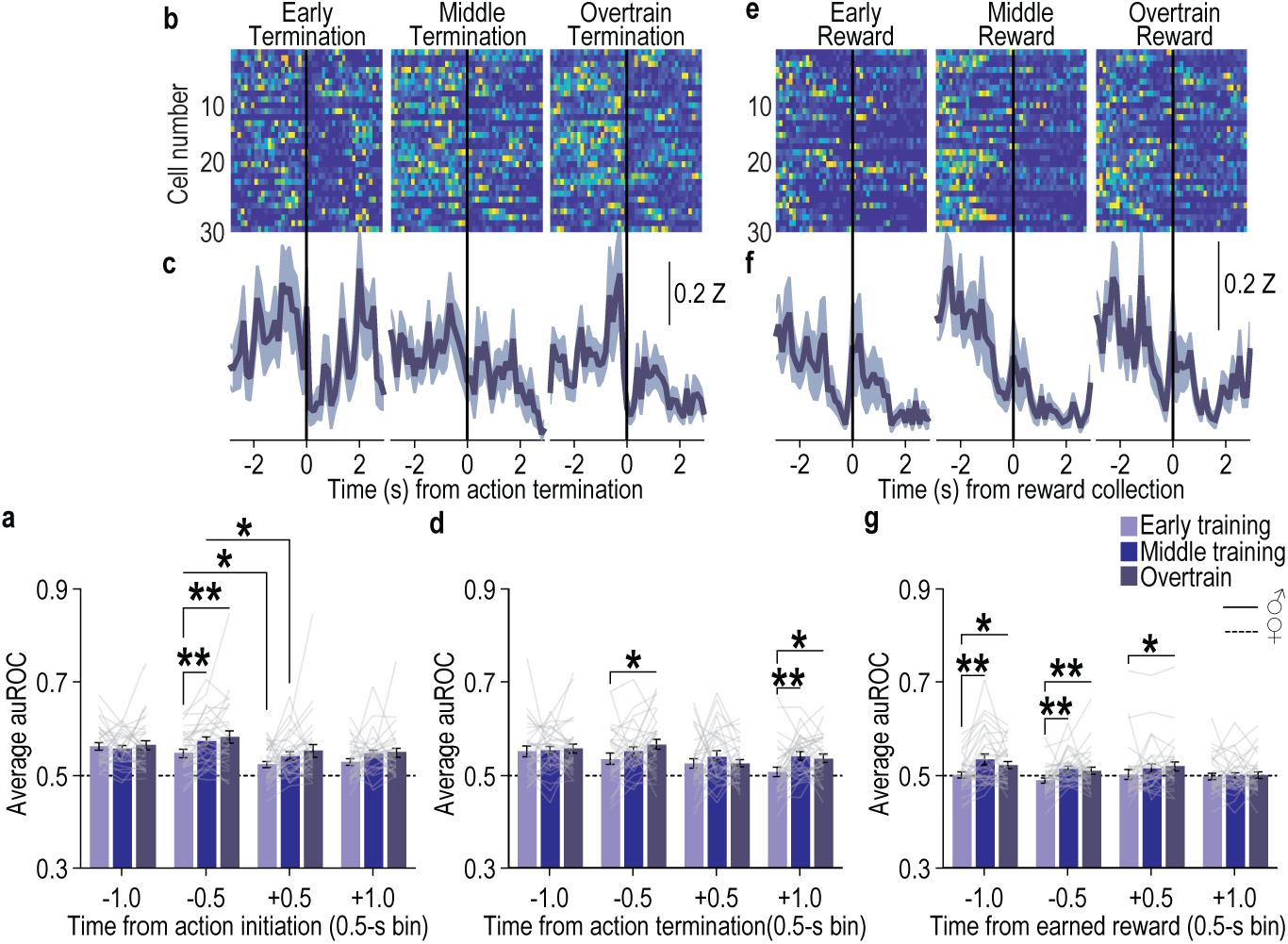
A subensemble of stable DMS D1^+^ action-initiation excited neurons develop encoding of action termination and earned reward with training. We identified an ensemble (*N =* 31 neurons/4 D1-cre mice, 1 male; average 7.75 neurons/mouse, s.e.m. = 2.56) of D1^+^ neurons that stably encoded action-initiation across all phases of training. **(a)** Modulation, averaged in 0.5-s bins around action initiation of these neurons. 2-way ANCOVA, Training: F_1.70, 49.21_ = 0.04, *P* = 0.94; Time bin: F_1.73, 50.27_ = 5.02, *P* = 0.01; Training x Time: F_2.28, 123.97_ = 2.78, *P* = 0.03. Stable action-initiation excited D1^+^ neurons are more activated prior to action initiation than after and this modulation improves with training. **(b-c)** Heat map of minimum to maximum deconvolved activity (sorted by total activity) (b) and Z-scored (c) activity around action termination across training of the D1^+^ stable action-initiation excited neurons. **(d)** Modulation, averaged in 0.5-s bins around action termination of these neurons. 2-way ANCOVA, Training: F_1.87, 54.10_ = 1.01, *P* = 0.37; Time bin: F_2.00, 57.95_ = 1.48, *P* = 0.23; Training x Time: F_3.62, 104.97_ = 1.98, *P* = 0.11. **(e-f)** Heat map (e) and Z-scored (f) activity around reward collection across training of D1^+^ stable action-initiation excited neurons. **(g)** Modulation, averaged in 0.5-s bins around earned reward of these neurons. 2-way ANCOVA, Training: F_1.99, 57.57_ = 2.61, *P* = 0.08; Time bin: F_1.84, 53.36_ = 0.80, *P* = 0.45; Training x Time: F_3.87, 112.10_ = 1.74, *P* = 0.15. Data presented as mean ± s.e.m. Males = solid lines, Females = dashed lines.

**Extended Data Figure 3-1:**
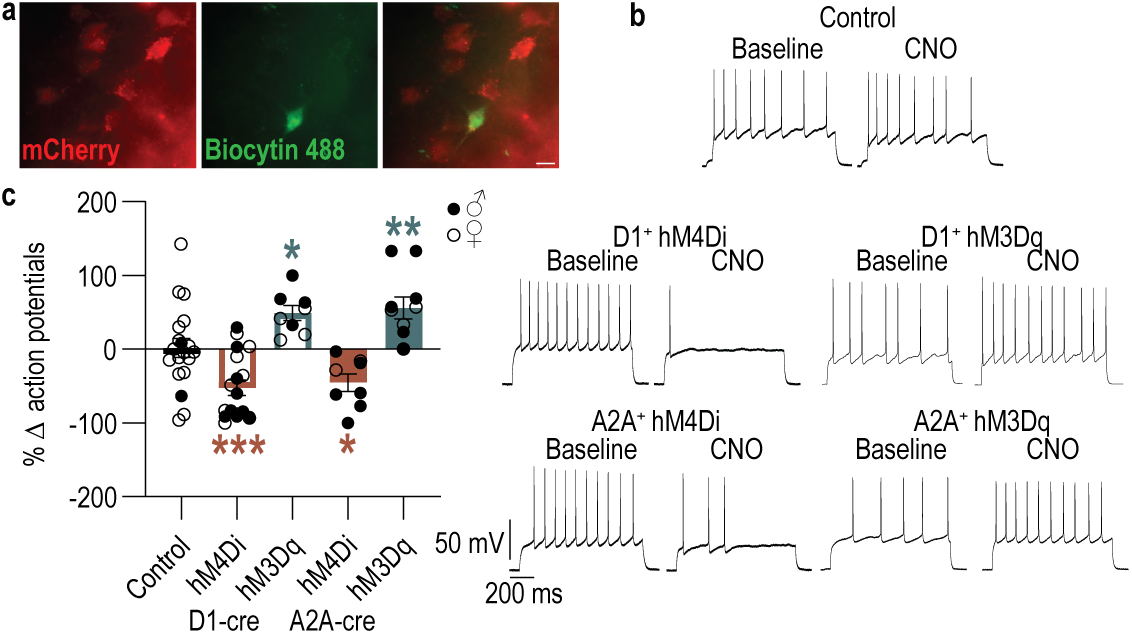
Validation of chemogenetic approach to inhibit or activate DMS D1^+^ or A2A^+^ neurons. Fluorescence-guided, whole-cell patch clamp recordings in current-clamp mode were used to validate the efficacy of chemogenetic manipulation of DMS D1^+^ and A2A^+^ neurons. **(a)** Representative immunofluorescent image of a biocytin-filled, mCherry-positive cell. **(b)** Representative recordings of action potentials in single cells before (Baseline) and after CNO (1 µM for hM3Dq and 100 µM^128,129^ for hM4Di) bath application. Current injection intensity applied to induce action potential firing was 200 (Control, D1^+^ hM3Dq, A2A^+^ hM3Dq) or 250 pA (all others), with the same intensity used for baseline and CNO recordings. Cell membrane potentials were held at –80 mV using negative current to ensure consistent baseline conditions across recordings. **(c)** Percent change in action potentials under CNO, relative to pre-CNO baseline. 1-way ANOVA, F_4,58_ = 13.29, *P* < 0.0001. Control, *N* = 19 cells, 4 D1-cre mice (1 male); D1-cre, hm4Di: *N* = 8 cells, 4 mice (3 males), hM3Dq: *N* = 10 cells, 5 mice (3 males); A2A-cre, hM4Di: *N* = 8 cells, 4 mice (3 males), hM3Dq: *N* = 10 cells, 5 mice (3 males). Data presented as mean ± s.e.m. Males = closed circles, Females = open circles. Scale bar = 10 µm. \**P* < 0.05, \*\**P* < 0.01, \*\*\**P* < 0.001. Under these ex vivo conditions, CNO altered excitability in the expected direction. It is important to note that the CNO concentration used for inhibition is high and might have had off-target effects. Lower concentrations were not tested.

**Extended Data Figure 3-2.**
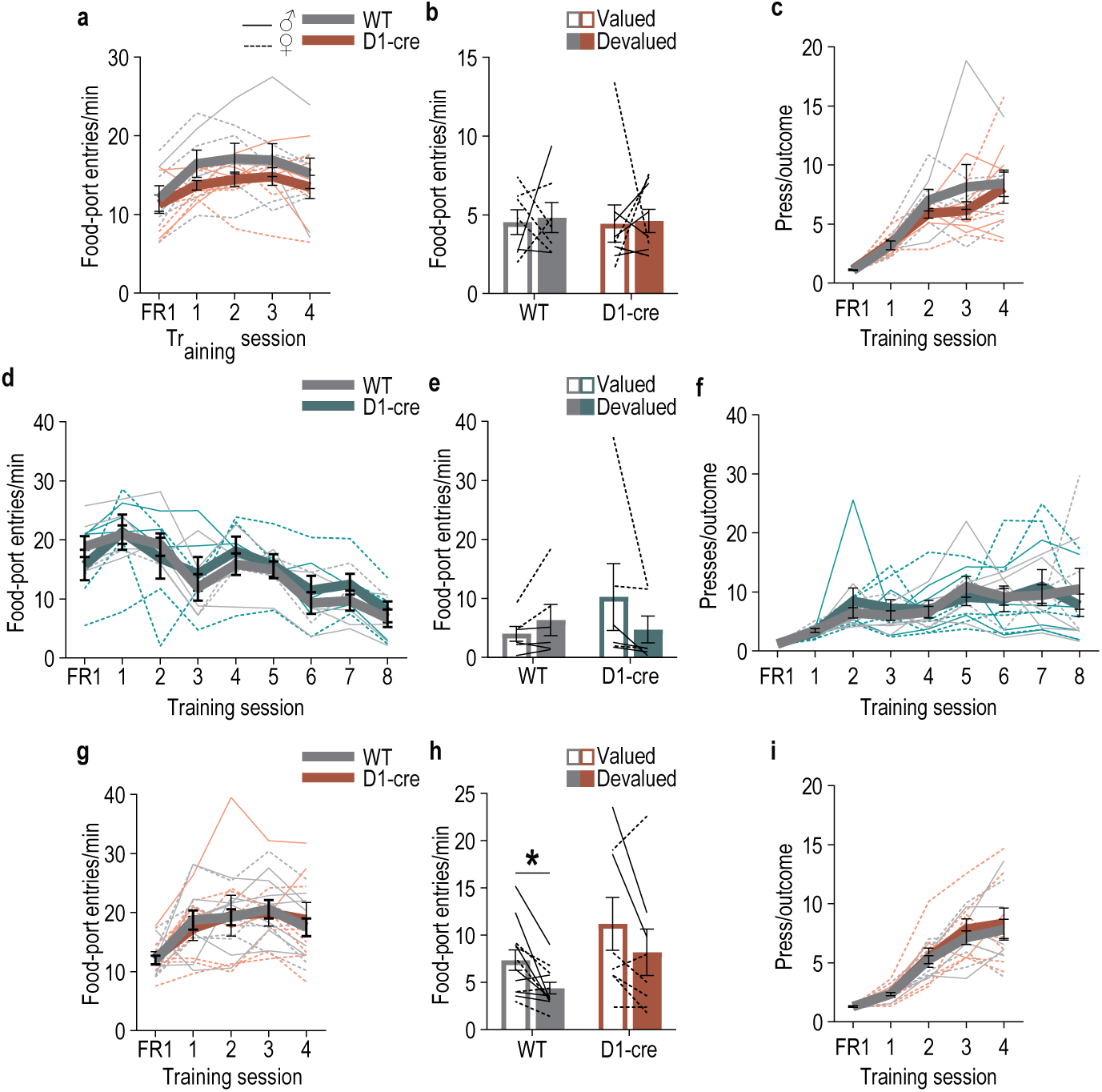
Food-port entries during training and test with DMS D1^+^ neuron chemogenetic manipulation. (a-c) Chemogenetic inactivation of DMS D1^+^ neurons during learning. WT: *N* = 7 (2 males); D1-cre: *N* = 9 (5 males). **(a)** Training entry rate. 2-way ANOVA, Training: F_2.35, 32.91_ = 7.40, *P* = 0.001; Genotype: F_1,14_ = 1.30, *P* = 0.27; Training x Genotype: F_4, 56_ = 0.46, *P* = 0.77. **(b)** Test entry rate. 2-way ANOVA, Value x Genotype: F_1, 14_ = 0.003, *P* = 0.96; Value: F_1, 14_ = 0.05, *P* = 0.83; Genotype: F_1, 14_ = 0.03, *P* = 0.86. **(c)** Training average presses/earned reward outcome. 2-way ANOVA, Training: F_2.05, 28.68_ = 32.52, *P* < 0.0001; Genotype: F_1,14_ = 0.57, *P* = 0.46; Training x Genotype: F_4, 56_ = 0.77, *P* = 0.55. **(d-f)** Chemogenetic activation of DMS D1^+^ neurons during learning. WT: *N* = 8 (5 males); D1-cre: *N* = 9 (4 males). **(d)** Training entry rate. 2-way ANOVA, Training: F_2.38, 23.79_ = 12.75, *P* < 0.0001; Genotype: F_1, 10_ = 0. 21, *P* = 0.74; Training x Genotype: F_8, 80_ = 0.74, *P* = 0.65. **(d)** Test entry rate. 2-way ANOVA, Value x Genotype: F_1, 10_ < 0.0001, *P* > 0.99; Value: F_1, 10_ = 0.89, *P* = 0.37; Genotype: F_1, 10_ = 0.51, *P* = 0.49. **(f)** Training average presses/earned reward outcome. 2-way ANOVA, Training: F_4.03, 60.47_ = 9.42, *P* < 0.0001; Genotype: F_1, 15_ = 0.0001, *P* = 0.99; Training x Genotype: F_8, 120_= 0.60, *P* = 0.78. **(g-i)** Chemogenetic inactivation of DMS D1^+^ neurons at test after learning. WT: *N* = 12 (7 males); D1-cre: *N* = 8 (2 males). **(g)** Training entry rate. 2-way ANOVA, Training: F_3.01, 54.19_ = 13.85, *P* < 0.0001; Genotype: F_1, 18_ < 0.0001, *P* > 0.99; Training x Genotype: F_4, 72_ = 0.34, *P* = 0.85. **(h)** Test entry rate. 2-way ANOVA, Value x Genotype: F_1, 18_ = 0.001, *P* = 0.97; Value: F_1, 18_ = 11.24, *P* = 0.004; Genotype: F_1, 18_ = 3.02, *P* = 0.10. **(i)** Training average presses/earned reward outcome. 2-way ANOVA, Training: F_1.86, 44.59_ = 96.52, *P* < 0.0001; Genotype: F_2, 24_= 0.13, *P* = 0.88; Training x Genotype: F_8, 96_ = 0.13, *P* > 0.99. Data presented as mean ± s.e.m. Males = solid lines, Females = dashed lines. \**P* < 0.05. Neither chemogenetic inhibition nor activation of DMS D1^+^ neurons affected checks of the food-delivery port or altered the press-reward action-outcome relationship.

**Extended Data Figure 3-3.**
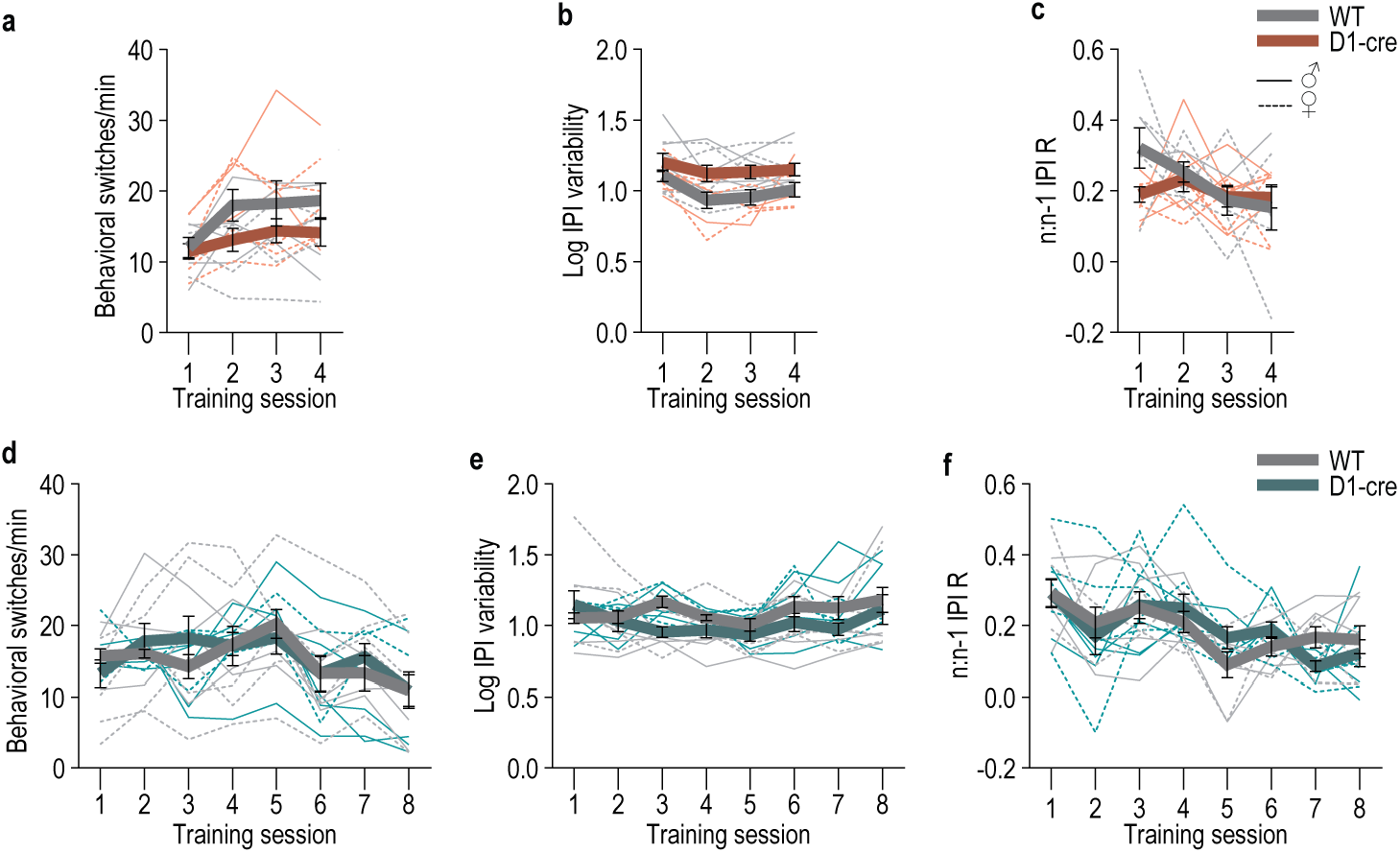
Behavioral engagement and organization variables during training with DMS D1^+^ neuron chemogenetic manipulation. (a-c) Chemogenetic inactivation of DMS D1^+^ neurons during learning. WT: *N* = 7 (2 males); D1-cre: *N* = 9 (5 males). Behavioral switch rate (a; how often subjects stop one behavior to transition to another i.e., press ◊ food-port check or food-port check ◊ press; 2-way ANOVA, Training: F_2.25, 31.49_ = 5.74, *P* = 0.006; Genotype: F_1, 14_ = 2.25, *P* = 0.16; Training x Genotype: F_2.25, 31.49_ = 1.16, *P* = 0.33), interpress interval variability (b; standard deviation of the log-transformed interpress-intervals; 2-way ANOVA, Genotype: F_1, 14_ = 7.64, *P* = 0.02; Training: F_2.38, 33.33_ = 3.05, *P* = 0.05; Training x Genotype: F_2.38, 33.33_ = 0.43, *P* = 0.68), and relationship to past trial behavior (c; correlation between the current interpress interval (n) and the interpress interval of the prior press (n-1); 2-way ANOVA, Genotype: F_1, 14_ = 0.77, *P* = 0.39; Training: F_2.62, 36.68_ = 3.04, *P* = 0.05; Training x Genotype: F_2.62, 36.68_ = 1.93, *P* = 0.15) across days of training. Consistent the evidence from the linear regression model of the variables associated with habit, inhibition of DMS D1^+^ neurons caused slightly lower behavioral switch rate and significantly lower consistency (higher interpress interval variability). It did not significantly affect how much current trial interpress wait time related to past trial wait times. **(d-f)** Chemogenetic activation of DMS D1^+^ neurons during learning. WT: *N* = 8 (5 males); D1-cre: *N* = 9 (4 males). Behavioral switch rate (d; 2-way ANOVA, Training: F_2.98, 44.64_ = 5.53, *P* = 0.003; Genotype: F_1, 15_ = 0.03, *P* = 0.87; Training x Genotype: F_2.98, 44.64_ = 0.94, *P* = 0.43), interpress interval variability (e; Training: F_2.21, 33.10_ = 2.08, *P* = 0.13; Genotype: F_1, 15_ = 1.76, *P* = 0.21; Training x Genotype: F_2.21, 33.10_ = 1.43, *P* = 0.25), and relationship to past trial behavior (f; 2-way ANOVA, Genotype: F_1, 15_ = 0.003, *P* = 0. 69; Training: F_2.21, 33.10_ = 2.08, *P* = 0.14; Training x Genotype: F_2.21, 33.10_ = 1.43, *P* = 0.25) across days of training. Excitation of DMS D1^+^ neurons did not affect behavioral engagement and organizational variables. Data presented as mean ± s.e.m. Males = solid lines, Females = dashed lines.

**Extended Data Figure 4-1:**
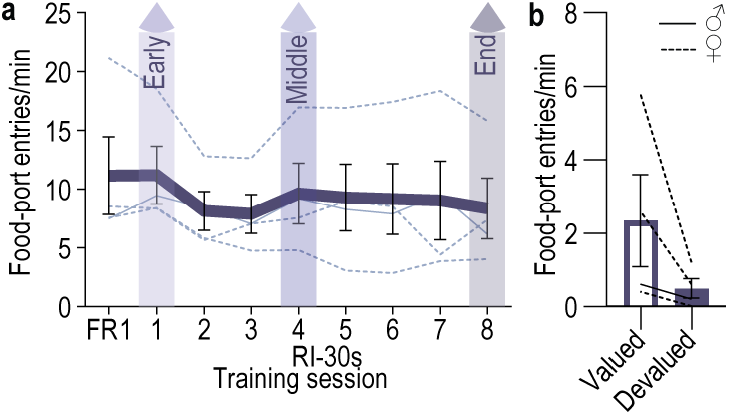
Entries into the food-delivery port during training and test for DMS A2A^+^ imaging experiment-habitual subjects. **(a)** Training entry rate. 1-way ANOVA, Training: F_2.69, 8.06_ = 1.93, *P* = 0.20 **(b)** Test entry rate. 2-tailed t-test, t_3_ = 1.87, *P* = 0.16, 95% CI –5.00 – 1.30. Data presented as mean ± s.e.m. Males = solid lines, Females = dashed lines.

**Extended Data Figure 4-2.**
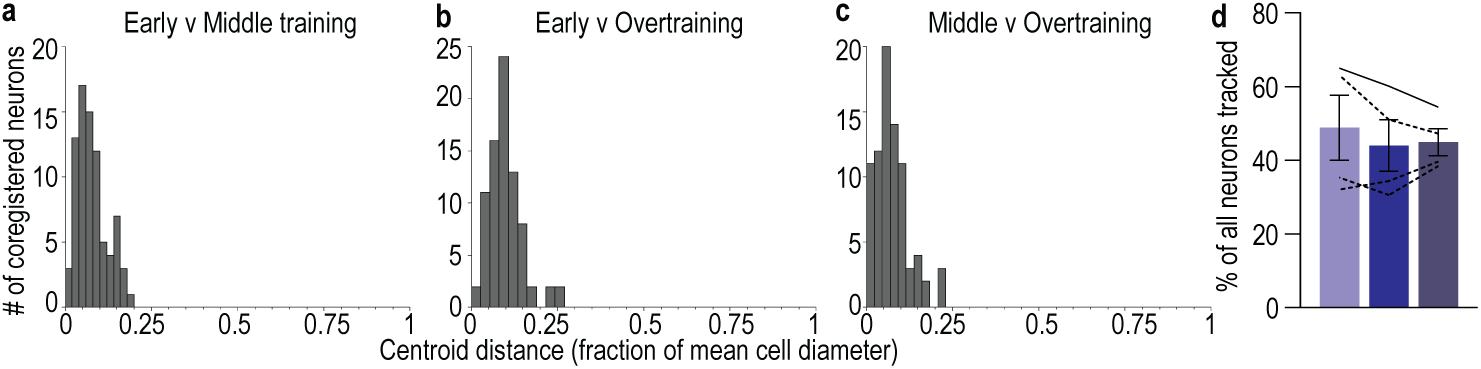
Representative example of coregistration of DMS A2A^+^ neurons across training. **(a)** Distribution of the distance between cell centroids (as a fraction of total cell diameter) of coregistered cell pairs in the 1^st^ (early) and 4^th^ (middle) training sessions. **(b)** Distribution of centroid distance of coregistered cell pairs in the 1^st^ and 8^th^ (overtrain) training sessions. **(c)** Distribution of centroid distance of coregistered cell pairs in the 4th and 8^th^ training sessions. **(d)** Percent of all neurons coregistered across training. 1-way ANOVA, F_1.18, 3.54_ = 0.78, *P* = 0.45. A2A-cre: *N* = 4 (1 male). We were able to coregister on average 47% (s.e.m. 3.60%) of the A2A^+^ neurons across all three training phases. Data presented as mean ± s.e.m. Males = solid lines, Females = dashed lines.

**Extended Data Figure 4-3.**
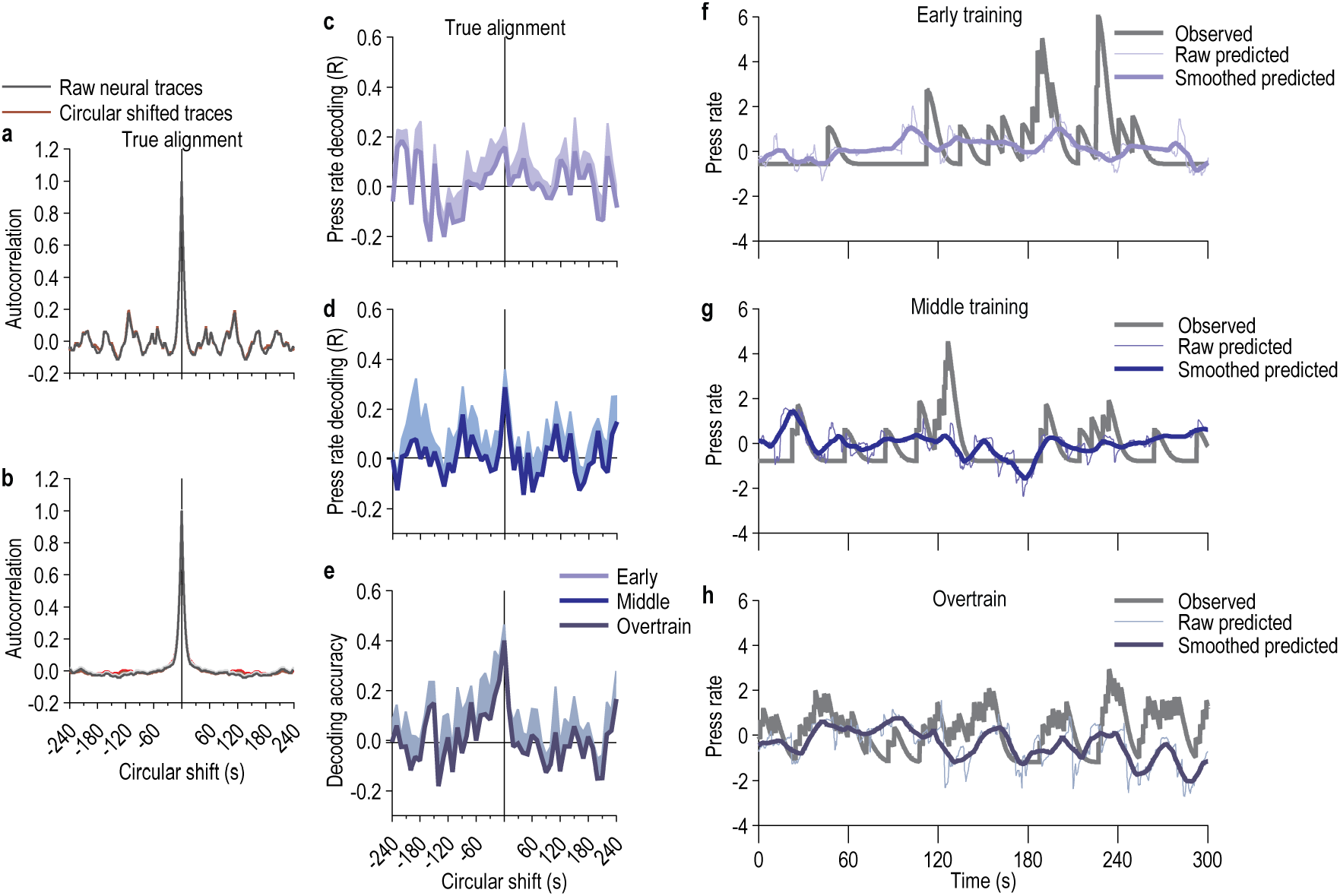
Representative decoded press rate from the activity of coregistered DMS A2A^+^ neurons across training. (a-b) Circular shuffle null model preserves population autocorrelation structure. (a) Autocorrelation function of a representative DMS A2A^+^ neuron temporal trace compared to its circularly shifted counterpart. (b) DMS A2A^+^ neuronal population-averaged autocorrelation function for example subject. **(c-e)** Accuracy of press-rate decoding as a function of time lag. Linear press-rate decoding performance was evaluated as a continuous function of population circular shift magnitude (from –240 s to +240 s in 10-s increments) for the 1^st^ (c, early), 4^th^ (d, middle), and 8^th^ (e, overtrain) training session. Horizonal line indicates average absolute values beyond ±30s. Decoding performance peaks at zero lag and drops off within ±30 s. **(f-h)** Actual observed and predicted lever-press rate from the 1^st^ (f, early), 4^th^ (g, middle), and 8^th^ (h, overtrain) training session for a single subject.

**Extended Data Figure 4-4.**
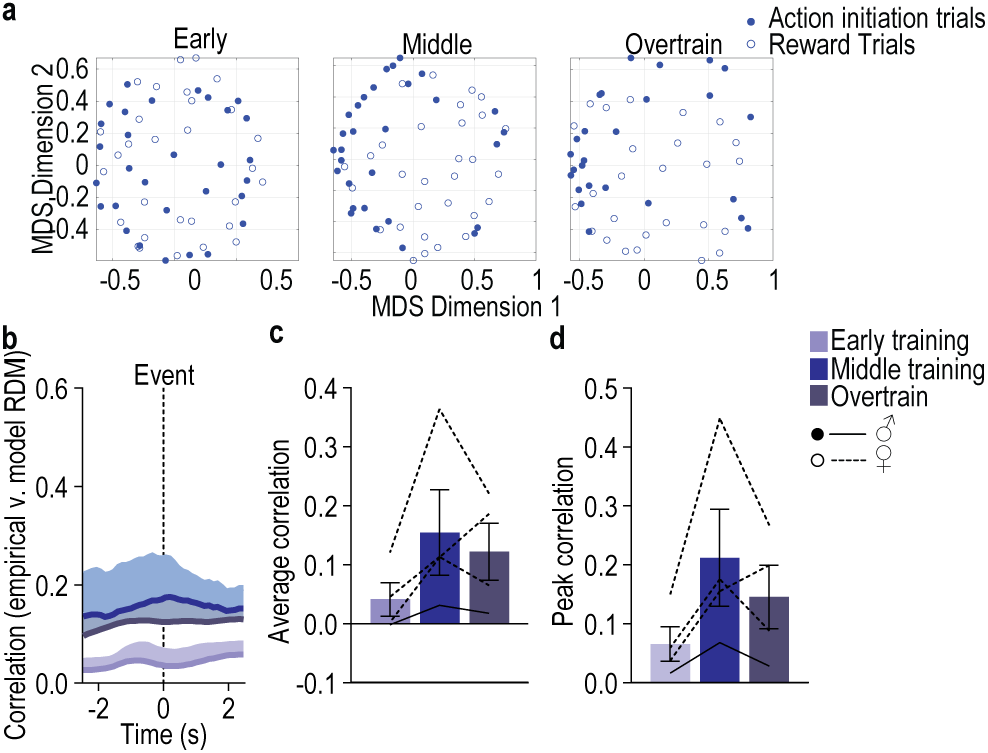
DMS A2A^+^ population representation of action and reward does not become more similar with learning. We used a time-resolved Representational Similarity analysis to assess how the neural activity patterns for action initiation and reward evolve over the course of learning. **(a)** Representative single-subject example multidimensional scaling (MDS) plots of the Representational Dissimilarity Matrix (RDM) of pairwise dissimilarities between the DMS A2A^+^ population activity patterns around action initiation and earned reward from the 1^st^ (early), 4^th^ (middle), and 8^th^ (overtrain) training session. **(b-d)** Spearman’s rank correlation values between the empirical RDM of the A2A^+^ population action initiation and reward activity patterns and those of a theoretical RDM generated to hypothesize complete dissimilarity between action initiation and reward (lower correlation indicates similar neural representations between action initiation and reward) across time centered around each event (b), average (c; 1-way RM ANOVA, F_1.93, 5.08_ = 3.62, *P* = 0.10), or peak (d; 1-way RM ANOVA, F_1.69, 5.07_ = 5.06, *P* = 0.06) ± 2.5 s around the events. With learning and continued training the DMS A2A^+^ neuronal population representation of action initiation and earned reward does not significantly change, but trends towards becoming less similar. A2A-cre: *N* = 4 (1 male). Data presented as mean ± s.e.m. Males = closed circles/solid lines, Females = open circles/dashed lines.

**Extended Data Figure 5-1.**
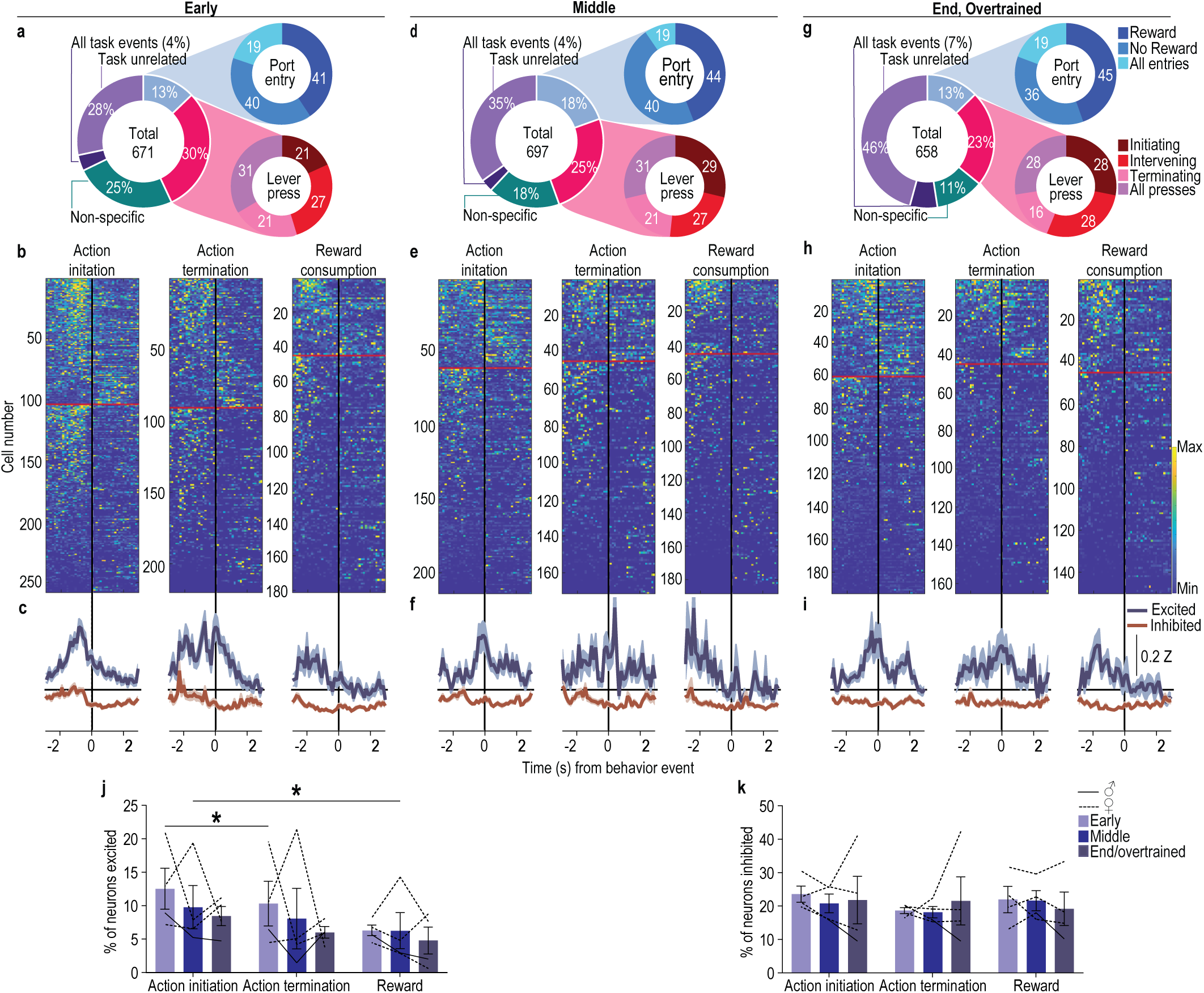
DMS A2A^+^ neurons are modulated by actions during instrumental learning and overtraining. To ask how DMS A2A^+^ neurons encode task events during action-outcome learning and the transition to habit, we computed auROC values and classified neurons as significantly modulated by critical behavioral events: lever-press action initiation (first press in the session, after earned reward collection, or after a non-reinforced food-port check), action termination (last press before reward collection or non-reinforced food-port check), intervening lever presses (all other presses), non-reinforced food-port checks, and reward collection. **(a-c)** Activity of DMS A2A^+^ neurons on the 1^st^ (Early) session of RI training. (a) Percent of all recorded neurons (*N* = 671) significantly modulated by lever presses, food-delivery port checks, and reward. (b) Heat map of minimum to maximum deconvolved activity (sorted by total activity) of each DMS A2A^+^ neuron significantly modulated around lever-press action initiation (left), action termination (middle), or reward consumption (right). Above red line = excited, below = inhibited. (c) Z-scored activity of each population of modulated neurons. **(d-f)** Activity of DMS A2A^+^ neurons on the 4^th^ (Middle) training session. (d) Percent of all recorded neurons (*N* = 697) significantly modulated by lever presses, food-delivery port checks, and reward. (e) Heat map of minimum to maximum deconvolved activity of each DMS A2A^+^ neuron significantly modulated around lever-press action initiation, action termination, or reward. (f) Z-scored activity of each population of modulated neurons. **(g-i)** Activity of DMS A2A^+^ neurons on the 8^th^ (End, Overtrain) training session. (g) Percent of all recorded neurons (*N* = 658) significantly modulated by lever presses, food-delivery port checks, and reward. (h) Heat map of minimum to maximum deconvolved activity of each DMS A2A^+^ neuron significantly modulated around lever-press action initiation, action termination, or reward. (i) Z-scored activity of each population of modulated neurons. **(j-k)** Percent of all recorded neurons (Early: *N* = 671, Middle: *N* = 697; End *N* = 658) classified (auROC values >95th percentile of the distribution of shuffled auROCs within 2 s before or after event) as significantly excited (j; 2-way ANOVA, Event: F_1.12, 3.36_ = 23.83, *P* = 0.01; Training session: F_1.32, 3.97_ = 0.05, *P* = 0.54; Training x Event: F_1.86, 5.59_ = 0.29, *P* = 0.77) or inhibited (k; 2-way ANOVA, Event: F_1.90, 5.69_ = 2.03, *P* = 0.23; Training session: F_1.12, 3.36_ = 0.05, *P* = 0.86; Training x Event: F_1.45, 4.33_ = 0.90, *P* = 0.44) around initiating lever presses, terminating lever presses, or reward consumption. Action initiation excited more DMS A2A^+^ neurons than termination or reward. Similar proportions of the DMS A2A^+^ neurons were inhibited around each event type. Data presented as mean ± s.e.m. Males = solid lines, Females = dashed lines. \**P* < 0.05.

**Extended Data Figure 5-2.**
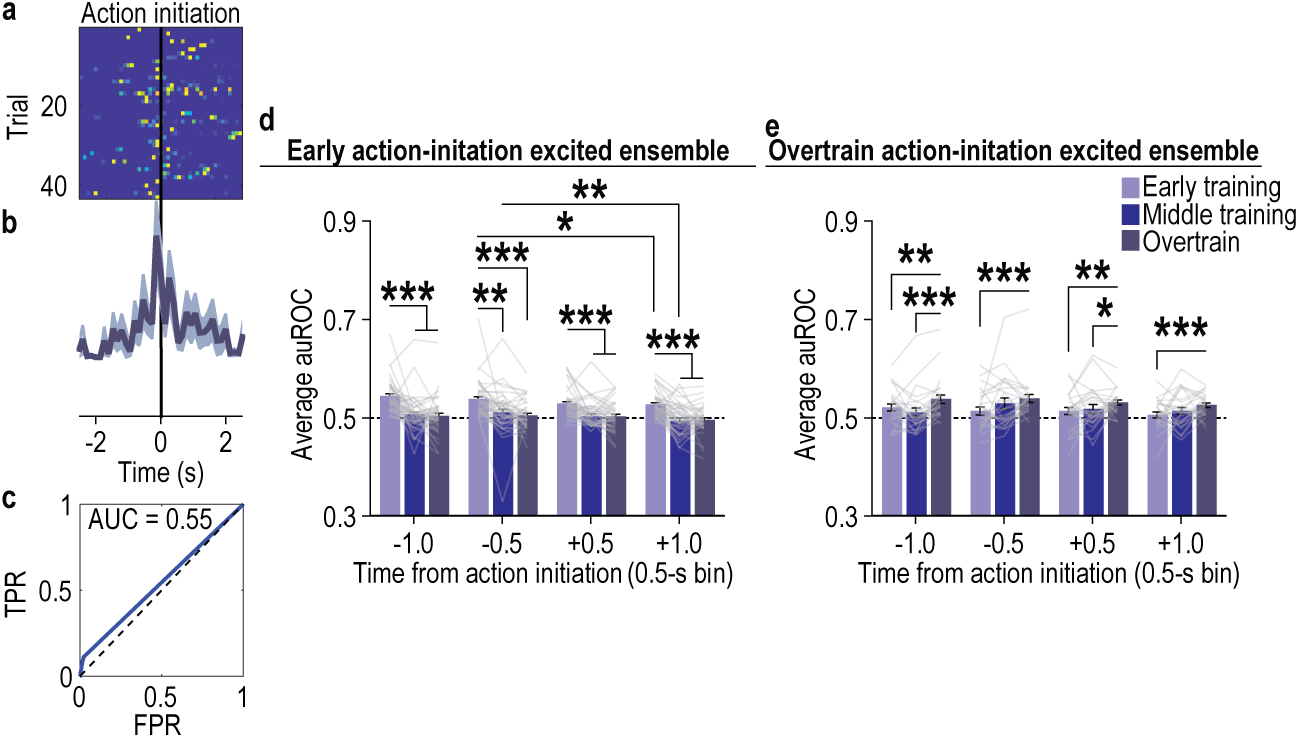
Example and quantification of the modulation of DMS A2A^+^ action-initiation excited neurons. (a-c) Representative single-subject/single-session example of the trial x trial activity of a single action-initiation excited DMS A2A^+^ neuron. (a) Heat map of minimum to maximum deconvolved activity sorted by trial of a single A2A^+^ action-initiation excited neuron. (b) Z-scored, trial-averaged activity of this neuron. (c) ROC plot with true positive rate (TPR) and false positive rate (FPR) for this neuron compared to chance level discrimination (dashed line), with the area under the ROC curve indicated. **(d)** Modulation across training of DMS A2A^+^ early action-initiation excited neurons. Modulation index averaged across 0.5-s bins around action initiation. 2-way ANCOVA, Training: F_1.74, 69.82_ = 4.10, *P* = 0.03; Time bin: F_1.70, 68.14_ = 0.18, *P* = 0.80; Training x Time: F_4.70, 187.84_ = 0.90, *P* = 0.68. **(e)** Modulation across training of A2A^+^ overtrain action-initiation excited neurons. Modulation index around action initiation. 2-way ANCOVA, Training: F_1.75, 40.22_ = 1.15, *P* = 0.32; Time bin: F_1.74, 39.90_ = 0.10, *P* = 0.88; Training x Time: F_4.23, 97.45_ = 0.64, *P* = 0.70. Data presented as mean ± s.e.m. \**P* < 0.05, \*\**P* < 0.01, \*\*\**P* < 0.001.

**Extended Data Figure 5-3.**
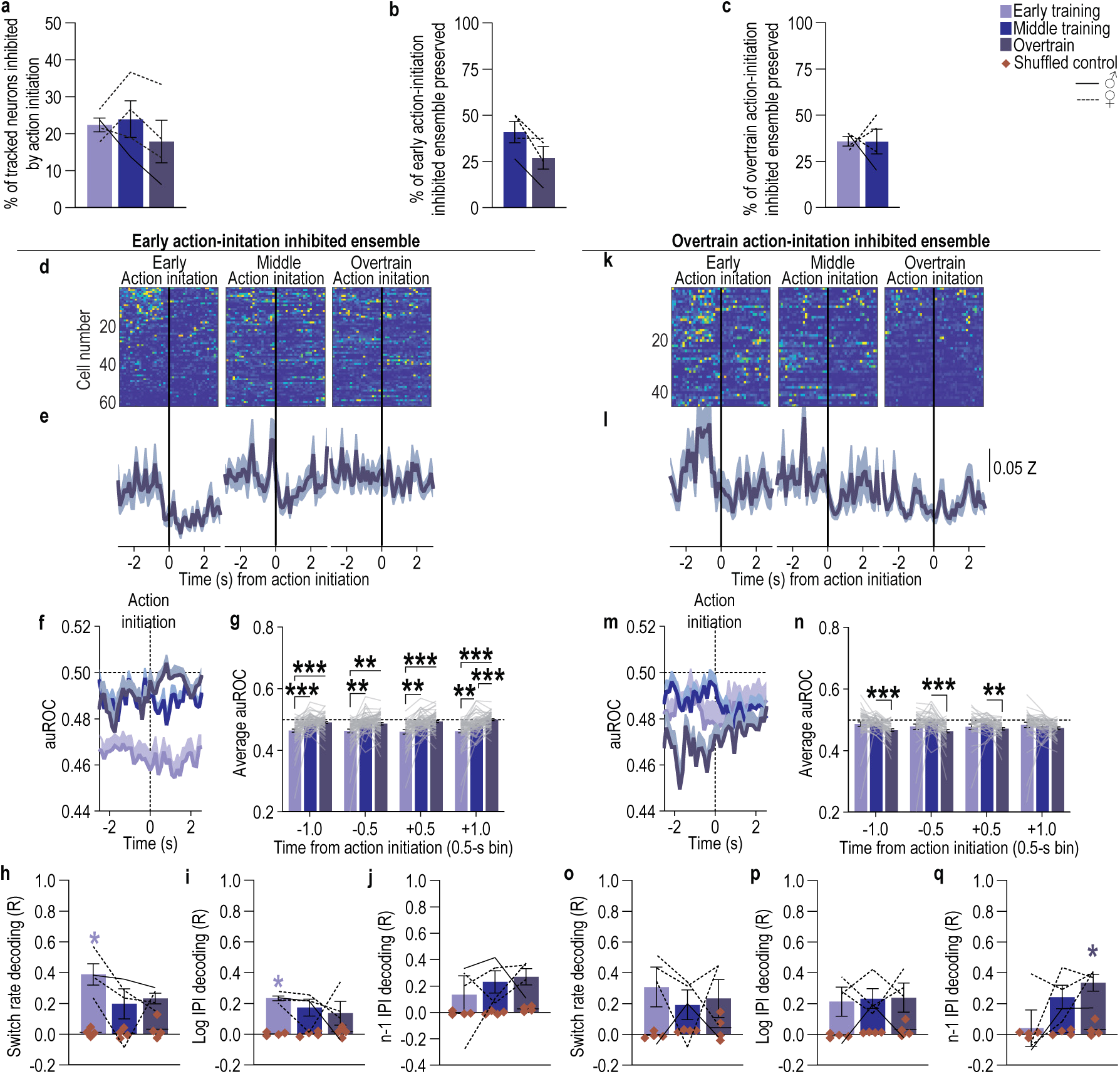
The ensemble of DMS A2A^+^ neurons inhibited by action initiation shifts as habits form. **(a)** Percent of all recorded coregistered DMS A2A^+^ neurons (Average 73.25 coregistered neurons/mouse, s.e.m. 17.09) significantly inhibited by action initiation. Approximately 18 – 24% of coregistered DMS A2A^+^ neurons were inhibited by action initiation and this did not significantly change across training. 1-way ANOVA, F_1.05, 3.16_ = 1.14, *P* = 0.37. **(b)** Percent of DMS A2A^+^ early action-initiation-inhibited neurons that were then also significantly inhibited by action initiation on the 4^th^ (middle) and 8^th^ (overtrain) training sessions. Approximately 27 – 41% of the early action-initiation inhibited ensemble continued to be inhibited around action initiation during the middle and overtraining phases of training. The proportion preserved did not change with training. 2-tailed t-test, t_3_ = 2.70, *P* = 0.07, 95% CI –30.41 – 2.52. **(c)** Percent of DMS A2A^+^ overtrain action-initiation-inhibited neurons that were significantly modulated by action initiation on prior the 1^st^ (early) and 4^th^ (middle) training sessions. Approximately 36% of the overtraining action-initiation inhibited ensemble was also inhibited around action initiation during the preceding early and middle training phases. The proportion preserved did not change with training. 2-tailed t-test, t_3_ = 0.01, *P* = 0.99, 95% CI –29.03 – 28.80. **(d-f)** Activity and modulation across training of DMS A2A^+^ early action-initiation-inhibited neurons. Heat map of minimum to maximum deconvolved activity (sorted by total activity) (d), Z-scored activity (e), and area under the receiver operating characteristic curve (auROC) modulation index (f) of these cells around action initiation across training. **(g)** auROC modulation index averaged across 0.5-s bins around action initiation. 2-way ANCOVA, Training: F_1.38, 83.91_ = 1.38, *P* = 0.26; Time bin: F_2.48, 151.16_ = 3.28, *P* = 0.02; Training x Time: F_5.10, 311.30_ = 3.80, *P* = 0.001. This early action-initiation inhibited ensemble became less inhibited by action initiation as training progressed. **(h-j)** Linear decoding accuracy from DMS A2A^+^ early action-initiation inhibited neuron activity behavioral switch rate (h; 2-way ANOVA, Neuron activity: F_1, 3_ = 24.52, *P* = 0.01; Training: F_1.62, 4.85_ = 3.48, *P* = 0.12; Neuron activity x Training: F_1.37, 4.09_ = 1.54, *P* = 0.30), log interpress intervals (i; 2-way ANOVA, Neuron activity: F_1, 3_ = 448.40, *P* = 0.0002; Training: F_1.03, 3.10_ = 0.46, *P* = 0.55; Neuron activity x Training: F_1.04, 3.12_ = 0.55, *P* = 0.48), and the prior press interpress interval (j; 2-way ANOVA, Neuron activity: F_1, 3_ = 13.53, *P* = 0.03; Training: F_1.45, 4.36_ = 0.61, *P* = 0.54; Neuron activity x Training: F_1.49, 4.48_ = 0.29, *P* = 0.70). Behavioral switch rate, relative interpress interval, and prior press interpress interval, could be decoded from the activity of the early action-initiation inhibited A2A^+^ neurons. **(k-m)** Activity and modulation across training of coregistered DMS A2A^+^ overtrain action-initiation-inhibited neurons. Heat map of minimum to maximum deconvolved activity (k), Z-scored activity (l), and area under the receiver operating characteristic curve (auROC) modulation index (m) of these cells around action initiation across training. **(n)** auROC modulation index averaged across 0.5-s bins around action initiation. 2-way ANCOVA, Training: F_1.21, 53.15_ = 5.67, *P* = 0.005; Time bin: F_2.62, 115.18_ = 2.16, *P* = 0.10; Training x Time: F_4.48, 196.92_ = 3.56, *P* = 0.006. This overtrain action-initiation inhibited ensemble became more inhibited prior to action initiation as training progressed. **(o-q)** Linear decoding accuracy from DMS A2A^+^ overtrain action-initiation inhibited neuron activity of behavioral switch rate (o; 2-way ANOVA, Neuron activity: F_1, 3_ = 6.21, *P* = 0.09; Training: F_1.78, 5.34_ = 0.20, *P* = 0.80; Neuron activity x Training: F_1.55, 4.64_ = 0.70, *P* = 0.64), the variability of the interpress intervals (p; 2-way ANOVA, Neuron activity: F_1, 3_ = 8.72, *P* = 0.06; Training: F_1.24, 3.71_ = 0.26, *P* = 0.68; Neuron activity x Training: F_1.25, 3.76_ = 0.01, *P* = 0.95), and the prior press interpress interval (q; 2-way ANOVA, Neuron activity: F_1, 3_ = 7.95, *P* = 0.07; Training: F_1.62, 4.87_ = 6.40, *P* = 0.05; Neuron activity x Training: F_1.86, 5.57_ = 2.49, *P* = 0.17). Behavioral switch rate, relative interpress interval, and prior press interpress interval, could not be significantly decoded from the activity of the overtrain action-initiation inhibited A2A^+^ neurons. A2A-cre: *N* = 4 (1 male). Data presented as mean ± s.e.m. Males = solid lines, Females = dashed lines. \**P* < 0.05, \*\**P* < 0.01, \*\*\**P* < 0.001.

**Extended Data Figure 5-4.**
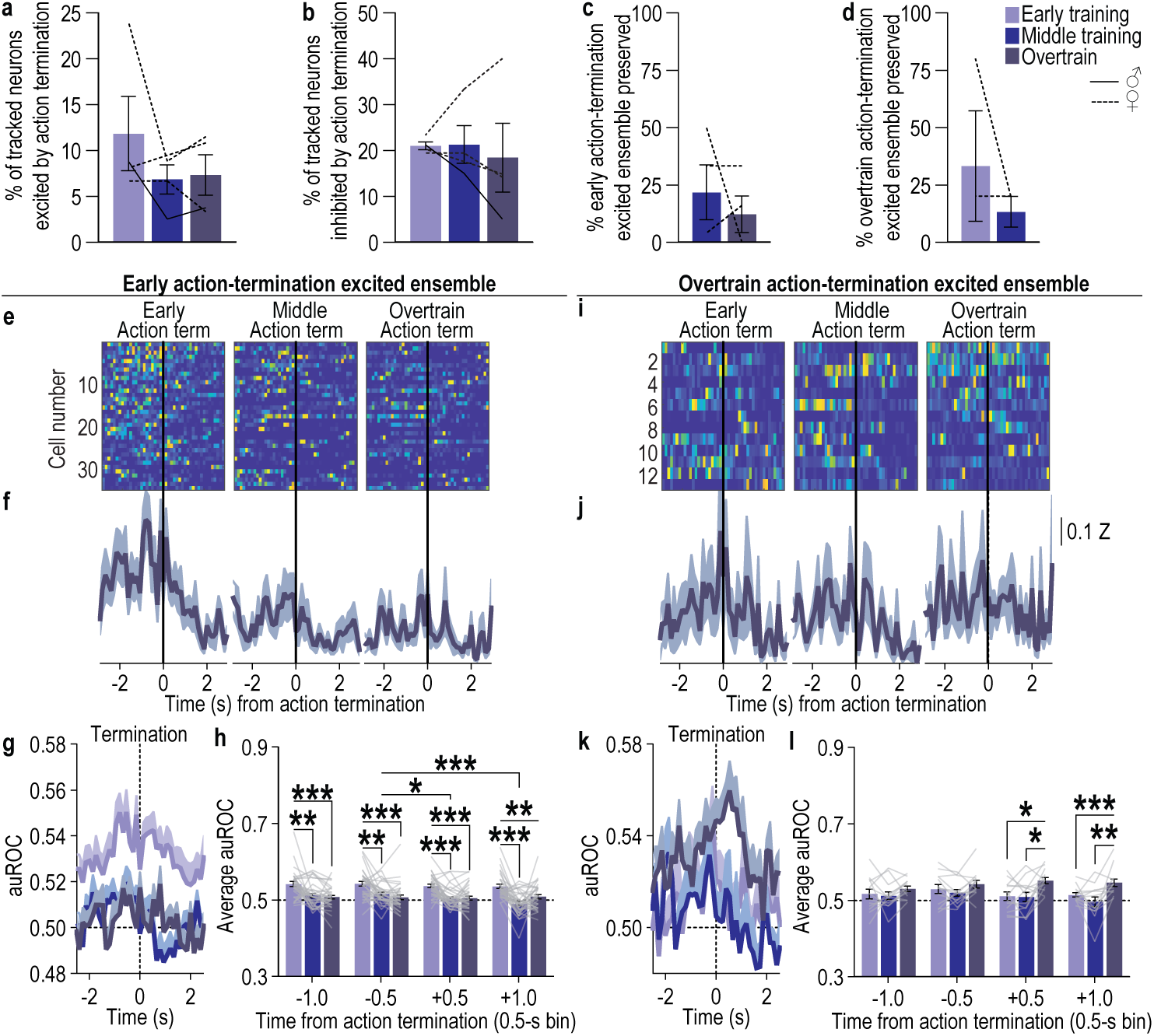
The ensemble of DMS A2A^+^ neurons encoding action termination shifts as habits form. (a-b) Percent of all recorded coregistered DMS A2A^+^ neurons (Average 73.25 coregistered neurons/mouse, s.e.m. 17.09) significantly excited (a; 1-way ANOVA, F_1.48, 4.43_ = 1.12, *P* = 0.38) or inhibited (b; 1-way ANOVA, F_1.02, 3.05_ = 0.20, *P* = 0.69) by action termination. Approximately 4 – 10% of coregistered DMS A2A^+^ neurons were excited by action termination and 19 – 21% were inhibited. This did not change with training. F_1.00, 3.02_ = 1.09, *P* = 0.37. **(c)** Percent of DMS A2A^+^ early action-termination excited neurons that were also significantly excited by action termination on the 4^th^ (middle) and 8^th^ (overtrain) training sessions. Only 12 – 22% of the early action-termination ensemble continued to be excited by action termination during the middle and overtraining phases of training. The proportion preserved did not change with training. Middle v. Overtrain t_3_ = 0.69, *P* = 0.54. **(d)** Percent of DMS A2A^+^ overtrain action-termination excited neurons that were also significantly excited by action termination on the 1^st^ (early) and 4^th^ (middle) training sessions. Only 13 – 33% of the overtraining action-termination ensemble was also excited by action termination during the preceding early and middle training phases. The proportion preserved did not change with training. Early v. Middle t_3_ = 1.00, *P* = 0.42. **(e-h)** Activity and modulation across training of DMS A2A^+^ early action-termination excited neurons. Heat map of minimum to maximum deconvolved activity (sorted by total activity) (e), Z-scored activity (f), and area under the receiver operating characteristic curve (auROC) modulation index (g) of these cells around action termination across training. (h) auROC modulation index averaged across 0.5-s bins around action termination. Training: F_1.66, 54.75_ = 3.77, *P* = 0.03; Time bin: F_1.82, 60.19_ = 1.17, *P* = 0.33; Training x Time: F_4.19, 138.28_ = 1.01, *P* = 0.41. This early action-termination ensemble became less modulated by action termination as training progressed. **(i-l)** Activity and modulation across training of DMS A2A^+^ overtrain action-termination excited neurons. Heat map of minimum to maximum deconvolved activity (i), Z-scored activity (j), and area under the receiver operating characteristic curve (auROC) modulation index (j) of these cells around action termination across training. (k) auROC modulation index averaged across 0.5-s bins around action termination. Training: F_1.82, 19.98_ = 3.20, *P* = 0.07; Time bin: F_1.55, 17.10_ = 2.28, *P* = 0.10; Training x Time: F_3.25, 37.70_ = 0.31, *P* = 0.83. This overtrain action-termination ensemble became slightly more modulated by action termination as training progressed. A2A-cre: *N* = 4 (1 male). Data presented as mean ± s.e.m. Males = solid lines, Females = dashed lines. \**P* < 0.05, \*\**P* < 0.01, \*\*\**P* < 0.001.

**Extended Data Figure 5-5.**
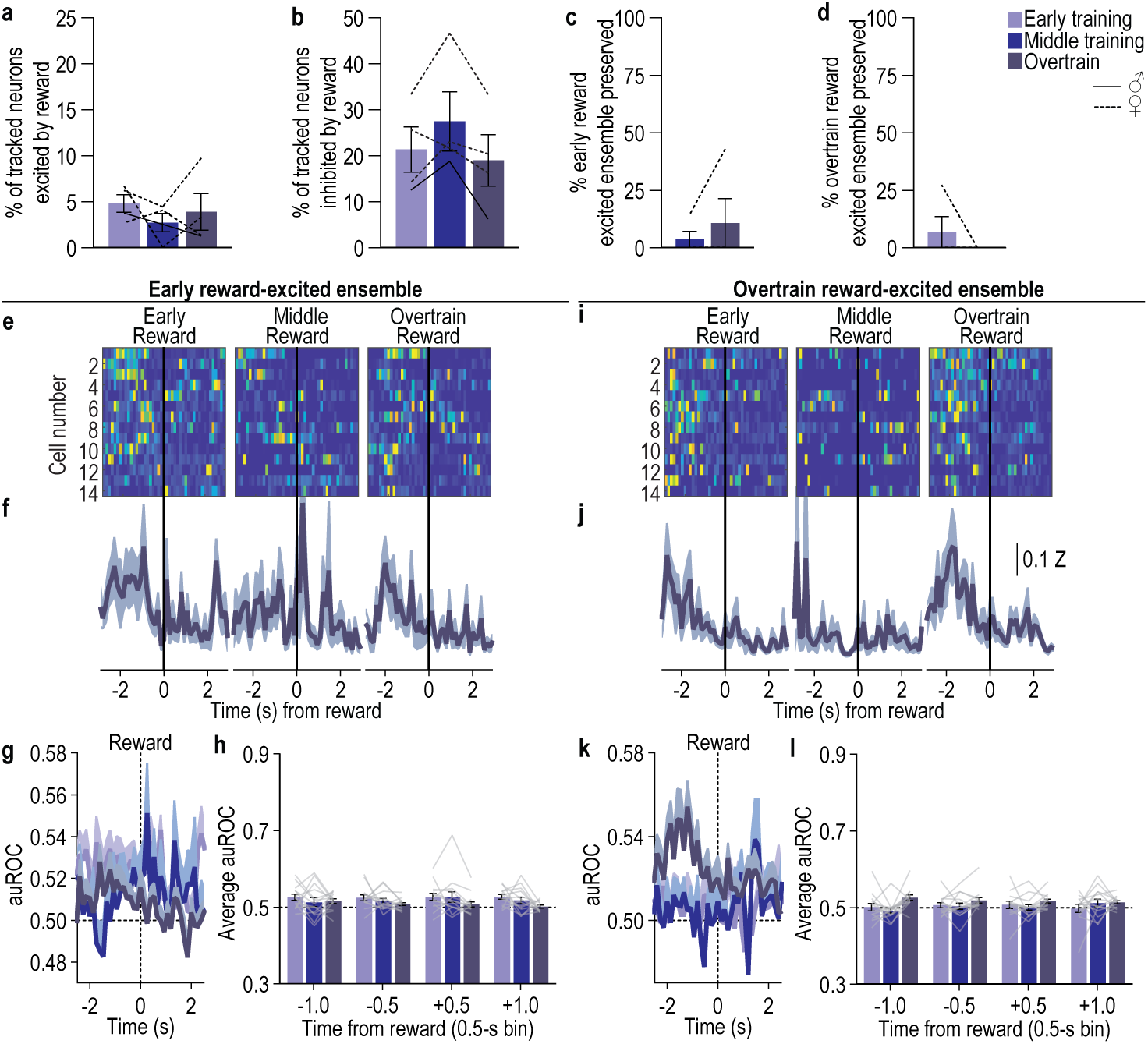
Few DMS A2A^+^ neurons are excited by reward. (a-b) Percent of coregistered DMS A2A^+^ neurons significantly excited (a; 1-way ANOVA, F_1.89, 5.68_ = 0.75, *P* = 0.51) or inhibited (b; 1-way ANOVA, F_1.76, 5.28_ = 3.52, *P* = 0.11) by collection of the earned reward. Only 3 – 5% of coregistered DMS A2A^+^ neurons were excited by reward, substantially more (19 – 27%) were inhibited by reward. **(c)** Percent of DMS A2A^+^ early reward excited neurons that were then significantly excited by reward on the 4^th^ (middle) and 8^th^ (overtrain) training sessions. Only 4 – 11% of the small early reward ensemble continued to be modulated by reward during the middle and overtraining phases of training. The proportion preserved did not change with training. Middle v. Overtrain 2-tailed Wilcoxon signed rank test, W = 1.00, P > 0.99. **(d)** Percent of DMS A2A^+^ overtrain reward excited neurons that were significantly excited by reward on the 1^st^ (early) and 4^th^ (middle) training sessions. Only 0 – 7% of the small overtraining reward ensemble was also modulated by reward during the preceding early and middle training phases. The proportion preserved did not change with training. Early v. Middle 2-tailed Wilcoxon signed rank test, W = –1.00, P > 0.99. **(e-h)** Activity and modulation across training of DMS A2A^+^ early reward excited neurons. Heat map of minimum to maximum deconvolved activity (e), Z-scored activity (f), and area under the receiver operating characteristic curve (auROC) modulation index (g) of these cells around reward across training. (h) auROC modulation index averaged across 0.5-s bins around reward. Training: F_1.53, 18.36_ = 0.492, *P* = 0.57; Time bin: F_1.54, 18.49_ = 0.64, *P* = 0.50; Training x Time: F_2.58, 30.98_ = 0.41, *P* = 0.71. **(i-l)** Activity and modulation across training of DMS A2A^+^ overtrain reward excited neurons. Heat map of minimum to maximum deconvolved activity (i), Z-scored activity (j), and area under the receiver operating characteristic curve (auROC) modulation index (k) of these cells around reward on across training. (l) auROC modulation index averaged across 0.5-s bins around reward. Training: F_1.51, 18.10_ = 1.07, *P* = 0.36; Time bin: F_1.98, 23.76_ = 2.78, *P* = 0.08; Training x Time: F_2.94, 35.24_ = 0.65, *P* = 0.58. A2A-cre: *N* = 4 (1 male). Data presented as mean ± s.e.m. Males = solid lines, Females = dashed lines.

**Extended Data Figure 5-6.**
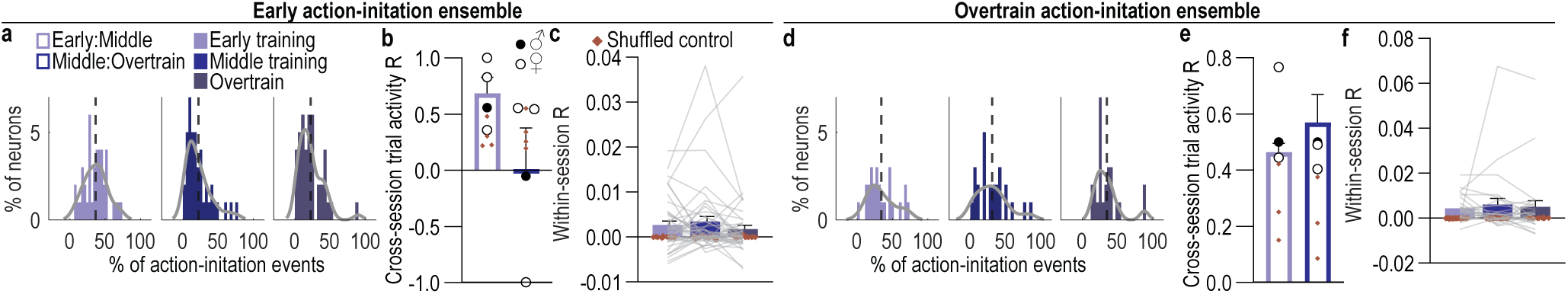
Fidelity with which DMS A2A^+^ neurons encode action initiation. (a-c) Fidelity with which A2A^+^ early action-initiation excited neurons encode action initiation. **(a)** Distribution of the percentage of early action-initiation excited neurons as a function of the percentage of action-initiation events to which they respond for each training phase. **(b)** Cross-session correlation of the response distributions. 2-way ANOVA, Neuron activity distribution (v. shuffled): F_1, 3_ = 0.03, *P* = 0.88; Training sessions: F_1, 3_ = 1.34, *P* = 0.33; Distribution x Training: F_1, 3_ = 2.99, *P* = 0.18. Early action-initiation A2A^+^ neurons tend to respond on less than half of the action-initiation events and this decreases with training and is not correlated across training sessions. **(c)** Within-session correlation of the activity around action initiation of each early action-initiation excited neuron. 2-way ANCOVA, Neuron activity: F_1, 36_ = 2.71, *P* = 0.11; Training: F_2, 72_= 2.47, *P* = 0.09; Activity x Time: F_2, 72_ = 2.45, *P* = 0.09. The activity of early action-initiation excited A2A^+^ neurons around action initiation is not significantly correlated within a training session. **(d-e)** Fidelity with which A2A^+^ overtrain action-initiation excited neurons encode action initiation. **(d)** Distribution of the percentage of overtrain action-initiation excited neurons as a function of the percentage of action-initiation events to which they respond for each training phase. **(e)** Cross-session correlation of the response distributions. 2-way ANOVA, Neuron activity distribution: F_1, 2_ = 32.64, *P* = 0.03; Training sessions: F_1, 2_ = 0.54, *P* = 0.54; Distribution x Training: F_1, 2_ = 0.45, *P* = 0.57. Overtrain action-initiation A2A^+^ neurons tend to respond on less than half the action-initiation events across training and this is consistent across training. **(f)** Within-session correlation of the activity around action initiation of each overtrain action-initiation excited neuron. 2-way ANCOVA, Neuron activity: F_1, 22.00_= 0.40, *P* = 0.53; Training: F_1.24, 27.30_ = 0.63, *P* = 0.47; Activity x Time: F_1.23, 27.09_ = 0.65, *P* = 0.46. The activity of overtrain action-initiation excited A2A^+^ neurons around action initiation is not correlated above shuffled control within a training session. A2A-cre: *N* = 4 (1 male). Data presented as mean ± s.e.m. Males = closed circles, Females = open circles.

**Extended Data Figure 5-7.**
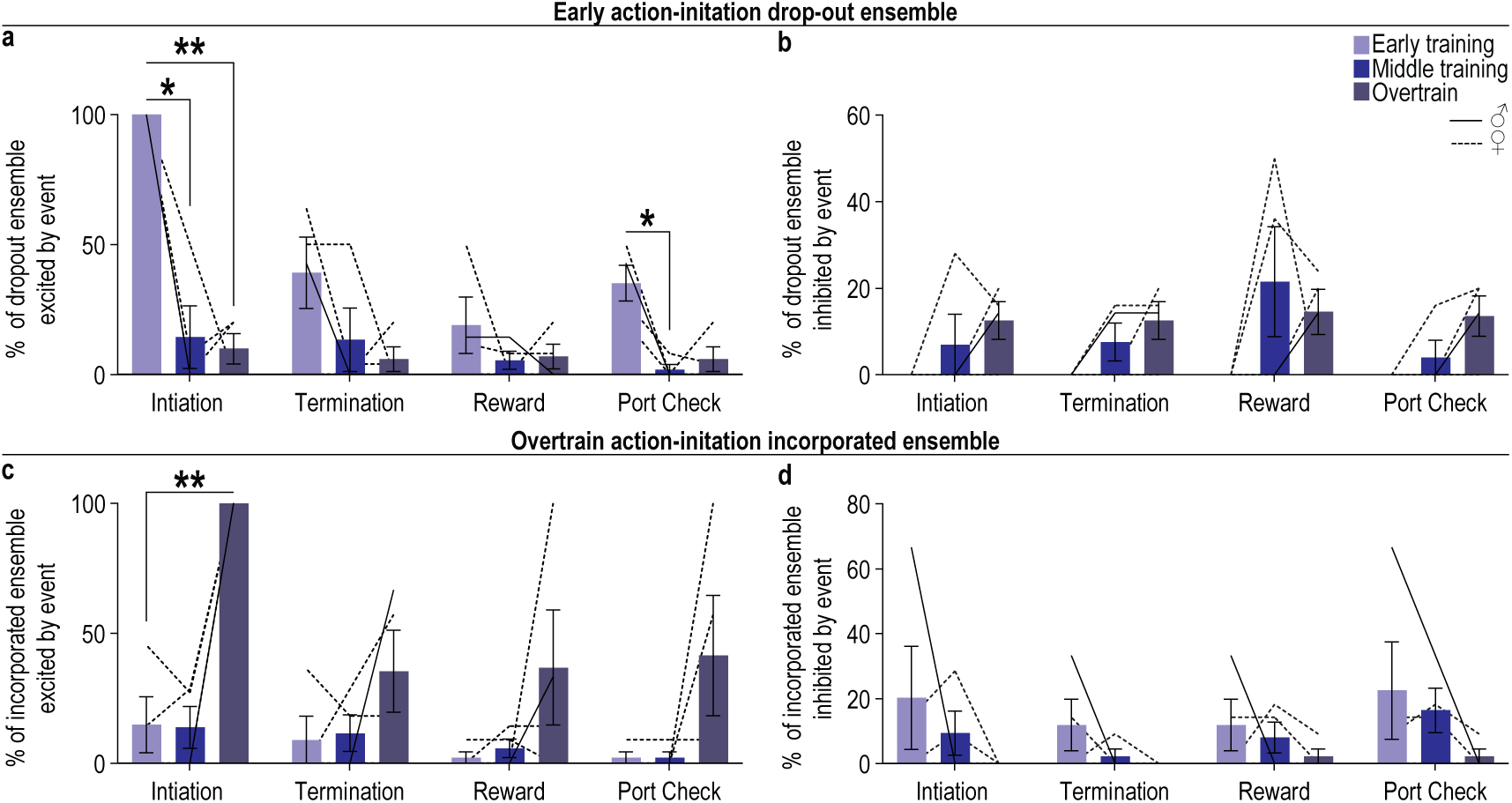
Encoding of other task variables by A2A^+^ neurons that drop out of the early action-initiation ensemble or get incorporated into the overtrain action-initiation excited ensemble. (a-b) Percentage of the neurons that drop out of the early action-initiation excited ensemble that are excited (a; 2-way ANOVA, Event x Training: F_1.79, 5.37_ = 7.05, *P* = 0.03; Event: F_1.21, 3.63_ = 30.74, *P* = 0.006; Training: F_1.13, 3.40_ = 14.46, *P* = 0.02) or inhibited (b; 2-way ANOVA, Event: F_1.98, 5.94_ = 1.64, *P* = 0.27; Training: F_1.50, 4.50_ = 2.77, *P* = 0.17; Event x Training: F_1.46, 4.37_ = 1.25, *P* = 0.35) by other events (terminating lever press, reward, food-delivery port check). By definition, these neurons stop being activated by action initiation with training. They do not begin to be activated by other task events with training. Instead, fewer of these neurons are activated by other task events and some of them become inhibited by tasks events. **(c-d)** Percentage of the neurons that are incorporated into the overtrain action-initiation excited ensemble that are excited (c; 2-way ANOVA, Training: F_1.1, 3.48_ = 11.06, *P* = 0.03; Event: F_2.24, 6.71_ = 3.67, *P* = 0.08; Event x Training: F_1.54, 4.62_ = 2.41, *P* = 0.19) or inhibited (d; 2-way ANOVA, Event: F_1.16, 3.48_ = 1.95, *P* = 0.25; Training: F_1.09, 3.28_ = 1.24, *P* = 0.35; Event x Training: F_1.89, 5.68_ = 0.83, *P* = 0.48) by other events (terminating lever press, reward, food-delivery port check). By definition, these neurons became activated by action initiation at overtraining. Earlier in training very few of these neurons were activated by other events. A small proportion of these neurons were inhibited by task events earlier in training and this proportion decreased to near 0 with training. Data presented as mean ± s.e.m. Males = solid lines, Females = dashed lines. \**P* < 0.05, \*\**P* < 0.01.

**Extended Data Figure 5-8:**
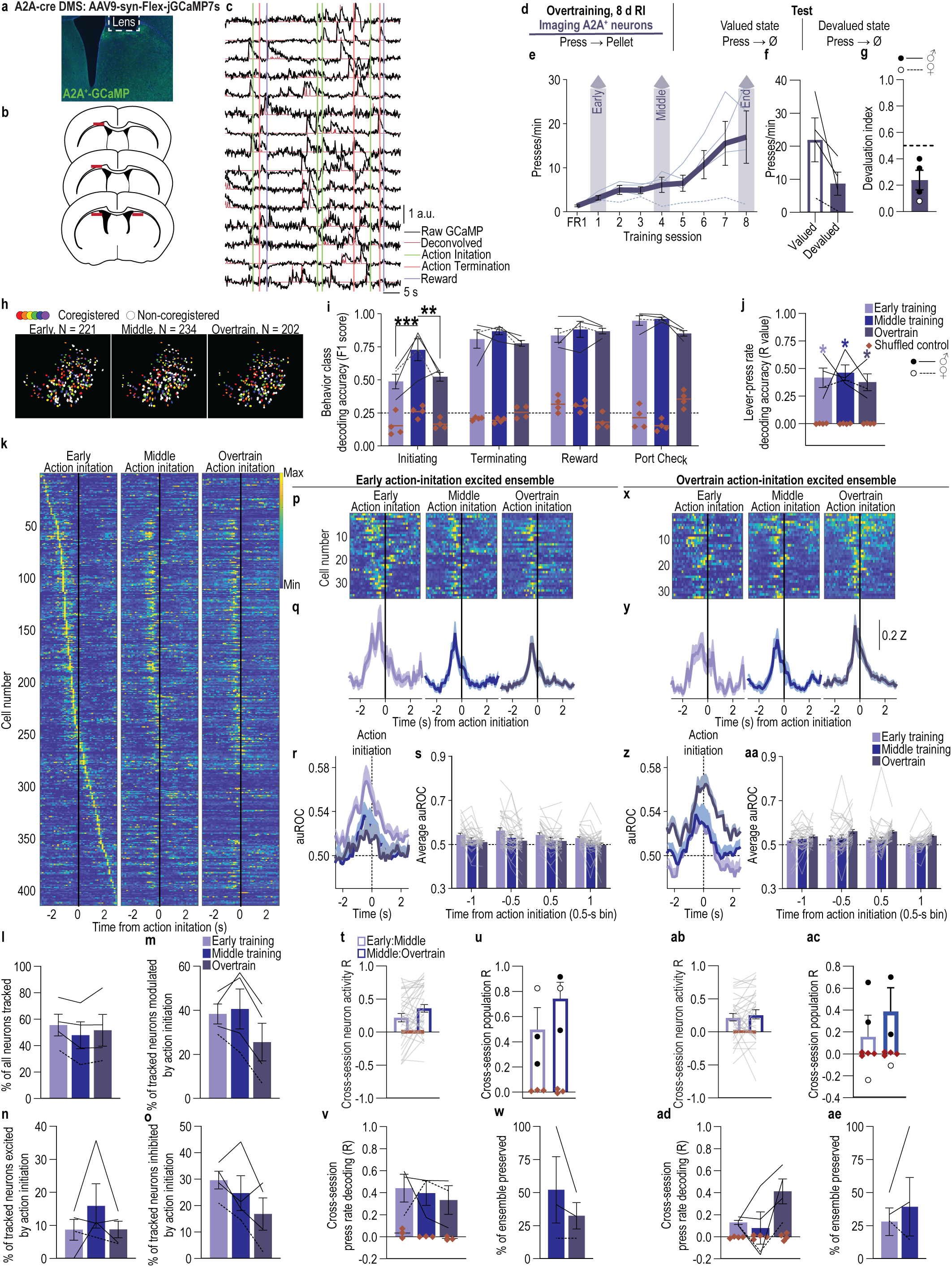
The ensemble of DMS A2A^+^ neurons that encodes action initiation is more stable if subjects remain goal-directed with overtraining. **(a)** Representative image of cre-dependent jGCaMP7s expression in DMS A2A^+^ neurons. **(b)** Map of DMS GRIN lens placements for all subjects. **(c)** Representative raw fluorescence (normalized to max) and deconvolved calcium signal. **(d)** Procedure. RI, random-interval reinforcement schedule. **(e)** Training press rate. 2-way ANOVA, Training: F_1.49, 4.48_ = 5.58, *P* = 0.06. **(f)** Test press rate. 2-tailed t-test, t_3_ = 2.30, *P* = 0.11, 95% CI –31.60 – 5.10. **(g)** Devaluation index [(Devalued condition presses)/(Valued condition presses + Devalued presses)]. One-tailed Bayes factor, BF_10_ = 5.40. **(h)** Representative recorded A2A^+^ neuron spatial footprints during the first (early), 4^th^ (middle), and 8^th^ (overtrain) random-interval training session. Colored, coregistered neurons; white, non-coregistered neurons. **(i)** Behavior class (initiating lever press, terminating press, reward collection, non-reinforced food-port check) decoding accuracy from A2A^+^ coregistered neuron activity compared to shuffled control. Line = chance. 3-way ANOVA, Neuron activity (v. shuffled): F_1, 3_ = 548.47, *P* < 0.001; Training session: F_2, 6_ = 4.11, *P* = 0.08; Behavior class: F_3, 9_ = 40.4, *P* < 0.001; Neuron activity x Training: F_2, 6_ = 5.95, *P* = 0.038; Neuron activity x Behavior: F_3, 9_ = 16.24, *P* < 0.001; Training x Behavior: F_6, 18_ = 2.55, *P* = 0.02; Neuron activity x Training x Behavior: F_6, 18_ = 5.11, *P* = 0.003. **(j)** Lever-press rate decoding accuracy from A2A^+^ coregistered neuron activity. R = correlation between observed and decoded values. 2-way ANOVA, Neuron activity: F_1, 3_ = 195.5, *P* = 0.0008; Training: F_1.64, 4.92_ = 0.23, *P* = 0.77; Neuron activity x Training: F_1.62, 4.86_ = 0.23, *P* = 0.76. **(k)** Heat map of minimum to maximum deconvolved activity (sorted by total activity) of each coregistered DMS A2A^+^ neuron around lever-press action initiation. **(l)** Percent of all A2A^+^ neurons coregistered across training in subjects that did not form habits with overtraining. 1-way ANOVA, F_1.32, 3.95_ = 2.82, *P* = 0.17. **(m-o)** Percent of coregistered neurons (Average 106.5 coregistered neurons/mouse, s.e.m. 40.34) significantly modulated (m; 1-way ANOVA, F_1.05, 3.16_ = 8.14, *P* = 0.06), excited (n; 1-way ANOVA, F_1.14, 3.43_ = 1.72, *P* = 0.28), or inhibited (o; 1-way ANOVA, F_1.90, 5.70_ = 3.37, *P* = 0.11) around action initiation. **(p-s)** Activity and modulation across training of DMS A2A^+^ early action-initiation excited neurons. Heat map (p), Z-scored activity (q), and area under the receiver operating characteristic curve (auROC) modulation index (r) of early action-initiation excited neurons around action initiation across training. (s) Modulation index averaged across 0.5-s bins around action initiation. 2-way ANCOVA, Training: F_1.82, 64.00_ = 0.48, *P* = 0.62; Time bin: F_2.18, 76.27_ = 0.07, *P* = 0.94; Training x Time: F_3.78, 132.35_ = 1.61, *P* = 0.18. One subject without any neurons significantly excited around action initiation during early training was removed from these and subsequent related analysis. **(t-u)** Cross-session correlation of the activity around action initiation of each early action-initiation excited neuron (t; 2-way ANCOVA, Neuron activity: F_1, 35_ = 0.02, *P* = 0.90; Training: F_1, 35_ = 0.001, *P* = 0.98; Activity x Time: F_1, 35_ = 0.01, *P* = 0.91) or the population activity of these neurons (u; 2-way ANOVA, Neuron activity: F_1, 2_ = 31.38, *P* = 0.03; Training: F_1, 2_ = 1.25, *P* = 0.38; Activity x Time: F_1, 2_ = 1.21, *P* = 0.39). **(v)** Cross-session decoding accuracy of lever-press rate from the activity of A2A^+^ early action-initiation-excited neuron population activity on the 1^st^ training session. 2-way ANOVA, Neuron activity: F_1, 2_ = 23.74, *P* = 0.04; Training: F_1.03, 2.05_ = 0.36, *P* = 0.61; Neuron activity x Training: F_1.04, 2.07_ = 0.07, *P* = 0.83. When we trained a linear regression model to decode action rate from the activity of early action-initiation excited A2A^+^ neurons on the first training session, we could reliably decode action rate not only from this session, but also similarly from the following training sessions. **(w)** Percent of A2A^+^ early action-initiation excited neurons that continued to be significantly excited by action initiation on the 4^th^ and 8^th^ training sessions. 2-tailed t-test, t_2_ = 1.28, *P* = 0.33, 95% CI –85.86 – 46.47. **(x-aa)** Activity and modulation across training of A2A^+^ overtrain action-initiation excited neurons. Heat map (x), Z-scored activity (y), and auROC modulation index (z) of overtrain action-initiation excited neurons around action initiation across training. (aa) Modulation index around action initiation. 2-way ANCOVA, Training: F_1.65, 51.23_ = 0.43, *P* = 0.43; Time bin: F_2.16, 67.03_ = 0.17, *P* = 0.86; Training x Time: F_4.31, 133.64_ = 0.59, *P* = 0.68. **(ab-ac)** Cross-session correlation of the activity around action initiation of each overtrain action-initiation excited neuron (ab; 2-way ANCOVA, Neuron activity: F_1, 31_ = 1.77, *P* = 0.19; Training: F_1, 31_ = 0.28, *P* = 0.60; Activity x Time: F_1, 31_ = 0.20, *P* = 0.66) or the population activity of these neurons (ac; 2-way ANOVA, Neuron activity: F_1, 3_ = 2.37, *P* = 0.22; Training: F_1, 3_ = 1.20, *P* = 0.35; Activity x Time: F_1, 3_ = 1.13, *P* = 0.37). **(ad)** Cross-session decoding accuracy of lever-press rate from the activity of A2A^+^ overtrain action-initiation-excited neuron population activity on the 8^th^ training session. 2-way ANOVA, Neuron activity: F_1, 3_ = 6.91, *P* = 0.08; Training: F_1.87, 5.60_ = 5.28, *P* = 0.05; Neuron activity x Training: F_1.90, 5.69_ = 3.47, *P* = 0.10. **(ae)** Percent of A2A^+^ overtrain action-initiation excited neurons that were also significantly excited by action initiation on 1^st^ and 4^th^ training sessions. 2-tailed t-test, t_3_ = 0.82, *P* = 0.47, 95% CI –32.59 – 55.21. A2A-cre: *N* = 4 (3 male). Data presented as mean ± s.e.m. Males = closed circles/solid lines, Females = open circles/dashed lines. \**P* < 0.05, \*\**P* < 0.01, \*\*\**P* < 0.001.

**Extended Data Figure 6-1.**
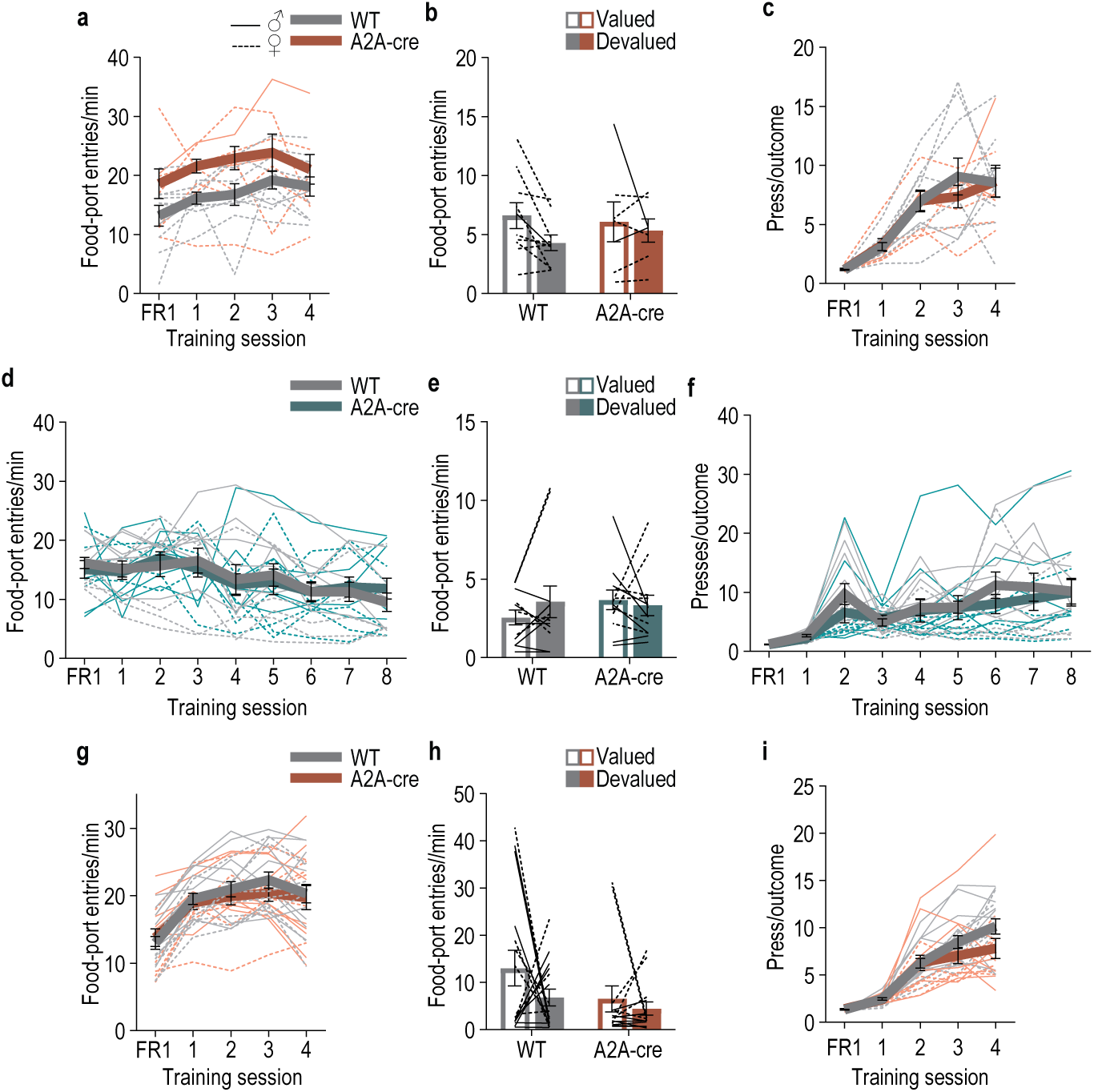
Food-port entries during training and test with DMS A2A^+^ neuron chemogenetic manipulation. (a-c) Chemogenetic inactivation of DMS A2A^+^ neurons during learning. WT: *N* = 10 (1 male); A2A-cre: *N* = 7 (2 males). **(a)** Training entry rate. 2-way ANOVA, Training: F_2.87, 43.12_ = 3.50, *P* = 0.02; Genotype: F_1, 15_ = 6.63, *P* = 0.02; Training x Genotype: F_4, 60_ = 0.31, *P* = 0.87. **(b)** Test entry rate. 2-way ANOVA, Value x Genotype: F_1, 15_ = 1.08, *P* = 0.32; Value: F_1, 15_ = 4.06, *P* = 0.06; Genotype: F_1, 15_ = 0.03, *P* = 0.86. **(c)** Training average lever presses/earned reward outcome. 2-way ANOVA, Training: F_2.38, 35.73_ = 32.06, *P* < 0.0001; Genotype: F_1, 15_ = 0.08, *P* = 0.78; Training x Genotype: F_4, 60_ = 0.47, *P* = 0.76. **(d-f)** Chemogenetic activation of DMS A2A^+^ neurons during learning. WT: *N* = 12 (7 males); A2A-cre: *N* = 12 (6 males). **(d)** Training entry rate. 2-way ANOVA, Training: F_4.22, 92.89_ = 6.10, *P* = 0.0002; Genotype: F_1, 22_ = 0.00008, *P* = 0.98; Training x Genotype: F_8, 176_ = 0.54, *P* = 0.83. **(e)** Test entry rate. 2-way ANOVA, Value x Genotype: F_1, 22_ = 1.50, *P* = 0.23; Value: F_1, 22_ = 0.35, *P* = 0.56; Genotype: F_1, 22_ = 0.27, *P* = 0.61. **(f)** Training average lever presses/earned reward outcome. 2-way ANOVA, Training: F_2.42, 53.34_ = 16.33, *P* < 0.0001; Genotype: F_1, 22_ = 0.35, *P* = 0.56; Training x Genotype: F_8, 176_ = 0.76, *P* = 0.64. **(g-i)** Chemogenetic inactivation of DMS A2A^+^ neurons during test after learning. WT: *N* = 16 (9 males); A2A-cre: *N* = 14 (8 males). **(g)** Training entry rate. 2-way ANOVA, Training: F_1.93, 53.92_ = 28.28, *P* < 0.0001; Genotype: F_1, 28_ = 0.23, *P* = 0.63; Training x Genotype: F_4, 112_ = 0.70, *P* = 0.59. **(h)** Test entry rate. 2-way ANOVA, Value x Genotype: F_1, 28_ = 0.47, *P* = 0.50; Value: F_1, 28_ = 1.82, *P* = 0. 19; Genotype: F_1, 28_ = 4.08, *P* = 0.05. **(i)** Training average lever presses/earned reward outcome. 2-way ANOVA, Training: F_2.19, 61.42_ = 81.93, *P* < 0.0001; Genotype: F_1, 28_ = 1.55, *P* = 0.22; Training x Genotype: F_4, 112_ = 2.25, *P* = 0.07. Data presented as mean ± s.e.m. Males = solid lines, Females = dashed lines. Chemogenetic inhibition of DMS A2A^+^ neurons increased food-port entries during training, whereas activation did not significantly affect food-port entries. Neither manipulation significantly altered the number of lever presses per earned reward.

**Extended Data Figure 6-2.**
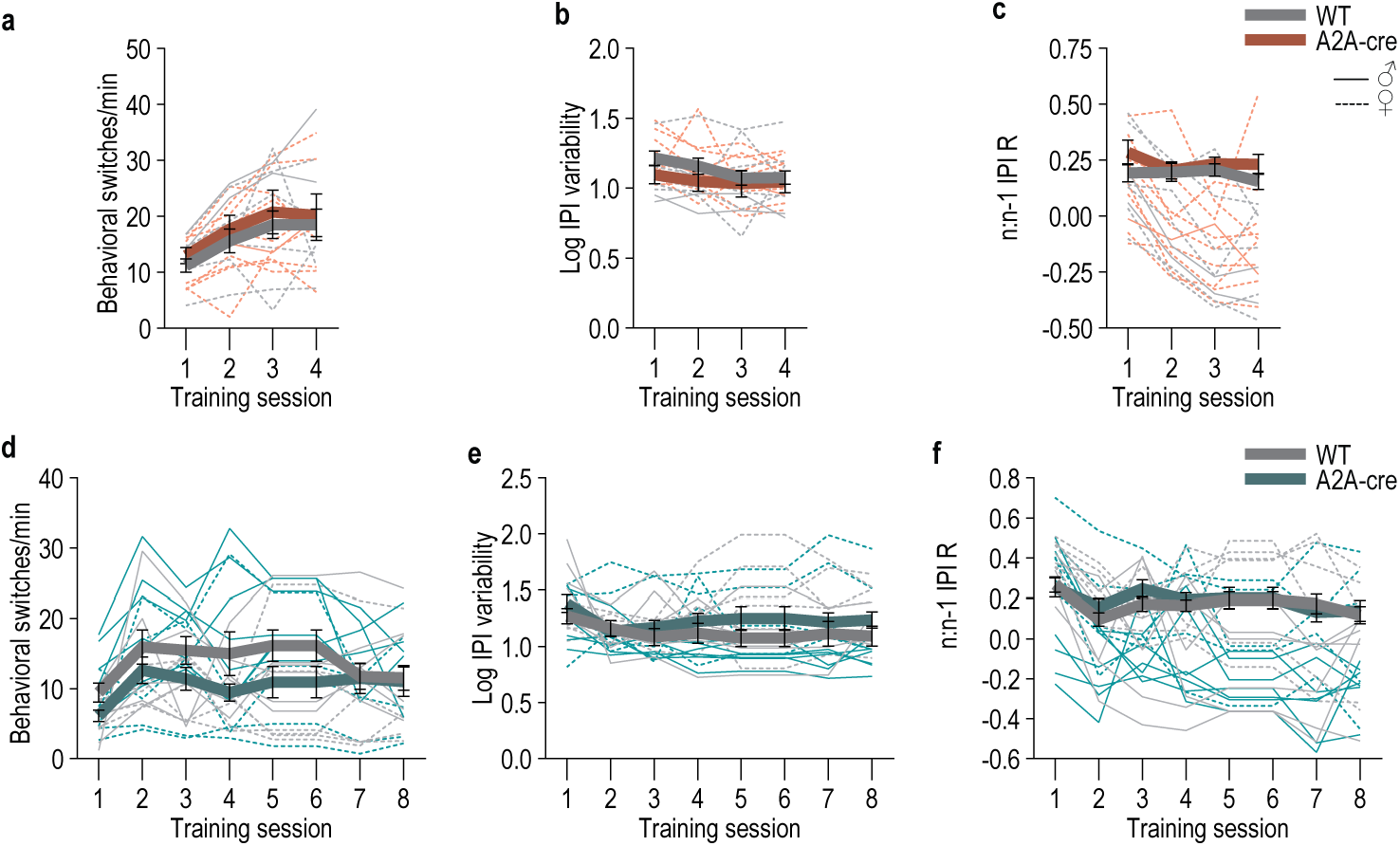
Behavioral engagement and organization variables during training with DMS A2A^+^ neuron chemogenetic manipulation. (a-c) Chemogenetic inactivation of DMS A2A^+^ neurons during learning. WT: *N* = 10 (1 male); A2A-cre: *N* = 7 (2 males). Behavioral switch rate (a; how often subjects stop one behavior to transition to another i.e., press ◊ food-port check or food-port check ◊ press; 2-way ANOVA, Training: F_2.30, 36.79_ = 16.17, *P* < 0.0001; Genotype: F_1, 16_ = 0.02, *P* = 0.89; Training x Genotype: F_2.30, 36.79_ = 0.11, *P* = 0.92), interpress interval variability (b; standard deviation of the log-transformed interpress-intervals; 2-way ANOVA, Training: F_2.64, 42.21_ = 2.47, *P* = 0.08; Genotype: F_1, 16_ = 0.96, *P* = 0.34; Training x Genotype F_2.64, 42.21_ = 0.59, *P* = 0.60), and relationship to past trial behavior (c; correlation between the current interpress interval (n) and the interpress interval of the prior press (n-1); 2-way ANOVA, Genotype: F_1, 16_ = 1.27, *P* = 0.28; Training: F_2.54, 40.70_ = 1.15, *P* = 0.33; Training x Genotype: F_2.54, 40.70_ = 1.06, *P* = 0.37) across days of training. **(d-f)** Chemogenetic activation of DMS A2A^+^ neurons during learning. WT: *N* = 12 (7 males); A2A-cre: *N* = 12 (6 males). Behavioral switch rate (d; 2-way ANOVA, Training: F_3.38, 74.41_ = 4.51, *P* = 0.004; Genotype: F_1, 22_ = 2.39, *P* = 0.13; Training x Genotype: F_3.38, 74.41_ = 1.09, *P* = 0.36), interpress interval variability (e; Training: F_2.97, 65.26_ = 2.40, *P* = 0.08; Genotype: F_1, 22_ = 1.43, *P* = 0.24; Training x Genotype: F_2.97, 65.26_ = 0.66, *P* = 0.58), and relationship to past trial behavior (f; 2-way ANOVA, Genotype: F_1, 22_ = 0.21, *P* = 0.65; Training: F_3.14, 69.01_ = 3.34, *P* = 0.02; Training x Genotype: F_3.14, 69.01_ = 0.55, *P* = 0.65) across days of training. Neither inhibition nor excitation of DMS A2A^+^ neurons affected behavioral engagement and organization variables. Data presented as mean ± s.e.m. Males = solid lines, Females = dashed lines.

**Extended Data Figure 6-3.**
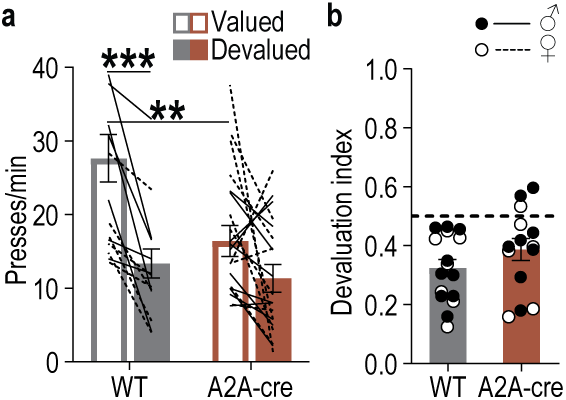
Press rate during devaluation test with DMS A2A^+^ neuron inhibition. **(a)** Test press rate. 2-way ANOVA, Value x Genotype: F_1, 28_ = 6.46, *P* = 0.02; Value: F_1, 28_ = 28.86, *P* < 0.001; Genotype: F_1, 28_ = 5.15, *P* = 0.03. **(b)** Devaluation index. 2-tailed t-test, t_28_ = 1.36; *P* = 0.18, 95% CI –0.03 – 0.16. Data presented as mean ± s.e.m. Males = closed circles/solid lines, Females = open circles/dashed lines. \*\**P* < 0.01, \*\*\**P* < 0.001.

## SUPPLEMENTAL TABLES

**Supplemental Table 1:**
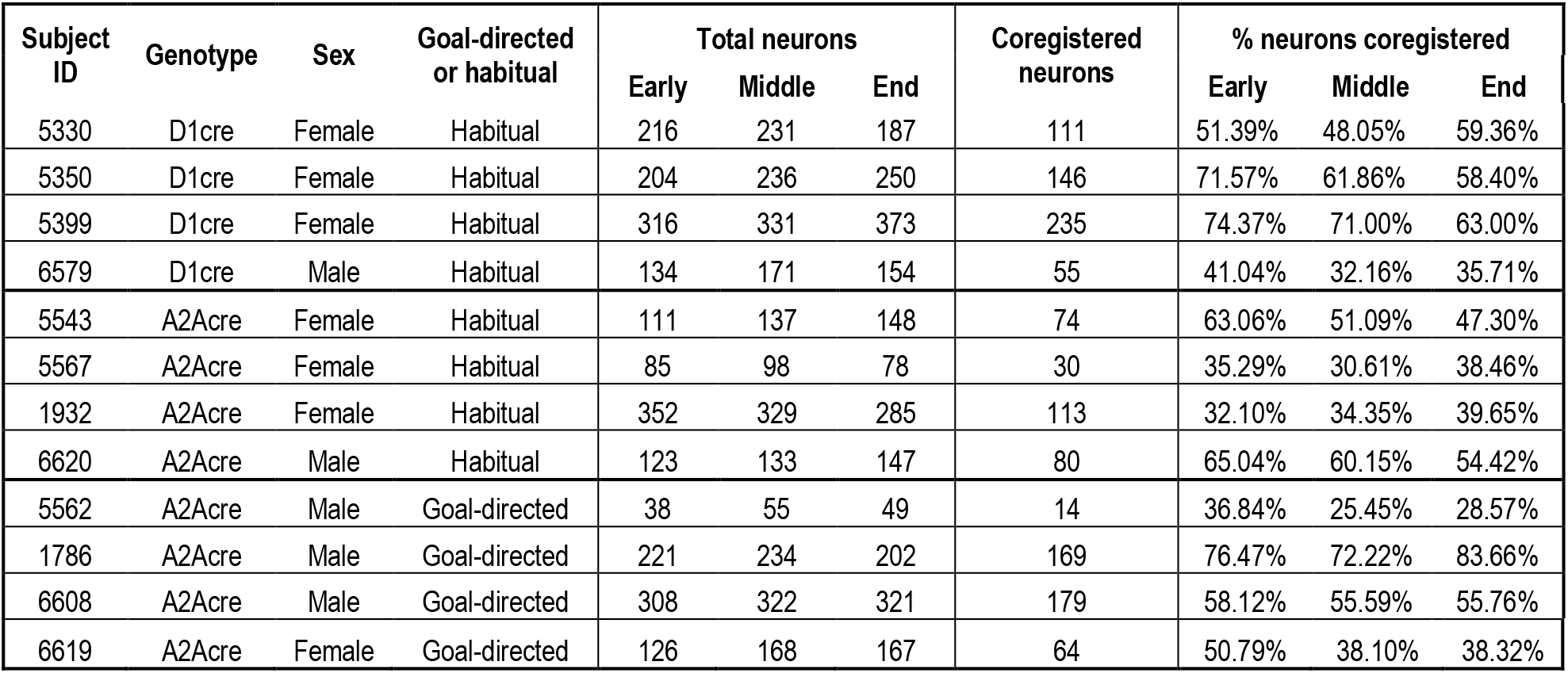
Neurons recorded and tracked for each subject.

**Supplemental Table 2:** Full statistical reporting for main text figures.

| Figure | Statistical test | Statistics |
|---|---|---|
| 1f | 1-way ANOVA | Training: F<sub>1.81, 5.43</sub> = 4.84, *P* = 0.06 |
| 1g | 2-tailed t-test | t<sub>3</sub> = 1.98, *P* = 0.14, 95% confidence interval (CI) -0.85 - 3.65 |
| 1h | One-tailed Bayes factor | BF<sub>10</sub> = 0.18 |
| 1i | 3-way ANOVA | Neuron activity (v. shuffled): F<sub>1, 3</sub> = 1092.83, *P* < 0.001; Training session: F<sub>1.17, 3.51</sub> = 2.40, *P* = 0.21; Behavior class: F<sub>1.76, 5.28</sub> = 15.56, *P* = 0.007; Neuron activity x Training: F<sub>1.24, 3.72</sub> = 4.60, *P* = 0.10; Neuron activity x Behavior: F<sub>1.81, 5.43</sub> = 6.49, *P* = 0.04; Training x Behavior: F<sub>2.15, 6.46</sub> = 3.25, *P* = 0.10; Neuron activity x Training x Behavior: F<sub>1.95, 5.86</sub> = 0.69, *P* = 0.54 |
| 1j | 2-way ANOVA | Neuron activity: F<sub>1, 3</sub> = 12.16, *P* = 0.04; Training: F<sub>1.42, 4.25</sub> = 0.07, *P* = 0.87; Neuron activity x Training: F<sub>1.22, 3.68</sub> = 0.08, *P* = 0.84 |
| 1k | 2-way ANOVA | Neuron activity: F<sub>1, 3</sub> = 33.57, *P* = 0.01; Training: F<sub>1.30, 3.89</sub> = 4.28, *P* = 0.11; Neuron activity x Training: F<sub>1.15, 3.44</sub> = 0.27, *P* = 0.66 |
| 1l | 2-way ANOVA | Neuron activity: F<sub>1, 3</sub> = 8.91, *P* = 0.05; Training: F<sub>1.37, 4.11</sub> = 0.76, *P* = 0.48; Neuron activity x Training: F<sub>1.23, 3.74</sub> = 0.18, *P* = 0.75 |
| 1m | 2-way ANOVA | Neuron activity: F<sub>1, 3</sub> = 22.10, *P* = 0.02; Training: F<sub>1.12, 3.36</sub> = 1.62, *P* = 0.29; Neuron activity x Training: F<sub>1.06, 3.19</sub> = 0.26, *P* = 0.67 |
| 1p | 1-way ANOVA | Training: F<sub>1.60, 4.79</sub> = 8.46, *P* = 0.03 |
| 1q | 1-way ANOVA | Training: F<sub>1.02, 3.07</sub> = 29.20, *P* = 0.01 |
| 2b | 1-way ANOVA | F<sub>1.13, 3.63</sub> = 0.14, *P* = 0.76 |
| 2c | 1-way ANOVA | F<sub>1.29, 3.86</sub> = 0.14, *P* = 0.79 |
| 2g | 2-way ANCOVA | Neuron activity: F<sub>1, 75</sub> = 22.92, *P* < 0.001; Training: F<sub>1, 75</sub> = 0.40, *P* = 0.53; Activity x Training: F<sub>1, 75</sub> = 0.24, *P* = 0.63 |
| 2h | 2-way ANOVA | Neuron activity: F<sub>1, 3</sub> = 39.10, *P* = 0.008; Training: F<sub>1, 3</sub> = 0.70, *P* = 0.18; Activity x Training: F<sub>1, 3</sub> = 0.16, *P* = 0.72 |
| 2i | 3-way ANOVA | Neuron activity (v. shuffled): F<sub>1, 3</sub> = 1118.27, *P* < 0.001; Training: F<sub>2, 6</sub> = 2.79, *P* = 0.14; Behavior class: F<sub>3, 9</sub> = 16.94, *P* < 0.001; Neuron activity x Training: F<sub>2, 6</sub> = 2.10, *P* = 0.20; Neuron activity x Behavior: F<sub>3, 9</sub> = 3.53, *P* = 0.06; Training x Behavior: F<sub>6, 18</sub> = 3.29, *P* = 0.23; Neuron activity x Training x Behavior: F<sub>6, 18</sub> = 0.30, *P* = 0.93 |
| 2j | 2-way ANOVA | Neuron activity: F<sub>1, 3</sub> = 21.00, *P* = 0.02; Training: F<sub>1.02, 3.05</sub> = 0.75, *P* = 0.45; Neuron activity x Training: F<sub>1.01, 3.03</sub> = 0.26, *P* = 0.65 |
| 2k | 2-way ANOVA | Neuron activity: F<sub>1, 3</sub> = 30.27, *P* = 0.01; Training: F<sub>1.29, 4.47</sub> = 4.47, *P* = 0.10; Neuron activity x Training: F<sub>1.18, 3.55</sub> = 2.34, *P* = 0.21 |
| 2l | 2-way ANOVA | Neuron activity: F<sub>1, 3</sub> = 24.74, *P* = 0.01; Training: F<sub>1.38, 4.13</sub> = 1.56, *P* = 0.30; Neuron activity x Training: F<sub>1.63, 4.84</sub> = 2.99, *P* = 0.14 |
| 2m | 2-way ANOVA | Neuron activity: F<sub>1, 3</sub> = 21.31, *P* = 0.02; Training: F<sub>1.38, 4.13</sub> = 1.56, *P* = 0.30; Neuron activity x Training: F<sub>1.25, 3.75</sub> = 0.84, *P* = 0.44 |
| 2n | Planned, Bonferroni corrected, 2-tailed t-tests | Early: t<sub>6</sub> = 7.77, *P* = 0.0007, 95% CI 0.24 - 0.59; Middle: t<sub>6</sub> = 4.47, *P* = 0.01, 95% CI 0.06 - 0.41; Overtrain: t<sub>6</sub> = 4.37, *P* = 0.01, 95% CI 0.06 - 0.41 |
| 2r | 2-way ANCOVA | Neuron activity: F<sub>1, 74</sub> = 21.06, *P* < 0.001; Training: F<sub>1, 74</sub> = 0.23, *P* = 0.64; Activity x Time: F<sub>1, 74</sub> = 0.26, *P* = 0.61 |
| 2s | 2-way ANOVA | Neuron activity: F<sub>1, 3</sub> = 8.25, *P* = 0.06; Training: F<sub>1, 3</sub> = 1.76, *P* = 0.28; Activity x Time: F<sub>1, 3</sub> = 2.29, *P* = 0.23 |
| 2t | 3-way ANOVA | Neuron activity: F<sub>1, 3</sub> = 262.75, *P* < 0.001; Training: F<sub>2, 6</sub> = 2.78, *P* = 0.14; Behavior F<sub>3, 9</sub> = 11.31, *P* = 0.002; Neuron activity x Training: F<sub>2, 6</sub> = 2.16, *P* = 0.20; Neuron activity x Behavior: F<sub>3, 9</sub> = 3.86, *P* = 0.05; Training x Behavior: F<sub>6, 18</sub> = 3.27, *P* = 0.025; Neuron activity x Training x Behavior: F<sub>6, 18</sub> = 0.52, *P* = 0.79 |
| 2u | 2-way ANOVA | Neuron activity: F<sub>1, 3</sub> = 21.00, *P* = 0.02; Training: F<sub>1.02, 3.05</sub> = 0.75, *P* = 0.45; Neuron activity x Training: F<sub>1.01, 3.03</sub> = 0.26, *P* = 0.65 |
| 2v | 2-way ANOVA | Neuron activity: F<sub>1, 3</sub> = 52.93, *P* = 0.005; Training: F<sub>1.27, 3.80</sub> = 3.77, *P* = 0.13; Neuron activity x Training: F<sub>1.62, 4.85</sub> = 3.95, *P* = 0.10 |
| 2w | 2-way ANOVA | Neuron activity: F<sub>1, 3</sub> = 24.70, *P* = 0.02; Training: F<sub>1.06, 3.17</sub> = 4.41, *P* = 0.12; Neuron activity x Training: F<sub>1.02, 3.06</sub> = 3.61, *P* = 0.15 |
| 2x | 2-way ANOVA | Neuron activity: F<sub>1, 3</sub> = 20.44, *P* = 0.02; Training: F<sub>1.40, 4.20</sub> = 1.85, *P* = 0.26; Neuron activity x Training: F<sub>1.38, 4.14</sub> = 0.85, *P* = 0.45 |
| 2y | Planned, Bonferroni corrected, 2-tailed t-tests | Early: t<sub>6</sub> = 5.70, *P* = 0.004, 95% CI 0.13 to 0.48; Middle: t<sub>6</sub> = 6.95, *P* = 0.001, 95% CI 0.19 - 0.54; Overtrain: t<sub>6</sub> = 10.33, *P* = 0.0001, 95% CI 0.38 - 0.72 |
| 2z | 2-tailed Wilcoxon signed rank test | W = -1.00, P > 0.99 |
| 2aa | 2-tailed t-test | t<sub>3</sub> = 2.02, *P* = 0.14, 95% CI -4.52 - 20.15 |
| 2ac | 2-way ANOVA | Event: F<sub>2, 6</sub> = 7.81, *P* = 0.02; Training: F<sub>2, 6</sub> = 4.37, *P* = 0.07; Event x Training: F<sub>1.86, 5.59</sub> = 1.76, *P* = 0.20 |
| 3d | 2-way ANOVA | Training: F<sub>2.22, 31.01</sub> = 27.65, *P* < 0.0001; Genotype: F<sub>1, 14</sub> = 1.27, *P* = 0.28; Training x Genotype: F<sub>4, 56</sub> = 1.24, *P* = 0.30 |
| 3e | 2-way ANOVA | Value x Genotype: F<sub>1, 14</sub> = 14.30, *P* = 0.002; Value: F<sub>1, 14</sub> = 0.49, *P* = 0.49; Genotype: F<sub>1, 14</sub> = 0.00005, *P* = 0.99 |
| 3f | 2-tailed t-test | t14 = 4.01; P = 0.001, 95% CI -0.16 - -0.05 |
| 3j | 2-way ANOVA | Training: F<sub>2.44, 24.43</sub> = 3.23, *P* = 0.048; Genotype: F<sub>1, 10</sub> = 0.31, *P* = 0.59; Training x Genotype: F<sub>8, 80</sub> = 0.68, *P* = 0.71 |
| 3k | 2-way ANOVA | Value x Genotype: F<sub>1, 10</sub> = 3.37, *P* = 0.10; Value: F<sub>1, 10</sub> = 0.56, *P* = 0.47; Genotype: F<sub>1, 10</sub> = 0.30, *P* = 0.59 |
| 3l | 2-tailed t-test | t<sub>10</sub> = 3.07; *P* = 0.01, 95% CI -0.45 - -0.07 |
| 3p | 2-way ANOVA | Training: F<sub>1.59, 28.63</sub> = 68.95, *P* < 0.0001; Genotype: F<sub>1, 18</sub> = 0.74, *P* = 0.40; Training x Genotype: F<sub>4, 72</sub> = 0.44, *P* = 0.78 |
| 3q | 2-way ANOVA | Value x Genotype: F<sub>1, 18</sub> = 4.23, *P* = 0.05; Value: F<sub>1, 18</sub> = 11.93, *P* = 0.003; Genotype: F<sub>1, 18</sub> = 0.64, *P* = 0.44 |
| 3r | 2-tailed t-test | t<sub>18</sub> = 2.15; *P* = 0.045, 95% CI 0.003 - 0.20 |
| 4f | 1-way ANOVA | Training: F<sub>1.39, 4.18</sub> = 3.56, *P* = 0.12 |
| 4g | 2-tailed t-test | t<sub>3</sub> = 1.50, *P* = 0.23, 95% CI -1.86 - 5.16 |
| 4h | One-tailed Bayes factor | BF<sub>10</sub> = 0.19 |
| 4i | 3-way ANOVA | Neuron activity (v. shuffled): F<sub>1, 3</sub> = 1570.57, *P* < 0.001; Training session: F<sub>2, 6</sub> = 5.14, *P* = 0.05; Behavior class: F<sub>3, 9</sub> = 23.33, *P* < 0.001; Neuron activity x Training: F<sub>2, 6</sub> = 3.72, *P* = 0.09; Neuron activity x Behavior: F<sub>3, 9</sub> = 17.65, *P* < 0.001; Training x Behavior: F<sub>6, 18</sub> = 2.00, *P* = 0.12; Neuron activity x Training x Behavior: F<sub>6, 18</sub> = 1.66, *P* = 0.19 |
| 4j | 2-way ANOVA | Neuron activity: F<sub>1, 3</sub> = 65.01, *P* = 0.004; Training: F<sub>1.14, 3.41</sub> = 1.95, *P* = 0.25; Neuron activity x Training: F<sub>1.40, 4.20</sub> = 2.40, *P* = 0.2 |
| 4k | 2-way ANOVA | Neuron activity: F<sub>1, 3</sub> = 42.56, *P* = 0.007; Training: F<sub>1.80, 5.41</sub> = 2.10, *P* = 0.21; Neuron activity x Training: F<sub>1.16, 3.49</sub> = 0.08, *P* = 0.83 |
| 4l | 2-way ANOVA | Neuron activity: F<sub>1, 3</sub> = 70.36, *P* = 0.004; Training: F<sub>1.65, 4.94</sub> = 8.78, *P* = 0.03; Neuron activity x Training: F<sub>1.88, 5.65</sub> = 5.12, *P* = 0.06 |
| 4m | 2-way ANOVA | Neuron activity: F<sub>1, 3</sub> = 82.59, *P* = 0.003; Training: F<sub>1.26, 3.78</sub> = 0.39, *P* = 0.62; Neuron activity x Training: F<sub>1.27, 3.80</sub> = 0.26, *P* = 0.69 |
| 4p | 1-way ANOVA | F<sub>1.42, 4.26</sub> = 1.09, *P* = 0.39 |
| 4q | 1-way ANOVA | Training: F<sub>1.12, 3.35</sub> = 0.67, *P* = 0.48 |
| 5b | 1-way ANOVA | F<sub>1.16, 3.48</sub> = 1.79, *P* = 0.27 |
| 5c | 1-way ANOVA | F<sub>1.05, 3.16</sub> = 1.14, *P* = 0.37 |
| 5g | 2-way ANCOVA | Neuron activity: F<sub>1, 38</sub> = 1.12, *P* = 0.30; Training: F<sub>1, 38</sub> = 7.59, *P* = 0.009; Activity x Training: F<sub>1, 38</sub> = 1.04, *P* = 0.32 |
| 5h | 2-way ANOVA | Neuron activity: F<sub>1, 3</sub> = 3.07, *P* = 0.18; Training: F<sub>1, 3</sub> = 0.68, *P* = 0.47; Activity x Training: F<sub>1, 3</sub> = 0.64, *P* = 0.48 |
| 5i | 3-way ANOVA | Neuron activity (v. shuffled): F<sub>1, 3</sub> = 169.03, *P* < 0.001; Training session: F<sub>2, 6</sub> = 5.55, *P* = 0.04; Behavior class: F<sub>3, 9</sub> = 50.84, *P* < 0.001; Neuron activity x Training: F<sub>2, 6</sub> = 5.27, *P* = 0.05; Neuron activity x Behavior: F<sub>3, 9</sub> = 4.63, *P* = 0.03; Training x Behavior: F<sub>6, 18</sub> = 1.79, *P* = 0.16; Neuron activity x Training x Behavior: F<sub>6, 18</sub> = 0.34, *P* = 0.91 |
| 5j | 2-way ANOVA | Neuron activity: F<sub>1, 3</sub> = 4.80, *P* = 0.12; Training: F<sub>1.50, 4.50</sub> = 1.51, *P* = 0.30; Neuron activity x Training: F<sub>1.36, 4.07</sub> = 1.39, *P* = 0.32 |
| 5k | 2-way ANOVA | Neuron activity: F<sub>1, 3</sub> = 3.74, *P* = 0.15; Training: F<sub>1.15, 4.49</sub> = 0.21, *P* = 0.76; Neuron activity x Training: F<sub>1.62, 4.08</sub> = 0.62, *P* = 0.52 |
| 5l | 2-way ANOVA | Neuron activity: F<sub>1, 3</sub> = 1.10, *P* = 0.37; Training: F<sub>1.45, 4.37</sub> = 1.94, *P* = 0.24; Neuron activity x Training: F<sub>1.34, 4.00</sub> = 3.95, *P* = 0.12 |
| 5m | 2-way ANOVA | Neuron activity: F<sub>1, 3</sub> = 2.53, *P* = 0.21; Training: F<sub>1.01, 3.09</sub> = 1.51, *P* = 0.31; Neuron activity x Training: F<sub>1.10, 3.29</sub> = 1.96, *P* = 0.25 |
| 5n | Planned, Bonferroni corrected, 2-tailed t-tests | Early: t<sub>6</sub> = 2.59, *P* = 0.12, 95% CI -0.07 - 0.55; Middle: t<sub>6</sub> = 3.02, *P* = 0.07, 95% CI -0.02 - 0.59; Overtrain: t<sub>6</sub> = 0.29, *P* > 0.9999, 95% CI -0.28 - 0.34 |
| 5r | 2-way ANCOVA | Neuron activity: F<sub>1, 23</sub> = 0.18, *P* = 0.68; Training: F<sub>1, 23</sub> = 7.13, *P* = 0.01; Activity x Time: F<sub>1, 23</sub> = 0.14, *P* = 0.72 |
| 5s | 2-way ANOVA | Neuron activity: F<sub>1, 3</sub> = 2.01, *P* = 0.25; Training: F<sub>1, 3</sub> = 0.20, *P* = 0.69; Activity x Time: F<sub>1, 3</sub> = 0.31, *P* = 0.62 |
| 5t | 3-way ANOVA | Neuron activity: F<sub>1, 3</sub> = 1296.08, *P* < 0.001; Training session: F<sub>2, 6</sub> = 2.75, *P* = 0.14; Behavior class: F<sub>3, 9</sub> = 32.57, *P* < 0.001; Neuron activity x Training: F<sub>2, 6</sub> = 1.99, *P* = 0.22; Neuron activity x Behavior: F<sub>3, 9</sub> = 11.86, *P* = 0.002; Training x Behavior: F<sub>6, 18</sub> = 2.36, *P* = 0.07; Neuron activity x Training x Behavior: F<sub>6, 18</sub> = 2.91, *P* = 0.04 |
| 5u | 2-way ANOVA | Neuron activity: F<sub>1, 3</sub> = 1.74, *P* = 0.28; Training: F<sub>1.67, 5.01</sub> = 0.31, *P* = 0.71; Neuron activity x Training: F<sub>1.37, 4.12</sub> = 0.33, *P* = 0.67 |
| 5v | 2-way ANOVA | Neuron activity: F<sub>1, 3</sub> = 2.19, *P* = 0.24; Training: F<sub>1.35, 4.04</sub> = 0.04, *P* = 0.90; Neuron activity x Training: F<sub>1.34, 4.04</sub> = 0.11, *P* = 0.82 |
| 5w | 2-way ANOVA | Neuron activity: F<sub>1, 3</sub> = 1.93, *P* = 0.26; Training: F<sub>1.44, 4.33</sub> = 0.21, *P* = 0.75; Neuron activity x Training: F<sub>1.53, 4.56</sub> = 0.20, *P* = 0.77 |
| 5x | 2-way ANOVA | Neuron activity: F<sub>1, 3</sub> = 2.59, *P* = 0.21; Training: F<sub>1.69, 5.07</sub> = 0.98, *P* = 0.42; Neuron activity x Training: F<sub>1.79, 5.36</sub> = 2.35, *P* = 0.19 |
| 5y | Planned, Bonferroni corrected, 2-tailed t-tests | Early: t<sub>6</sub> = 0.59, *P* > 0.999, 95% CI -0.29 to 0.41; Middle: t<sub>6</sub> = 2.58, *P* = 0.13, 95% CI -0.08 - 0.62; Overtrain: t<sub>6</sub> = 2.39, *P* = 0.16, 95% CI -0.10 - 0.60 |
| 5z | 2-tailed t-test | t<sub>3</sub> = 0.37, *P* = 0.74, 95% CI -53.97 - 42.86 |
| 5aa | 2-tailed t-test | t<sub>3</sub> = 0.13, *P* = 0.31, 95% CI -18.88 - 17.44 |
| 5ab | 2-tailed t-test | t<sub>3</sub> = 3.55, *P* = 0.01, 95% CI -7.34 - -1.35 |
| 5ac | Pearson correlation | r<sub>7</sub> = -0.82, *P* = 0.02 |
| 5ad | 2-way ANOVA | Neuron activity: F<sub>1, 3</sub> = 9.03, *P* = 0.05; Training: F<sub>1.12, 3.37</sub> = 0.06, *P* = 0.84; Neuron activity x Training: F<sub>1.06, 3.19</sub> = 0.42, *P* = 0.57 |
| 5ae | 2-way ANOVA | Neuron activity: F<sub>1, 3</sub> = 31.44, *P* = 0.01; Training: F<sub>1.47, 4.40</sub> = 0.42, *P* = 0.62; Neuron activity x Training: F<sub>1.52, 4.55</sub> = 0.28, *P* = 0.71 |
| 6d | 2-way ANOVA | Training: F<sub>2.39, 35.84</sub> = 30.54, *P* < 0.0001; Genotype: F<sub>1, 15</sub> = 0.07, *P* = 0.79; Training x Genotype: F<sub>4, 60</sub> = 0.05, *P* = 0.99 |
| 6e | 2-way ANOVA | Value x Genotype: F<sub>1, 15</sub> = 9.67, *P* = 0.007; Value: F<sub>1, 15</sub> = 0.02, *P* = 0.88; Genotype: F<sub>1, 15</sub> = 0.96, *P* = 0.34 |
| 6f | 2-tailed t-test | t<sub>15</sub> = 3.10; *P* = 0.007, 95% CI 0.057 - 0.31. |
| 6j | 2-way ANOVA | Training: F<sub>1.87, 41.22</sub> = 15.50, *P* < 0.0001; Genotype: F<sub>1, 22</sub> = 0.80, *P* = 0.38; Training x Genotype: F<sub>8, 176</sub> = 0.25, *P* = 0.98 |
| 6k | 2-way ANOVA | Value x Genotype: F<sub>1, 22</sub> = 0.03, *P* = 0.87; Value: F<sub>1, 22</sub> = 0.20, *P* = 0.66; Genotype: F<sub>1, 22</sub> = 1.30, *P* = 0.27 |
| 6l | 2-tailed t-test | t<sub>22</sub> = 0.89; *P* = 0.38, 95% CI -0.26 - 0.10 |
| 6p | 2-way ANOVA | Training: F<sub>1.78, 49.82</sub> = 107.5, *P* < 0.0001; Genotype: F<sub>1, 28</sub> = 1.30, *P* = 0.26; Training x Genotype: F<sub>4, 112</sub> = 5.08, *P* = 0.008 |
| 6q | 2-way ANOVA | Value x Genotype: F<sub>1, 28</sub> = 0.04, *P* = 0.84; Value: F<sub>1, 28</sub> = 47.56, *P* < 0.0001; Genotype: F<sub>1, 28</sub> = 0.50, *P* = 0.49 |
| 6r | 2-tailed t-test | t<sub>28</sub> = 0.22; *P* = 0.83, 95% CI -0.09 to 0.11 |

**Supplemental Table 3:**
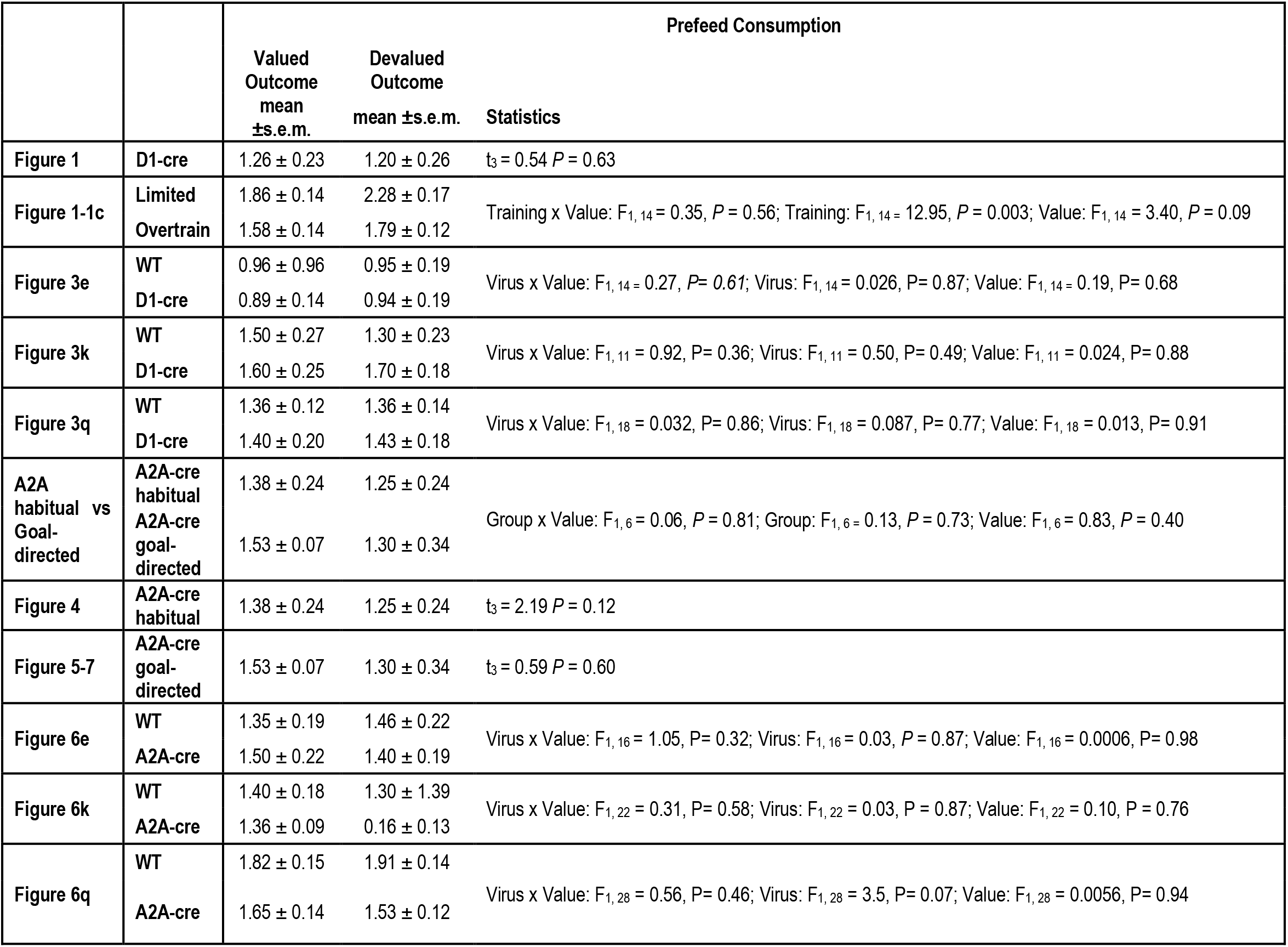
Sensory-specific satiety prefeed consumption. Values reflect average amount consumed in grams ± s.e.m.

|  |  | **Prefeed Consumption** |  |  |
|---|---|---|---|---|
|  |  | **Valued Outcome<br>mean ±s.e.m.** | **Devalued Outcome<br>mean ±s.e.m.** | **Statistics** |
| **Figure 1** | **D1-cre** | 1.26 ± 0.23 | 1.20 ± 0.26 | t<sub>3</sub> = 0.54 *P* = 0.63 |
| **Figure 1-1c** | **Limited** | 1.86 ± 0.14 | 2.28 ± 0.17 | Training x Value: F<sub>1, 14</sub> = 0.35, *P* = 0.56; Training: F<sub>1, 14 =</sub> 12.95, *P* = 0.003; Value: F<sub>1, 14</sub> = 3.40, *P* = 0.09 |
|  | **Overtrain** | 1.58 ± 0.14 | 1.79 ± 0.12 |  |
| **Figure 3e** | **WT** | 0.96 ± 0.96 | 0.95 ± 0.19 | Virus x Value: F<sub>1, 14 =</sub> 0.27, *P= 0.61*; Virus: F<sub>1, 14</sub> = 0.026, P= 0.87; Value: F<sub>1, 14 =</sub> 0.19, P= 0.68 |
|  | **D1-cre** | 0.89 ± 0.14 | 0.94 ± 0.19 |  |
| **Figure 3k** | **WT** | 1.50 ± 0.27 | 1.30 ± 0.23 | Virus x Value: F<sub>1, 11</sub> = 0.92, P= 0.36; Virus: F<sub>1, 11</sub> = 0.50, P= 0.49; Value: F<sub>1, 11</sub> = 0.024, P= 0.88 |
|  | **D1-cre** | 1.60 ± 0.25 | 1.70 ± 0.18 |  |
| **Figure 3q** | **WT** | 1.36 ± 0.12 | 1.36 ± 0.14 | Virus x Value: F<sub>1, 18</sub> = 0.032, P= 0.86; Virus: F<sub>1, 18</sub> = 0.087, P= 0.77; Value: F<sub>1, 18</sub> = 0.013, P= 0.91 |
|  | **D1-cre** | 1.40 ± 0.20 | 1.43 ± 0.18 |  |
| **A2A habitual vs Goal-directed** | **A2A-cre habitual** | 1.38 ± 0.24 | 1.25 ± 0.24 | Group x Value: F<sub>1, 6</sub> = 0.06, *P* = 0.81; Group: F<sub>1, 6 =</sub> 0.13, *P* = 0.73; Value: F<sub>1, 6</sub> = 0.83, *P* = 0.40 |
|  | **A2A-cre goal-directed** | 1.53 ± 0.07 | 1.30 ± 0.34 |  |
| **Figure 4** | **A2A-cre habitual** | 1.38 ± 0.24 | 1.25 ± 0.24 | t<sub>3</sub> = 2.19 *P* = 0.12 |
| **Figure 5-7** | **A2A-cre goal-directed** | 1.53 ± 0.07 | 1.30 ± 0.34 | t<sub>3</sub> = 0.59 *P* = 0.60 |
| **Figure 6e** | **WT** | 1.35 ± 0.19 | 1.46 ± 0.22 | Virus x Value: F<sub>1, 16</sub> = 1.05, P= 0.32; Virus: F<sub>1, 16</sub> = 0.03, *P* = 0.87; Value: F<sub>1, 16</sub> = 0.0006, P= 0.98 |
|  | **A2A-cre** | 1.50 ± 0.22 | 1.40 ± 0.19 |  |
| **Figure 6k** | **WT** | 1.40 ± 0.18 | 1.30 ± 1.39 | Virus x Value: F<sub>1, 22</sub> = 0.31, P= 0.58; Virus: F<sub>1, 22</sub> = 0.03, P = 0.87; Value: F<sub>1, 22</sub> = 0.10, P = 0.76 |
|  | **A2A-cre** | 1.36 ± 0.09 | 0.16 ± 0.13 |  |
| **Figure 6q** | **WT** | 1.82 ± 0.15 | 1.91 ± 0.14 | Virus x Value: F<sub>1, 28</sub> = 0.56, P= 0.46; Virus: F<sub>1, 28</sub> = 3.5, P= 0.07; Value: F<sub>1, 28</sub> = 0.0056, P= 0.94 |
|  | **A2A-cre** | 1.65 ± 0.14 | 1.53 ± 0.12 |  |

**Supplemental Table 4:** Average post-probe-test choice consumption. Values reflect average amount consumed in grams ± s.e.m.

|  |  | **Post-choice Consumption** |  |  |
|---|---|---|---|---|
|  |  | **Valued Outcome mean ±s.e.m.** | **Devalued Outcome mean ±s.e.m.** | **Statistics** |
| **Figure 1** | **D1-cre** | 0.14 ± 0.06 | 0.00 ± 0.02 | t<sub>7</sub> = 2.58 *P* = 0.04 |
| **Figure 1-1c** | **Limited** | 0.32 ± 0.06 | 0.03 ± 0.02 | Training x Value: F<sub>1, 30</sub> = 0.004, *P* = 0.95; Training: F<sub>1, 30</sub> = 0.41, *P* = 0.53; Value: F<sub>1, 30</sub> = 38.49, *P* < 0.001 |
|  | **Overtrain** | 0.29 ± 0.06 | 0.00 ± 0.00 |  |
| **Figure 3e** | **WT** | 0.18 ± 0.04 | 0.06 ± 0.04 | Group x Value: F<sub>1, 30</sub> = 0.038, P= 0.85; Group: F<sub>1, 30</sub> = 0.26, P= 0.62; Value: F<sub>1, 30</sub> = 12.22, P= 0.0015 |
|  | **D1-cre** | 0.16 ± 0.06 | 0.03 ± 0.02 |  |
| **Figure 3k** | **WT** | 0.14 ± 0.05 | 0.07 ± 0.03 | Group x Value: F<sub>1, 24</sub> = 0.98, P= 0.33; Group: F<sub>1, 24</sub> = 0.21, P= 0.65; Value: F<sub>1, 24</sub> = 12.8, P= 0.0015 |
|  | **D1-cre** | 0.15 ± 0.04 | 0.02 ± 0.02 |  |
| **Figure 3q** | **WT** | 0.26 ± 0.04 | 0.05 ± 0.01 | Group x Value: F<sub>1, 38</sub> = 0.047, P= 0.83; Group: F<sub>1, 38</sub> = 0.20, P= 0.66; Value: F<sub>1, 38</sub> = 57.41, P< 0.0001 |
|  | **D1-cre** | 0.28 ± 0.05 | 0.06 ± 0.03 |  |
| **A2A habitual vs Goal-directed** | **A2A-cre habitual** | 0.24 ± 0.10 | 0.01 ± 0.02 | Group x Value: F<sub>1, 14</sub> = 0.11, *P* = 0.75; Group: F<sub>1, 14</sub> = 0.26, *P* = 0.62; Value: F<sub>1, 14</sub> = 12.22, *P < 0.01* |
|  | **A2A-cre goal-directed** | 0.30 ± 0.11 | 0.02 ± 0.01 |  |
| **Figure 4** | **A2A-cre habitual** | 0.24 ± 0.10 | 0.01 ± 0.02 | t<sub>7</sub> = 2.23 P = 0.06 |
| **Figure 5-7** | **A2A-cre goal-directed** | 0.30 ± 0.11 | 0.02 ± 0.01 | t<sub>7</sub> = 2.72 P = 0.030 |
| **Figure 6e** | **WT** | 0.15 ± 0.04 | 0.00 ± 0.01 | Group x Value: F<sub>1, 34</sub> = 0.35, P= 0.56; Group: F<sub>1, 34</sub> = 0.012, P= 0.91; Value: F<sub>1, 34</sub> = 26.2, P< 0.0001 |
|  | **A2A-cre** | 0.13 ± 0.03 | 0.01 ± 0.02 |  |
| **Figure 6k** | **WT** | 0.19 ± 0.04 | 0.02 ± 0.04 | Group x Value: F<sub>1, 46</sub> = 1.9, P= 0.18; F<sub>1, 46</sub> = 0.07, P= 0.79; Value: F<sub>1, 46</sub> = 29.2, P< 0.0001 |
|  | **A2A-cre** | 0.16 ± 0.03 | 0.03 ± 0.02 |  |
| **Figure 6q** | **WT** | 0.29 ± 0.04 | 0.07 ± 0.02 | Group x Value: F<sub>1, 58</sub> = 0.55, P= 0.46; Group: F<sub>1, 58</sub> = 0.039, P= 0.85; Value: F<sub>1, 58</sub> = 72.62; P< 0.0001 |
|  | **A2A-cre** | 0.31 ± 0.04 | 0.04 ± 0.02 |  |

**Supplemental Table 5:**
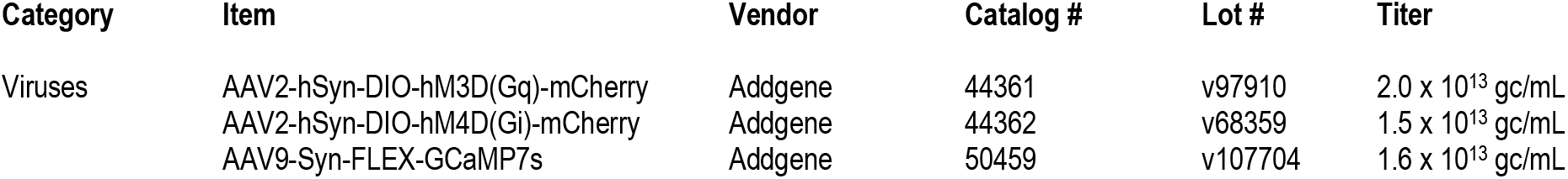
Key reagents information.

